# Multi-substrate specificity shaped the complex evolution of the aminotransferase family across the tree of life

**DOI:** 10.1101/2024.03.19.585368

**Authors:** Kaan Koper, Sang-Woo Han, Ramani Kothadia, Hugh Salamon, Yasuo Yoshikuni, Hiroshi A. Maeda

## Abstract

Aminotransferases (ATs) are an ancient enzyme family that play central roles in core nitrogen metabolism essential to all organisms. However, many of the AT enzyme functions remain poorly defined, limiting our fundamental understanding of the nitrogen metabolic networks that exist in different organisms. Here we traced the deep evolutionary history of the AT family by analyzing AT enzymes from 90 species spanning the tree of life (ToL). We found that each organism has maintained a relatively small and constant number of ATs. Mapping the distribution of ATs across the ToL uncovered that many essential AT reactions are carried out by taxon-specific AT enzymes due to wide-spread non-orthologous gene displacements. This complex evolutionary history explains the difficulty of homology-based AT functional prediction. Biochemical characterizations of diverse aromatic ATs further revealed their broad substrate specificity, unlike other core metabolic enzymes that evolved to catalyze specific reactions today. Interestingly, however, we found that these AT enzymes that diverged over billion years share common signatures of multi-substrate specificity by employing different non-conserved active site residues. These findings illustrate that AT evolution had leveraged their inherent substrate promiscuity to maintain a small yet distinct set of multi-functional AT enzymes in different taxa. This evolutionary history of versatile ATs likely contributed to the establishment of robust and diverse nitrogen metabolic networks that exist throughout the ToL. The study provides a critical foundation to systematically determine diverse AT functions and underlying nitrogen metabolic networks across the ToL.

**Significance Statement:** The ToL-wide analyses of the ubiquitous aminotransferases (AT) family revealed that the broad substrate promiscuity of ATs, which is unusual for core metabolic enzymes, allowed recruitment of distinct, non-orthologous ATs to carry out essential AT reactions in different taxa but without increasing their copy numbers. Some distantly related ATs were also found to exhibit a common signature of multi-substrate specificity by employing different non-conserved active site residues. The versatile evolutionary trajectory of the promiscuous AT enzyme family likely led to biochemical diversity of the robust nitrogen metabolic networks that exist among various extant organisms.

## Introduction

It is hypothesized that the primordial enzymes were highly promiscuous and were able to catalyze a broad spectrum of related reactions. Although such catalysts were likely inefficient, a small set of these multi-functional enzymes could provide the necessary biochemical diversity to sustain ancient metabolism^1–3^. Later, these primordial enzymes divergently evolved through gene duplication and specialization of each paralog to catalyze specific biochemical reactions, so as to increase the metabolic efficiency without needing to increase overall protein expression^4,5^. The resulting rapid expansion of biochemical toolkits likely shaped the core metabolism of the last universal common ancestor (LUCA)^6–9^

The evolution of many ancient protein families involved in the core metabolism has been studied across the tree of life (ToL). The Superfamily Classification of Protein (SCOP) database classifies proteins based on their structural and mechanistic similarities and articulates groups of modern enzyme families that likely originated from the common ancestral enzymes, or founder enzymes^10^. Some superfamilies—e.g., ribosomes^11^, aminoacyl-tRNA synthetases^12–14^, carbonic anhydrases^15^, peptidases^16,17^—are present across all extant organisms and hence were likely present in LUCA^18^. Many of these LUCA enzymes also diverged, functionalized, and specialized to create complex metabolic networks across the ToL. These essential core metabolic enzymes were, in general, inherited vertically to descendants, with some occurrences of lateral gene transfers that replaced the functions of orthologous enzymes that derived from the same founder enzyme^19–21^. Therefore, homology-based prediction of functional annotation is largely effective for most core metabolic enzymes.

Aminotransferases (ATs) are one notable exception and appear to retain high substrate promiscuity, posing significant challenges in the homology-based functional prediction of individual AT proteins or assigning specific AT activities to particular enzymes^26,27^. AT enzymes belong to pyridoxal 5′-phosphate (PLP)-dependent transferases superfamily (**Fig. 1*A*, Fig. S1*A*, Table S1**) and play central roles in core nitrogen metabolism, essential to all organisms^28,27^. For instance, alanine (Ala) ATs from various organisms transaminate Ala to pyruvate for gluconeogenesis, thereby connecting central carbon and nitrogen metabolisms^27,29^. AT-mediated distribution of nitrogen is a key determinant of a wide range of critical biological processes, such as amino acid and protein homeostasis, the synthesis of neurotransmitters, the pathogenesis of infectious diseases, and the recycling of nitrogen^27,30,31^. Thus, deeper understanding of AT functions can lead to better disease diagnostics (e.g., Ala AT as liver damage marker), improved production of nitrogenous compounds (e.g., essential amino acids, alkaloid natural products)^32^, and nitrogen use efficiency in crops^33^. However, the functions of many AT enzymes remain poorly defined, leading to the limited understanding of nitrogen metabolic networks that operate in different organisms.

**Fig. 1.**
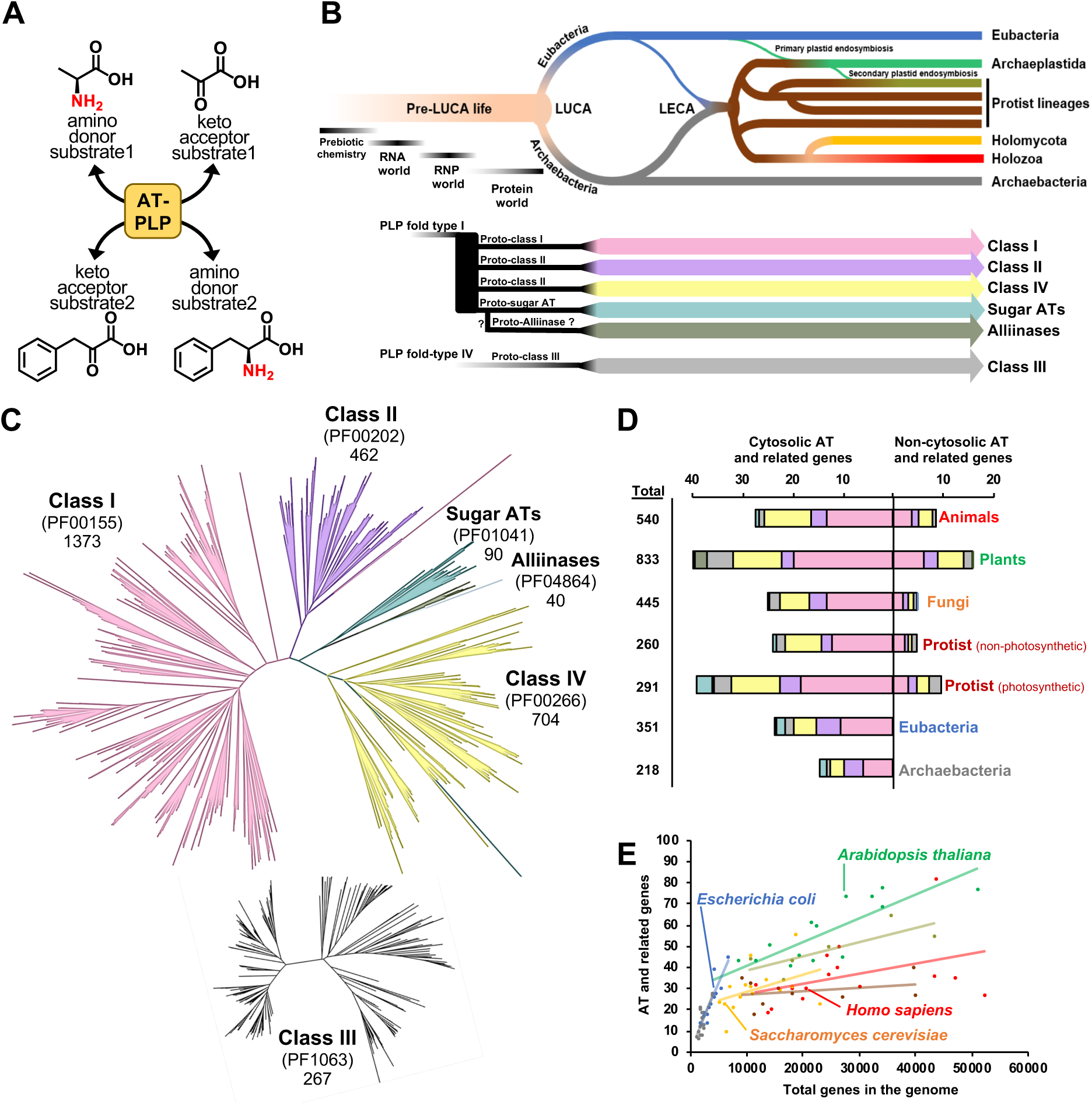
AT enzymes evolved with a limited copy number expansion across the Tree of Life (ToL). ***(A)*** Aminotransferases (ATs) are pyridoxal 5′-phosphate (PLP)-dependent enzymes, which catalyze reversible transamination reactions among at least four substrates. ***(B)*** Evolution of transamination since the origin of life to today. Pre-LUCA life likely had non-enzymatic, RNA, or RNP based transamination. At the interface of RNP and protein worlds, protein-based transaminases appeared in the form of proto-ATs classes. Proto-AT classes were inherited to LUCA and its descendants. ***(C)*** Phylogenetic analysis of AT candidate genes from seven Pfam domains known to contain ATs. Classes I, II and IV, alliinase and sugar ATs are part of PLP-fold type I, while class III is a part of the independently evolved PLP-fold type IV. ***(D)*** Average numbers, Pfam domain composition, and subcellular localization of AT and related genes from animals, plants, fungi, protists, eubacteria, and archaea. ***(E)*** The relationship between the number of AT and related genes versus total number of genes per species. Data corresponding to key model species is labeled and the lines show the overall trend per taxon. Blue, eubacteria; gray, archaea; brown, non-photosynthetic protists; army green, photosynthetic protists; orange, fungi; green, plants; red, animals.

It is estimated that ancestral ATs emerged during the ribonucleoprotein (RNP) world and diversified before LUCA^27^, and, today, are ubiquitously present across the ToL (**Fig. 1*B***). Regardless of their essentiality and billions of years of evolution, ATs still often retain a broad substrate specificity likely due to their inherit mechanistic constraint; ATs catalyze the reversible transamination reaction between amino donors and keto accepters using the ping-pong bi-bi mechanism in a single active site, requiring sufficient versatility to accommodate at least four different substrates. Also, the reactive PLP must be shielded from non-specific substrates to minimize inhibitions (**Fig. 1*A*, Fig. S1*A***)^27,34^. Nonetheless, all extant organisms must maintain robust and complex nitrogen metabolic networks^32,35^, allowing to distribute an essential nutrient, nitrogen, to every corner of organismal metabolism regardless of its availability in different environments. It is unknown, however, how the AT enzymes have overcome these inherent mechanistic constraints associated with the AT reactions and maintained the core nitrogen metabolism.

To address these questions, this study analyzed AT and related enzymes in 90 species across the ToL, which include bacteria, archaea, plants, fungi, animals, and various protist taxa (**Table S2, Fig. S1*B***). The deep evolutionary analyses revealed several interesting paths that the AT enzyme family uniquely adapted. Despite the huge variations in genome sizes and complexities across the ToL, the number of ATs per species remains relatively small and constant. Many essential AT reactions are carried out by taxon-specific AT enzymes due to wide-spread instances of non-orthologous gene displacements^36^ by distantly related AT groups. This finding explained why homology-based predictions of AT functions are difficult and highlighted the existence of yet unknown essential ATs in certain taxa. Some of these distantly related AT enzymes were further found to exhibit conserved multi-substrate specificity, which is accomplished by recruiting different non-conserved active site residues to recognize the same substrate. These results illuminate how AT evolution had overcome their inherent mechanistic constraints to maintain a relatively small number of but a distinct set of AT enzymes in different taxa that likely support robust and diverse nitrogen metabolic networks that exist across the ToL.

## Results

### A small set of ATs maintains essential core nitrogen metabolism across the ToL

In core metabolism, a single or a few enzymes, with high specificity and efficiency, are typically responsible for catalyzing each biochemical step. However, ATs often retain broad substrate specificity^27^, which is typically seen in secondary or specialized metabolic enzymes that underwent significant copy number expansion^37–42^. We therefore investigated how many ATs are present in various extant organisms to maintain core nitrogen metabolic networks by obtaining and analyzing all putative AT sequences from the genomes of 90 organisms across the ToL. We queried the genomes of 15 species from eubacteria^43^, archaea^44^, archaeplastida (plants)^45,46^, holozoa (animals)^47^, holomycota (fungi)^48^, as well as major protist taxa, spanning multiple phyla^49^ (**Fig. S1*B***, **Table S2**), for protein sequences containing conserved protein family (Pfam) domains known for AT enzymes (**Table S1** and **S2**). These species were selected based on the availability of high-quality genomes that represent the diversity of phyla and clades (**Fig. S1*B***, **Table S2**). Inspection of Pfam by HMMER^50^ corrected the Pfam annotations for 93 ATs (**Dataset S1**) and a length criterion (250–1200 amino acids) excluded potential pseudogenes, resulting in 2,938 putative AT genes (**Dataset S2**). Since our ability to detect homology fades over long evolutionary timescales^51,52^, a structure-assisted multiple sequence alignment, MAFFT-DASH^53^, was used to capture the deep evolutionary kinship among distantly related ATs (see Methods).

ATs are found among two independently evolved PLP-dependent enzyme fold-types, I and IV (**Fig. 1*B***^27,28^). The fold-type I includes canonical AT families of class I, II, and IV, as well as smaller families of sugar ATs and alliinases, whereas the fold-type IV only comprises class III ATs. We therefore separately aligned sequences from each Pfam family, having a close phylogenetic proximity^54^, and then merged into two master alignments for independently-evolved fold-type I and IV (**Fig. 1*C***). The phylogeny estimation using an approximately-maximum-likelihood method^55^ captured the monophyletic origins of AT and related enzymes from different Pfam families (**Fig. 1*C***). Overall, class I (PF00155) represented the largest clade having 1,373 out of the 2,938 putative AT enzymes, followed by 704 in class IV (PF00266), 462 in class II (PF00202), and 267 in class III (PF01063, **Fig. 1*C*, Table S2**). Ninety sugar ATs (PF01041), involved in bacterial cell wall biosynthesis^56,57^, were enriched in eubacteria and archaea, while forty alliinases (PF04864) were mainly restricted to plants and protists with secondary plastids, except for two amorphean protists without secondary plastids (*Thecamonas trahens* and *Monosiga brevicollis*, **Table S2**).

The copy number analyses showed that archaea and eubacteria on average have ∼15 and ∼25 AT and related genes, respectively, though a few prokaryotes having parasitic, pathogenic or symbiotic lifestyles^58–60^ had much lower numbers of ATs (**Fig. 1*D***, **Table S2**). In theory, up to 22 AT activities are needed to establish core nitrogen metabolism for synthesis of essential amino acids and nucleic acids^27^. Also, genome reconstruction studies of LUCA^61^ suggested that ∼25 AT and related genes (from classes I, II, III, IV and sugar ATs) were present at the base of the ToL. Therefore, these analyses revealed that the copy numbers of AT enzymes remained constant in prokaryotes.

Eukaryotes had a higher total number of AT and related genes than archaea and eubacteria (**Fig. 1*D*** and **S2*A, B***, **Dataset S3**), with plants and photosynthetic protists having the highest numbers (**Fig. 1*D*** and **S2*A-C*, Dataset S3**). This likely reflects the expansion of metabolic processes (e.g., photorespiration^104^, chlorophyll biosynthesis^62^) necessary to maintain an autotrophic lifestyle. Aside from these photosynthetic organisms, the higher AT number of eukaryotes was attributed to functionally homologous AT isoforms for organelle targeting (**Fig. 1*D***): e.g., three mammalian aspartate (Asp) AT copies with highly redundant functions^63,64^. Additionally, multicellular eukaryotes had tissue specific AT paralogs, such as human AlaAT1 and AlaAT2^65^ and Arabidopsis tryptophan (Trp) ATs, TAA1, TAR1, and TAR2^66,67^. However, the average number of AT candidate genes was only 1.5 and 2.4 times higher in animals and plants than in eubacteria, whereas the average number of genes in the genomes of animals and plants are 8.1 and 6.8 times larger than eubacteria (**Fig. 1*E*** and **S2*C***, **Table S2**). This indicates that, despite the substantial expansion of eukaryotic genome sizes and metabolic networks^68^, AT gene families did not experience substantial copy number expansion even within eukaryotes. This is in contrast to the evolutionary history of specialized metabolic enzymes, many of which are also promiscuous but underwent significant copy number expansion^37,38^. The data revealed that organisms can maintain a robust and complex core nitrogen metabolic network with a set of relatively small numbers of ATs.

### Essential AT reactions are redundantly catalyzed by distinct sets of distant, non-orthologous ATs in different taxa

Ancestral ATs diversified into at least six different classes before LUCA and, today, are ubiquitously present across the ToL (**Fig. 1*B***). However, it is unknown how these founder AT enzymes of LUCA evolved to catalyze essential AT reactions of core nitrogen metabolism, hindering the accurate prediction of AT functions and nitrogen metabolic networks that exist in various organisms. To address this issue, based on the large scale AT phylogeny illustrating an overall relationship of Pfam families (**Fig. 1*C***), we further built forty FastTree trees using multiple starting trees for each AT class to avoid a potential issue of local optima^69^. We then identified 62 distinct AT and related enzyme groups that represent monophyletic clades (bootstrap value ≥ 0.9) (**Fig. S3 to 7, Dataset S3**). The “AT group names” were assigned based on biochemically or genetically validated activities from the Uniprot database query **(Dataset S2)**, though it is important to note that some of them might have been further functionalized in certain taxa. We could not name 15 AT and related groups and hence noted as “uncharacterized (UC)”. Overall, class I contained the largest number of 22 groups with 12 AT, 5 non-AT, and 5 UC enzyme groups (**Fig. S3**) while 6 of 10 groups in class II, 7 of 11 groups in class III, and 2 of 11 groups in class IV as well as 3 groups in sugar AT and alliinases classes were AT groups (**Fig. S4 to S7**). The novel Asp ATs (PF12897, fold-type I)^30,70^, distinct from canonical Asp ATs of class I (PF00155), were found only in two eubacteria and an outgroup of the alliinase class (**Fig. 1*C***). The 17 non-AT groups, which belong to AT classes but likely have non-AT activity (e.g., synthases, lyases, and other transferases), were still included in the analysis, as they improved the phylogenetic relationships of ATs (**Dataset S4**), and some non-AT members can contain ATs, e.g., a Trp AT group found in the alliinases class^71^ (**Fig. S7**). Mapping of AT and related enzymes from four model species— *Escherichia coli*, *Saccharomyces cerevisiae*, *Homo sapiens*, *Arabidopsis thaliana*— suggested complex evolutionary history of AT groups, many of which appear to have been differentially lost in multiple taxa (**Fig. S3 to S7, Dataset S4**).

To further trace the deep evolutionary history of 62 AT and related groups since LUCA, we mapped and clustered different AT and related enzyme groups based on their presence and absence in the 90 species (**Fig. 2, Dataset S5**) and calculated their percent conservation within each taxon (**Fig. 2*A***). The most striking finding was that there was no AT group that is absolutely conserved across the ToL, which was unexpected for core metabolic enzymes catalyzing essential reactions. Only one AT group, class IV Alaglyoxylate ATs (AGTs, **Fig. S6**)—along with non-AT groups for CoA-utilizing synthases (class I, **Fig. S3**) and Cys desulfurases (class IV, **Fig. S6**)—was distributed across almost all taxa though still absent in some species especially in prokaryotes (cluster 1 or CT1 in **Fig. 2*B***). Five AT groups—Asp ATs, Ala ATs, phosphoserine (PSer) ATs, kynurenine AT, and ornithine ATs (CT2, **Fig. 2*B***)—were largely conserved among eukaryotes, but not in prokaryotes. Tyrosine (Tyr) AT and class II AGTs (CT4, **Fig. 2*B***) were present in animals and land plants but largely absent in fungi. Although the outcome might vary with the inclusion of other species, further investigations of selected AT groups (e.g., PSer AT, Tyr AT) using the NCBI Landmark model species database, containing high-quality genome assemblies of 27 species across the ToL (including ten from our 90 species, such as *A. thaliana*, *H. sapiens*, *E. coli*), also yielded consistent results with similar distribution patterns (**Fig. S8**).

**Fig. 2.**
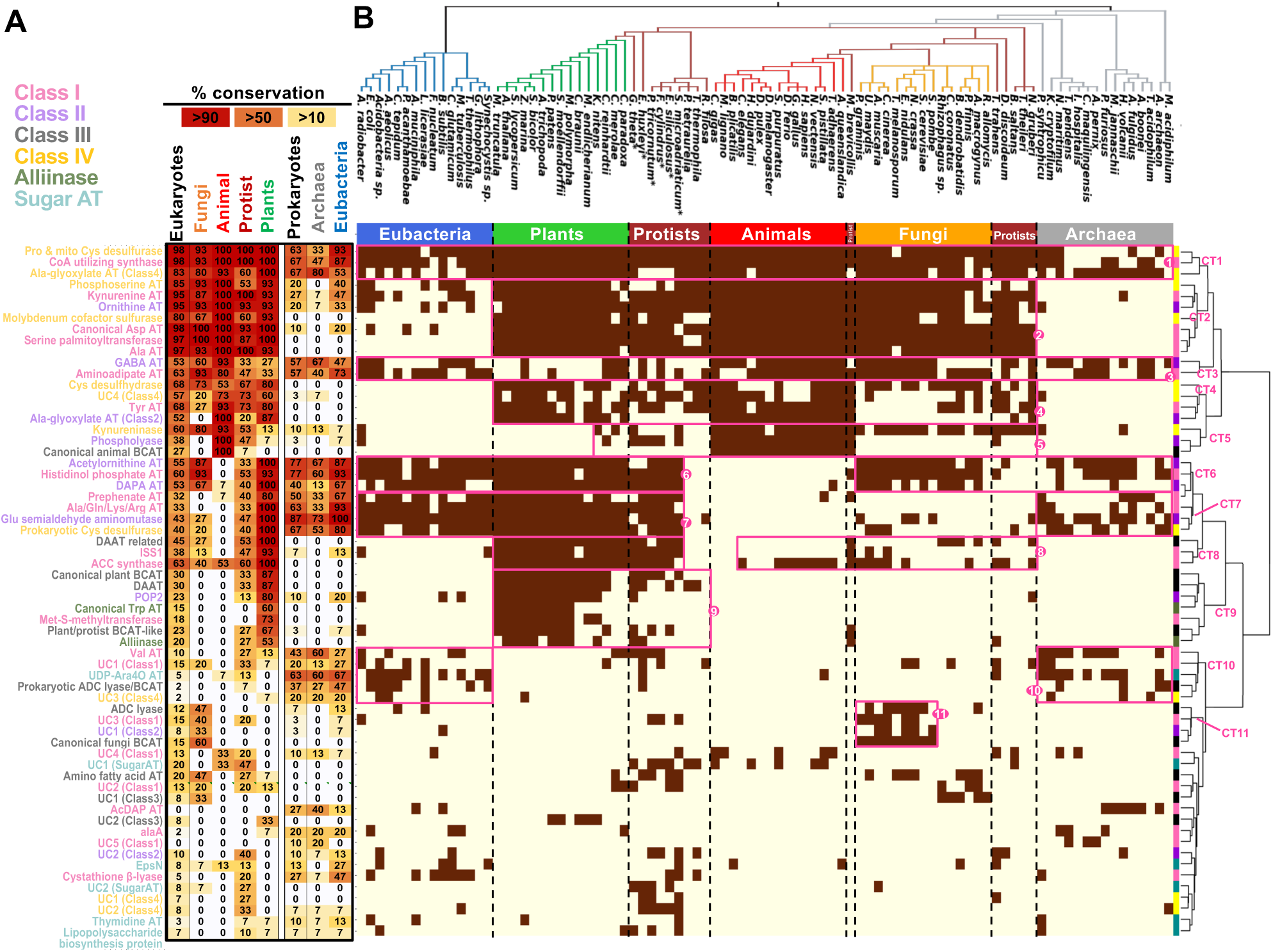
Poor conservation of AT groups across the ToL due to wide-spread replacement of ATs from distantly related non-orthologous AT groups. ***(A)*** Percent conservations (with red to yellow background colors) of 62 AT and related groups for different taxonomic groups and ranks. ***(B)*** AT and related groups were clustered based on the similarity of gene copy numbers within each species. Species were arranged based on the taxonomic relationship at the top: Gray, brown, orange, red, green and blue depict archaea, protists, fungi, animals, plants, and eubacteria, respectively. Species marked with a star (*) contain secondary plastids. Brown filled boxes indicate that certain enzymes (left) are present in the corresponding species (top). Magenta open boxes highlight 11 clusters (CTs) which are labeled at the corresponding branches of the clustering tree.

Histidinol-phosphate (HisP) ATs, 7,8-diaminopelargonic acid (DAPA) ATs, and acetylornithine ATs (CT6, **Fig. 2*B***) were conserved across the ToL except in animals and non-photosynthetic protists, owing to the lack of histidine (His), lysine (Lys), arginine (Arg) biosynthesis in most of these taxa^72–74^. Interestingly, HisP ATs were found in choanoflagellates (e.g., *M. brevicollis*, **Fig. 2*B*, Dataset S5**), suggesting that *de novo* His biosynthesis was likely lost after the divergence of animals and choanoflagellates ∼600 mya^75^. Ala/glutamine (Gln)/Lys/Arg ATs and prephenate ATs, along with glutamate (Glu) semialdehyde aminomutase involved in chlorophyll biosynthesis^76^, were highly conserved in bacteria, plants, and photosynthetic protists (CT8, **Fig. 2*B***). Some AT groups show highly lineage specific distribution, which includes animal- and fungi-specific branched-chain amino acid ATs (BCATs), and land plant-specific Trp ATs that likely functionalized from alliinases (CT5, 11, and 9, respectively, in **Fig. 2*B***). Interestingly, ISS1 (or also known as VAS1, class I, **Fig. S3**)^77^ and D-amino acid AT (DAAT)-related enzyme (class II, **Fig. S4**) groups, whose functions are not fully understood, were highly conserved among all plant species and photosynthetic protists, except diatom *Phaeodactylum tricornutum* (CT9, **Fig. 2*B***).

The above AT mapping across the ToL showed that most AT groups are not ubiquitous, unlike other core enzymes carrying out essential metabolic reactions^11,13,15,68,78–81^. Also, closely related ATs (e.g., class II ATs) did not at all cluster together based on taxonomic distribution (**Fig. 2**). This is likely because AT enzymes from a distant AT group were frequently recruited and replaced existing AT enzymes in certain taxa to carry out essential AT reactions—a phenomenon known as non-orthologous gene displacements^36^. One clear example was the enzymes responsible for Ala AT *activity*, which is essential in all organisms^24^. In eukaryotes, Ala AT activity is provided mainly by class I Ala ATs (97% conserved among eukaryotes, **Fig. 2*A***), and to a lesser extent by the side activities of class II and IV AGTs, and PSer ATs (**Fig. 2*C***, **Fig. S9*A***). In contrast, in some prokaryotes like *E. coli*, Ala AT activity can be redundantly provided by a number of ATs, e.g., avtA, aspC, tyrB, alaA, serC, and ilvE^82^ that are poorly conserved among prokaryotes (**Fig. 2**, **Fig. S9*A***). Since at least one AT enzyme with Ala AT activity was present for all species analyzed, Ala AT *activity* is conserved across the ToL but mediated by taxon-specific AT *enzymes* (**Fig. S9*A***). Additionally, the Tyr AT group (class I) is widespread among eukaryotes but not in many fungi (**Fig. 2**, **Fig. S3**, **Dataset S4**), whose Tyr AT genes were likely lost and replaced with aminoadipate ATs that belong to a distant clade of class I (e.g., yeast Aro8 and Aro9^83^, **Fig. S3**, **Fig. S9*B***). Asp AT activity is provided by canonical Asp AT enzymes (class I) in most organisms and additionally by prephenate AT enzymes in many photosynthetic organisms and some prokaryotes (**Fig. S9*C***). However, a few eubacteria has a non-canonical Asp ATs (PF12897)^30^ and we could not identify any known Asp AT enzymes in many archaea species (**Fig. S9*C***). Similarly, no clear orthologues were found in the land plant genomes for aminoadipate AT enzymes required for Lys metabolism (**Fig. 2*B***, **Fig. S3**). While some of these missing genes could be due to sequencing and/or annotation errors^84,85^, these results overall suggest that there are still unknown ATs responsible for catalyzing these essential AT reactions.

Overall, the global AT mapping across the ToL revealed that essential AT activities were not necessarily carried out by conserved, orthologous enzymes belonging to the same AT class and group in different taxonomic lineages. This likely explains the difficulty in homology-based assignment of AT functions. Instead, we observed many instances of widespread AT enzyme replacement from distantly related, *non-orthologous* AT groups, where ATs from unrelated groups and even different classes were recruited to redundantly or alternatively carry out specific AT reactions. Therefore, in contrast to other core metabolic enzymes, many ATs underwent a complex evolutionary history characterized by the extensive recruitment and replacement of distantly related ATs.

### Distantly related ATs exhibit conserved multi-substrate specificity

The frequent recruitment and displacement of AT enzymes from distant AT groups may be facilitated by the promiscuity and functional redundancy of AT enzymes among different groups. However, AT substrate specificity has not been extensively studied^24^. We therefore experimentally examined the extent of overlap in substrate specificities within the same and among different AT groups. Here we employed aromatic (Aro) AT activity as a testbed, as it utilizes structurally distinct substrates but is widely detected across distantly related ATs within class I (**Table S3**), which include the AT groups of ISS1^86^, canonical Asp AT^87^, aminoadipate AT^83^, HisP AT^88^, Tyr AT^89^, kynurenine AT^90^ and prephenate AT^91^ (**Fig. S3**). As their substrate specificity has not been fully defined, here we expressed, purified, and characterized recombinant enzymes from i) canonical Aro AT groups (as positive controls), ii) non-Aro AT groups with no reported canonical Aro ATs, and iii) non-AT group (i.e., *A. thaliana* C-S lyase, *At* SUR1, as a negative control, **Table S3**). Their substrate specificities were determined by a multi-substrate enzyme assay using 15 keto acceptors with Gln or Glu amino donor depending on the enzymes. AT reactions were initially carried out for an extended reaction time with high substrate concentrations. Then, multiple reaction products were analyzed simultaneously using LC-MS (**Fig. S10*A***) to calculate their percent conversion of the keto acid substrates to the corresponding amino acid products (**Fig. 3*A***).

**Fig. 3.**
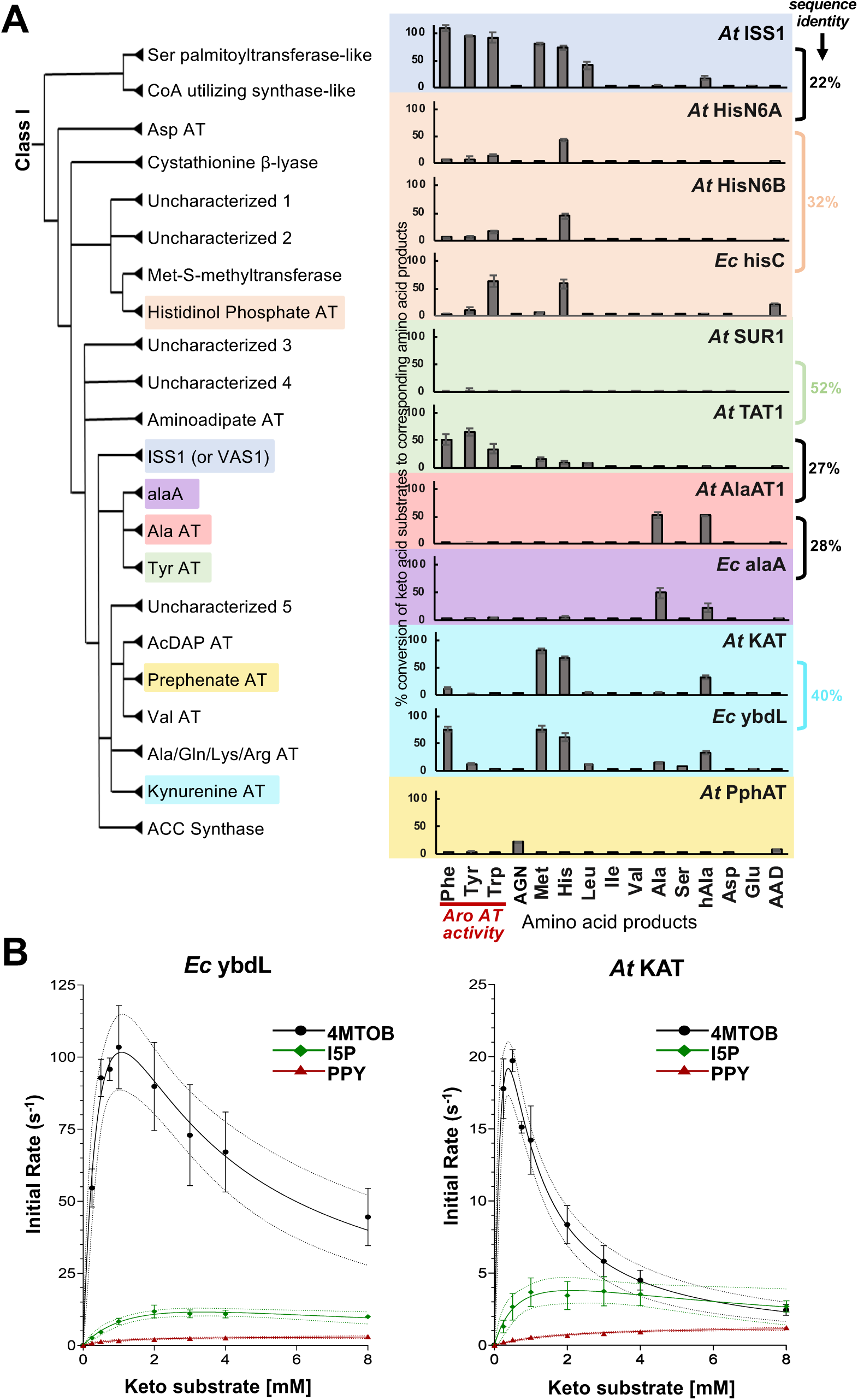
Conserved multi-substrate specificities across distantly related class I ATs. ***(A)*** Percent conversion of keto acids to amino acids by Aro ATs and other ATs. Substrate specificities of these ATs, including Aro AT activity (marked by dark red letters and line), were screened in an assay mixture employing a single amino donor (5 mM Gln or Glu) and 15 acceptors (1 mM each). X-axis shows the amino acid products formed by each enzyme. Amino acid standard curves were used to calculate the concentration of the formed amino acids. Molar ratio of amino acid product to the starting keto acid was used to calculate percent conversion, except for Tyr that was calculated based on the decreases in 4-HPP peak area. Each data point is an average of three separate assays (n=3), except serine which is from a single assay. Error bars show standard deviation among the assays. hAla, homo-Ala; AAD, α-aminoadipate; AGN, arogenate. ***(B)*** Kinetic characterization of *A. thaliana* (*At*) KAT and *E. coli* (*Ec*) ybdL. Enzymatic activity of *Ec* ybdL and *At* KAT were tested with 10 mM Gln as amino donor and varying concentrations of three prominent keto acid substrates: 4-methylthio-2-oxobutanoic acid (4MTOB), imidazol-5-yl pyruvate (I5P), and phenylpyruvate (PPY). Each data point is an average of three separate assays (*n* = 3). Error bars show standard error of the mean (SEM). For 4MTOB and I5P, a modified Michaelis-Menten equation that considers substrate inhibition was fitted using non-linear regression (**Fig. S10**). For PPY, the standard Michaelis-Menten equation was fitted. Percent amino acid sequence identities between key enzymes are given on the right, and complete list is given at **Table S4**.

No Aro AT activity was detected in the Ala AT of *A. thaliana* (*At* AlaAT1) and *E. coli* (*Ec* alaA), nor in *At* SUR1 (**Fig. 3*A***), consistent with prior reports^27,92,93^. In contrast, Aro AT activity was detectable, though at varied degrees, in all of the analyzed non-Aro AT group enzymes (**Fig. 3*A*, Table S3**), which include *At* HisN6 and kynurenine AT (*At* KAT) that are previously not known to have Aro AT activity in plants^94^. This result revealed that Aro AT activity is widespread in many ATs beyond that are currently designated as canonical Aro ATs. To our surprise, HisP ATs from *A. thaliana* and *E. coli* (*At* HisN6A, *At* HisN6B and *Ec* hisC, respectively), having only 32% sequence identity (**Fig. 3*A*, Table S4**), were able to produce the same five amino acids with similar relative substrate preference, e.g., Trp > Tyr > phenylalanine (Phe, **Fig. 3*A***, **Table S5**). Similarly, *At* KAT and *Ec* ybdL, having 40% sequence identity, produced the same eight amino acids with a similar substrate preference, although *Ec* ybdL showed stronger activity with Phe than *At* KAT (**Fig. 3, Table S5**). We also detected the conservation of similar substrate promiscuity *across* different groups. All Aro ATs including ones from ISS1, His AT, Tyr AT, and kynurenine AT groups, except prephenate AT, showed activity with Met, His, Leu and homo-Ala (**Fig. 3*A*, Table S5**). Therefore, although many ATs show very broad substrate promiscuity, we identified the presence of common signatures of multi-substrate specificity among different ATs even after being separated for billions of years of evolution.

For quantitative comparison, we further conducted kinetic characterization of *Ec* ybdL and *At* KAT in their linear ranges of activity (**Fig. S10*B***). Both enzymes have the highest kinetic efficiency with 4-methylthio-2-oxobutanoic acid (4MTOB), followed by imidazol-5-yl pyruvate (I5P) and phenylpyruvate (PPY, keto acids of Met, His, and Phe, respectively, **Fig. 3*B***, **Fig. S10*C*** to ***H***), which largely agreed with the results of the initial substrate specificity screening (**Fig. 3*A***). Interestingly, *Ec* ybdL and *At* KAT showed very similar response curves to these three substrates, except for overall higher turnover rates (*k*_cat_) for *Ec* ybdL than *At* KAT (**Fig. S10*H***). Additionally, both enzymes showed similar substrate inhibition with 4MTOB and I5P, but not with PPY (**Fig. 3*B***), as was also reported for human KAT^95^. Importantly, unlike the animal KATs^90^ and *Ec* ybdL having kynurenine AT activity, *At* KAT was not capable of using kynurenine as an amino donor (**Fig. S11**). This result confirmed the lack of *Ec* ybdL contamination in the recombinant *At* KAT preparation and showed some functional specialization among kynurenine AT orthologs in different organisms.

Taken together, these results uncovered remarkable similarities of multi-substrate specificity and kinetic properties between ATs from the same groups but taxonomically distant species (e.g., *E. coli* vs. *Arabidopsis*). This conserved promiscuity may also represent their ancestral activity that may date back to LUCA. On the other hand, we detected the overlapping multi-substrate specificity among distantly related, non-orthologous AT groups within class I ATs (**Fig. 3*A***), which likely provides functional redundancy and facilitates the AT displacement between distant AT groups (**Fig. 2**).

### The common signature of substrate specificity among distantly related ATs is mediated via distinct yet functionally conserved residues

To elucidate underlying mechanisms of the overlapping substrate specificity, or functional redundancy, among different AT groups (**Fig. 3**), we examined if the presence of certain active site residues or motifs correlates with their substrate specificity. We first obtained experimental or homology-based structures of representative enzymes from each AT group and related (108 enzymes total, **Dataset S6**) and determined the active site residues manually by referring to two crystal structures bound with a hydrophilic Asp (PDB ID: 1ARG^96^) or a hydrophobic Phe (PDB ID: 1W7M)^97^ ligand, respectively. The original multiple sequence alignment was then used to calculate the consensus of the structurally conserved (≥10%) active site residues for each AT group (**Fig. 4*A***).

**Fig. 4.**
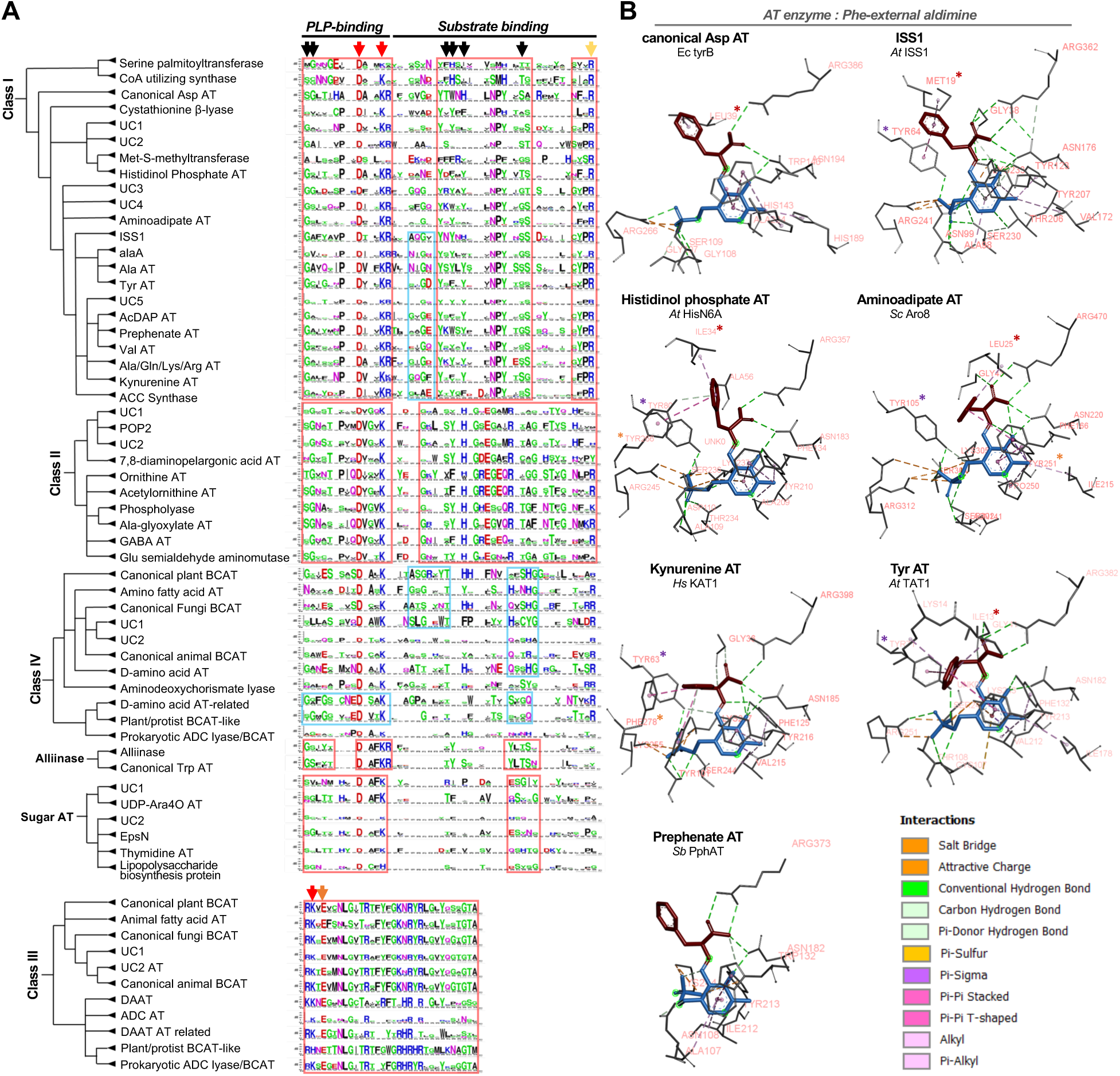
Distinct but functionally conserved residues underlie the conserved multi-substrate specificity among distantly related ATs. ***(A)*** Active site residues are extracted from representative enzymes from each AT group and are used to determine the consensus at those residue positions using the original multiple sequence alignment, which is shown in logo style^132^. Red and orange arrows show previously known well conserved residues for PLP-fold type I ATs, and black arrows show additional residues identified in this study. Red and blue boxes show motifs that are well conserved for an entire or a subset of each AT enzyme class, respectively. ***(B)*** Active site residues that interact with the PLP-Phe aldimine. The portions of the aldimine that corresponds to Phe and PLP are shown in red and blue sticks, respectively. Interacting residues are shown in black sticks and labeled with orange letters. The types of interactions (dashed lines) are colored based on the guide given on the figure. Residues having a conserved function in ligand interactions are marked with red, purple, and orange stars, respectively, as described in the main text.

As expected^27,28^, the Asp and Lys residues involved in cofactor PLP binding (red arrows in **Fig. 4*A***) were highly conserved across all AT groups belonging to PLP-fold type I. The C-terminal Arg residue, which recognizes substrate’s carboxylate (yellow arrows in **Fig. 4*A***), was mostly conserved with some exceptions, such as POP2 and DAPA AT (class II) that use non α-amino acid substrates having an increased molecular distance between the transferable amino and carboxyl groups^98,99^. Besides, several other residues were generally well conserved for most PLP-fold type I groups (black arrows in **Fig. 4*A***), each AT class had certain motifs that are conserved among most or only within a subset of AT groups (orange and blue boxes in **Fig. 4*A***, respectively). Class IV ATs do not share a ubiquitously conserved motif with other class ATs, consistent with its membership in a different fold-type of PLP enzymes ^28,100^ (**Fig. 1**).

Next, we analyzed if distantly-related AT groups with similar multi-substrate specificity have shared residues or motifs among each other, by comparing the active site residues of class I AT enzymes having Aro AT activity: ISS1, Tyr AT, kynurenine AT groups (**Fig. 3**), as well as canonical Asp AT and aminoadipate AT groups^83,86–91^. However, the active site sequences of these five AT groups were not more similar to each other than other class I ATs (**Fig. 4*A***) and did not share common residues or motifs in their active sites. Therefore, we then employed a structure-based comparison to identify ligand interacting residues by docking Phe-external aldimine complex, as a generic aromatic substrate, to the active sites of their homodimeric complex structures (**Fig. 4*B*, Fig. S12**). Interestingly, the unique binding pose of Phe in each AT group requires the involvement of several active site residues that were not necessarily conserved at the level of the primary sequence (**Fig. 4*A*, Table S6**). For example, non-conserved N-terminal aliphatic amino acids interact with the aromatic ring of Phe in Arabidopsis ISS1 (Met 19), Arabidopsis TAT1 (Ile 13), yeast Aro8 (Leu 25), *E. coli* tyrB (Leu 39), and Arabidopsis HisN6A (Ile 34) (red stars on **Fig. 4*B*** and **Fig. S13**). Furthermore, a corresponding residue is absent for human KAT1 and *Streptomyces bingchenggensis* prephenate AT (Sb PphAT), but present for non-Aro ATs such as human and Arabidopsis Ala ATs (Val 64 and Val 82, respectively, **Table S6**). Similarly, the conserved Tyr near the phosphate group of PLP (**Table S6**) interacts with both the aromatic ring of Phe and the phosphate group of PLP in Arabidopsis ISS1 (Tyr 64) and Arabidopsis TAT1 (Tyr 71), but only with PLP in the human KAT1 (Tyr 63), yeast Aro8 (Tyr 105) and Arabidopsis HisN6A (Tyr 80) (purple stars on **Fig. 4*B***). Instead, the Phe 278, Tyr 251, and Tyr 256 residues interact with the substrate Phe in KAT1, Aro8, and HisN6A, respectively (orange stars on **Fig. 4*B***).

We also examined why *Ec* ybdL^101^ and *At* KAT had different specificity towards Phe and kynurenine substrates (**Fig. 3, Fig S10** and **S11**). Docking of these substrates to these two enzymes showed that both enzymes had nearly identical active site residues (**Table S6**); however, there were five residues that are distant to the substrate and located around the active site entrance, which appear to make the *Ec* ybdL active site wider with higher solvent accessibility than *At* KAT, likely aiding the access of bulky aromatic substrates into the active site of *Ec* ybdL (**Fig. S14*A*** and ***B***). Additionally, the N-terminal helix that plays critical role in the substrate binding of ATs by conformational change^102–104^ had more hydrophobic surface in *At* KAT (I74 and V77) than *Ec* ybdL (A25 and Q28) (**Fig. S14*C*** and ***D***), potentially restricting the mobility of the N-terminal helix in *At* KAT.

Overall, these sequence and structural analyses of distantly-related Aro ATs suggest that they achieve similar multi-substrate specificity using different active residues that are not necessarily conserved at primary sequences. Additionally, residues that do not directly participate in substrate recognition appear to contribute to AT substrate specificity by influencing overall active site conformation^104^.

## Discussion

Nitrogen is essential for all life, but its availability significantly varies among different species and in various environments. Therefore, we expect considerable diversity in the number and functionalities of AT enzymes that are at the core of the nitrogen metabolic network. However, despite the ∼200-fold expansion of proteome sizes from the simplest prokaryotes to land plants and metazoans, associated with their metabolic and organismal complexity^68^, we found that the overall copy number of AT genes remains largely unchanged. This is in sharp contrast to many of specialized metabolic enzymes that also exhibit substrate promiscuity and underwent tremendous gene expansion to support chemodiversity and ecological adaption^39–42^. Considering the ancient origins of ATs, many new AT copies must have been generated throughout the evolution, as most eukaryotic lineages underwent whole genome duplication (WGD)^105–109^, and horizontal gene transfer (HGT) events are prevalent among prokaryotes^20,21^. However, the majority of these redundant AT copies were likely lost, resulting in constant AT copy numbers across the ToL. This is likely due to the metabolic cost of maintaining redundant copies involved in core nitrogen metabolism, as compared to relatively low metabolic cost of often conditionally induced specialized metabolism^110^. In support of this hypothesis, carbon-sulfur lyases involved in glucosinolate specialized metabolism^93^, within one branch of Tyr AT group (class I), underwent a rapid copy number expansion in *Arabidopsis thaliana* (**Fig. S3**) and other Brassicaceae species through gene duplication and neofunctionalization^89,94^. Thus, AT enzymes exhibit characteristics of both specialized and core metabolic enzymes; They display promiscuity akin to many specialized metabolic enzymes but didn’t expand their copy numbers similar to other core metabolic enzymes.

Mapping of different AT groups across the ToL showed poor conservation of many AT groups across different organisms (**Fig. 2**). While certain taxa exhibited the loss of specific AT activity and its associated pathways (such as the absence of HisP AT and histidine biosynthesis in animals), many AT reactions remain vital across all life forms. Consequently, the absence of orthologous ATs for these crucial activities in certain taxa suggests widespread occurrences of non-orthologous gene displacements^36^, wherein distantly related ATs have been recruited to catalyze these indispensable reactions. AT enzymes catalyzing essential AT reactions, such as Ala AT and Tyr AT reactions (**Fig. S9**), were replaced by an AT from a distantly related, *non-orthologous* group, such as distinct AT groups within the same AT class (i.e., ancient paralogues) or that belong to different AT classes originated from different founder enzymes (i.e., non-homologues). This is in contrast to other essential core metabolic enzymes that are typically carried out by *orthologous* enzymes^11–17^. This is also different from horizontal or endosymbiotic gene transfer of an ortholog from a different taxa, which sometime displaced the existing ortholog and contributed to the mosaic evolutionary origins of core metabolic pathways^111,112^. Prior studies documented non-orthologous gene displacements for enzymes involved in some essential biochemical pathways, such as cofactor biosynthesis and nucleotide metabolism, which underscored a major hurdle in reconstructing metabolic pathways based on comparative genomic analyses^36,113–115^. This complex evolutionary history of the AT enzyme family, therefore, highlights the challenges associated with homology-based prediction of AT enzyme functions, calling for the needs of additional experimental testing to elucidate and validate AT functions. These findings also suggest that the recruitment of ATs from non-orthologous groups likely led to diverse architecture and functionality of nitrogen metabolic networks that exist in different extant organisms.

What are the underlying mechanisms that potentially contributed to the complex history of AT evolution? The detailed analyses of distantly related class I Aro ATs (**Fig. 3*A*, Table S3-5**) identified that their multi-substrate specificity is highly conserved among ATs from taxonomically distant species, such as kynurenine ATs from *A. thaliana* (plants) and *E. coli* (eubacteria), which also showed striking similarities for their substrate inhibition (**Fig. 3*B*, Figs. S10** and **S11**)^95^. Additionally, a common signature of multi-substrate specificity with His, Met, and Leu, besides aromatic amino acids, was conserved among distantly related Aro ATs that belong to different AT groups (**Fig. 3*A***). These overlapping substrate specificities provide functional redundancy and hence mutation tolerance, allowing a certain AT to be lost or be functionalized without gaining a new enzyme—which will be detrimental to other essential core metabolic enzymes. This can be seen in *E. coli* quadruple mutant that lacks all three major Ala ATs (*alaA, avtA, alaC)* plus *serC* having Ala AT activity (**Fig. S9**) but is still not Ala auxotrophic^92^. Therefore, the extreme functional redundancy, provided by the overlapping multi-substrate specificities across distantly related ATs, likely enabled the widespread AT enzyme replacement without expanding AT gene copy numbers.

Prior studies propose that AT promiscuity likely arose from mechanistic constraints of transamination reactions that require sequential binding and transformation of two substrates, an amino donor and acceptor (**Fig. 1*A*, Fig. S1*A***)^27,116^. These often distinct molecules need to be accommodated in the AT active site, such as through the large-scale rearrangement of the hydrogen bond network or at hydrophilic and hydrophobic pockets^27,116^. Homology-based comparisons of Aro AT active sites failed to identify conserved motifs or residues (**Fig. 4*A*, Table S6**). Instead, we found functional versatility of AT active sites in recruiting new or already existing residues for novel functions (**Fig. 4*B*, Fig. S13**), which can act as a reservoir for the emergence of new AT functions. For example, *Mycobacterium tuberculosis* has an Aro AT (known as Ar AT) within the HisP AT group^88^ (**Fig. S3**), which might have arisen from the positive selection of weak Aro AT activity of an ancestral HisP AT to become a *bona fide* Aro AT enzyme. Additionally, our analysis suggests that residues outside of the active site can induce conformational changes to alter the topography of the active site (**Fig. S14**), which might have also contributed to the emerging specificity of Aro ATs. While more AT enzymes need to be characterized from other AT groups and classes, we hypothesize that the overlapping substrate specificity of distantly related ATs can convergently evolve in distantly related AT groups through recruitment of distinct but functionary conserved residues.

In summary, the multi-functionality of ATs and their functional redundancy increase mutation tolerance of AT genes and allow replacement of distinct ATs that belong to different AT groups or even classes, without necessarily expanding AT gene copy numbers per species. The mixing and matching of AT enzymes from different AT groups and classes, having different side activities and properties, likely contributed to the robust and diverse nitrogen metabolic networks present throughout the ToL^32,35^. This ToL-scale map of the entire AT family, coupled with the use of recently developed mass spectrometry (MS)-based high-throughput AT assays^67,117^, now enables systematic determination of the multi-substrate specificity of various AT enzymes, which remain poorly understood. This will allow further mechanistic studies of the AT functions and multi-substrate specificity and enable artificial intelligence (AI)-based prediction of AT sequence-structure-function relationships. Well-defined AT functionalities from different organisms will allow us to define diverse nitrogen metabolic networks that likely operate in various extant organisms dealing with different nitrogen availabilities and demands. The fundamental understanding of AT functions and nitrogen metabolism will in turn provide rational strategies to develop novel therapeutics against pathogens and metabolic disorders, redesign nitrogen metabolic networks though synthetic biology, and enhance nitrogen use efficiency for sustainable crop production.

## Materials and Methods

### Selection of species

Species used in this study were selected based on the availability of high-quality proteomes (designated as “reference proteome” by uniport, except some plant species queried from the Phytozome). Taxon sampling was performed for each kingdom and protists to include species that would best represent the diversity of the investigated taxonomic group. It is important to note that the selection of the number of species from each taxon is not impartial, as it over-represents eukaryotic lineages, particularly archaeplastida, holozoa and holomycota. However, since species from these groups are also disproportionately the sources for scientific research^118^, we concluded that their over-representation would benefit a broader audience.

### Acquisition of AT sequences

To identify putative AT sequences, we performed Pfam analysis. All protein sequences were downloaded in August 2021 for 90 species from three public database: Phytozome^46^ and Phycocosm^48^ for plants and UniProt^119^ for the others. To avoid redundant protein sequences, we used primary transcripts and non-redundant proteome data for plants and the others, respectively (**Dataset S7**). The proteins shorter than 250 amino acids were excluded owing to the potential for non-functional genes. The sequences after length filtration proceed to Pfam annotation by HMMER v3.3.1 (hmmscan with a default setting; E-value cutoff=0.01)^120^ using Pfam profile hidden Markov models obtained from Pfam v35.0^50^. In the case of multiple hits for the same region in the protein sequence, we chose the Pfam with lower E-value. Then, we obtained 2,938 putative AT sequences (**Dataset S2**) by searching Pfam IDs in which proteins display AT activity: PF00155, PF00202, PF00266, PF04864, PF01041, PF12897 (PLP fold type I), and PF01063 (PLP fold type For the extremely long proteins (>1200 amino acids), we extracted the only AT Pfam domain region. In the case that multiple AT Pfam domains exist, the longer AT region was selected.

### Multiple sequence alignment of AT sequences and the phylogenetic tree construction

For structure-guided multiple sequence alignment, amino acid sequences were aligned by a MAFFT-DASH^53^ using a BLOSUM62 scoring matrix (gap opening penalty=1.53). We assumed closer associations among the proteins belonging to the same Pfam. Thus, all putative AT sequences were divided into seven groups according to Pfam IDs, followed by multiple sequence alignment for each group with iterative refinement methods (L-INS-i for PF01041, PF04864, and PF12897; FFT-INS-i for PF00155, PF00202, PF0266, and PF01063). Then, considering the evolutionary divergence of PLP fold type I, we merged the alignments for six Pfam (i.e., PF00155, PF00202, PF00266, PF04864, PF01041, and PF12897) into a single multiple sequence alignment for PLP fold type I with progressive method (FFT-NS-2). The alignment for PF01063 was regarded as the one for PLP fold type IV. For further analysis, we deleted DASH sequences from the alignments to avoid misinterpretation.

To construct phylogenetic trees for large alignments, we employed an approximately maximum-likelihood method. The alignments for PLP fold type I and IV were analyzed by FastTree v2.1.11^55^ with LG+CAT model and the option “-mlacc 2 - slownni -spr 4 -pseudo”. To avoid a potential issue of a local optima described in the previous study^69^, we generated 40 FastTree trees using 20 random and 20 parsimony starting trees prepared by RAxML-NG (v1.2.0) with a JTT+G model, and then selected the most likelihood trees. For the PLP-fold type I, we additionally built 40 FastTree trees for each class (e.g., class I ATs, Alliinases, Sugar ATs) and then selected the most likelihood trees. To evaluate the reliability of each split in the tree, we used local support values^121^ computed in FastTree from 1,000 resamples. We analyzed the tree and modified the tree topology using an ETE 3 Toolkit^122^ in a Python environment (v3.6.13). To identify the highly supported phylogenetic branches, the nodes with low supporting values (< 90%) were deleted and then integrated into parental nodes. The condensed phylogenetic trees were manually drawn using Microsoft PowerPoint with local support values as branch support.

The functions of AT and related groups were inferred from the publicly-available experimental data. First, we picked four species whose enzymes have been well-defined (i.e., *Homo sapiens*, *Arabidopsis thaliana*, *Saccharomyces cerevisiae*, and *Escherichia coli*), and then inquired about the functions of AT candidates from literature search. Also, we collected additional 68 enzymes whose experimental data are available in the UniProtKB^119^, the BRENDA^123^, or the SABIO-RK^124^ databases. This information was utilized to annotate names for the AT and related groups. We labeled the group with “uncharacterized (UC)” when any group member was unavailable in the collection. Only enzyme clades that have more than five sequences from at least four species or three kingdoms with a bootstrap value higher than 90% were assigned AT groups names. Sequences that did not belong to any groups were indicated as “unassigned (UA)”.

### Mapping of AT groups on the taxonomic tree

We computed how many AT-coding genes each species carries for 62 AT and related groups. This dataset was used for two different analyses to examine taxonomic conservation and distribution. For the taxonomic conservation, the sum of the gene counts was calculated in each kingdom for each AT and related group. The values were used to generate a heatmap using Microsoft Excel.

To investigate the taxonomic distribution, we first transformed the gene counts into the possession (i.e., 0: absence, >1: presence) of individual AT and related groups. A heatmap was generated with columns being clustered (distance metric: ‘euclidean’, linkage method: ‘ward’) by the seaborn package v0.11.2^125^ in a Python environment, whereas rows were arranged phylogenetically by referring to the previous studies^43–45,47–49^ to be able to trace the evolutionary history of individual ATs.

To separately test the taxonomic distribution of selected AT groups, we searched the presence and absence of sequences from the alaA, Ala AT, Tyr AT, AcOAT, AGT1 (class IV), and PSer AT groups within the NCBI Landmark model species database, which includes non-redundant proteomes from the most recent representative assembly of 27 genomes spanning a wide taxonomic range (**Fig. S8**). The BlastP search was conducted to identify sequences that potentially belong to these AT groups using the query sequences marked in **Fig. S8**. To confirm the assignments of each sequence to corresponding AT group, three separate phylogenetic trees were generated for ATs in class I (alaA, Ala AT, Tyr AT), class II (AcOAT), and class IV (AGT1, PSerAT) using MEGA7 and the Maximum Likelihood method based on the JTT matrix-based model. AGT1 was initially absent in the genome assembly of *Caenorhabditis elegans* from the landmark database, despite other animals process AGT1 (**Fig. S8**); however, an AGT1 sequence (NP_495885) was later found in a different genome assembly of *C. elegans* (NP_495885) and was hence included in **Fig. S8**.

### Analysis of subcellular localization of AT and related enzymes

All AT candidates determined by Pfam analysis proceeded to the prediction of their subcellular localization. The presence of N-terminal pre-sequences for subcellular localization was analyzed using TargetP-2.0 server^126^. We focused on the mitochondrial and chloroplast transit peptide, and ignored the presence of signal peptides for secretory pathways. Based on this prediction, the AT candidates were classified into 3 groups according to subcellular location: mitochondria, chloroplast, and others.

### Protein expression and purification

AT enzymes were expressed and purified as previously described quarried^117^. Briefly, expression constructs for Arabidopsis ISS1 (AT1G80360), HisN6A (AT5G10330), HisN6B (AT1G71920), SUR1 (AT4G23600), TAT1 (AT5G53970), AlaAT1 (AT1G17290), KAT (AT1G77670) and PphAT (AT2G22250), and *E. coli* hisC (P06986), alaA (P0A960) and ybdL (P77806) were cloned to a modified pET28a vector, transformed to chemical competent KRX (DE3) *E. coli* cells (Promega, Madison, WI) and selected on LB agar with 50 µg/ml spectinomycin. Colonies for each construct were picked, inoculated in 10 mL LB medium with 50 µg/ml spectinomycin, and incubated overnight at 37°C, 200 rpm. Ten milliliters of the culture was transferred to 500 mL of fresh terrific broth medium (Teknova, Hollister, CA) and grown at 37°C, 200 rpm until OD_600_ reached ∼0.55, when the temperature was dropped to 22°C and isopropyl β-d-1-thiogalactopyranoside (IPTG) and L-rhamnose (Promega, Madison, WI) was added at 0.2 mM and 0.1% (w/v) final concentration, respectively. After overnight incubation at 22°C, 200 rpm, cells were harvested by centrifugation at 6000 RPM for 20 minutes at 4°C. The pellet was either stored at −80°C for later use, or resuspended in 10 mL of lysis buffer containing 50 mM sodium phosphate (pH 8.0), 300 mM NaCl, 25 µM PL, and 0.25 mg/mL lysozyme (Sigma-Aldrich, St. Louis, MO). After disrupting cells by three freeze-thaw cycles and sonication, the soluble extract was obtained by centrifugation at 4°C, 18,000 x *g* for 30 min. His-tagged recombinant proteins were purified using nickel conjugated HisTrap^TM^ Fastflow crude column (Cytiva, Marlborough, MA) with ÄKTA™ pure chromatography system (GE Healthcare Life Sciences, MA). Purified proteins were desalted with Sephadex^TM^ G-50 superfine resin (Cytiva, Marlborough, MA) into 100 mM phosphate buffer (pH 8.0) containing 0.025 mM PLP and 10% (v/v) glycerol. Recombinant proteins were separated by SDS-PAGE, and the gels were Coomassie stained and imaged (ChemiDoc, Bio-Rad, Hercules, CA), and their purity was assessed by ImageJ. The same procedure was repeated for the KRX cells without an expression plasmid to obtain a purification extract that can be used as a background control.

### Multi-substrate specificity assay of AT enzymes

Multi-substrate specificity screenings of ATs were conducted as previously described^117^. Briefly, transamination capacity of Arabidopsis ISS1, HisN6A, HisN6B, SUR1, TAT1, AlaAT1, KAT and PphAT, and *E. coli* hisC, alaA and ybdL and purification extract from the KRX cells without an expression plasmid were tested in a single assay mixture containing 14 keto acids (pyruvate, α-ketoglutarate, 4-methylthio-2-oxobutyrate, phenylpyruvate, imidazol-5-yl-pyruvate, 4-hydroxyphenylpyruvate, indole-3-pyruvate, oxaloacetate, hydroxypyruvate, 3-methyl-2-oxopentanoate, 4-methyl-2-oxopentanoate, 3-methyl-2-oxobutanoate, prephenate, α-ketobutyrate, oxoadipate) together with 5 mM amino donor (Gln for ISS1, KAT and ybdL, Glu for the rest), 100 mM phosphate buffer pH 8.0, 0.025 mM PLP, 1 mM ethylenediaminetetraacetic acid (EDTA), 20 ng/µL AT enzyme. Two separate master mixes (with Gln or Glu) without the enzymes were prepared, α-ketoglutarate was omitted from the master mix with Glu to prevent non-productive transamination. Master mixes were preheated to 37°C for 3 minutes and the reactions were initiated by adding the master mix to tubes with each AT enzyme. The reaction mixtures were incubated at 37°C for 90 min and terminated by adding 400 µL LC-MS grade methanol (final 80% (v/v) methanol). Samples were cooled down on ice, and the soluble phase was collected by centrifugation at 15,000 x *g* for 10 min. Transamination products were analyzed using Vanquish UHPLC system coupled with the Q Exactive Quadrupole-Orbitrap MS (Thermo Fisher Scientific, Waltham, MA) with negative ionization using settings as previously described^117^. Authentic standards of amino acids were used to convert peak areas to concentration. Percent conversion values were determined by calculating the molar ratio of amino acid product to the starting keto acid. The reactions conducted with KRX extract were used as the background controls.

### Kynurenine AT assay

The kynurenine aminotransferase reactions were tested in a reaction mixture with final concentration of 100 mM phosphate buffer pH 8.0, 0.025 mM PLP, 1 mM EDTA, 20 ng/µL AT enzyme, 1 mM 4-methylthio-2-oxobutyrate and 0.5 mM ^14^N-kynurenine or amino-^15^N-Gln or both ^14^N-kynurenine and amino-^15^N-Gln in 100 µL final volume. A master mix without the enzyme was prepared and then preheated to 37°C. The reactions were initiated by adding the master mix to tubes with AT enzyme, incubated at 37°C for 60 min and terminated by adding 400 µL LC-MS grade methanol (final 80% (v/v) methanol). Samples were cooled down on ice, and the soluble phase was collected by centrifugation at 15,000 x *g* for 10 minutes. Formation of ^14^N-Met (m/z: 148.0416 - 148.0460) or ^15^N-Met (149.0386 - 148.0430) was analyzed using the Vanquish UHPLC system coupled with the Q Exactive quadrupole-Orbitrap MS (Thermo Fisher Scientific, Waltham, MA) with negative ionization using settings as previously described^117^ to determine the utilization of kynurenine or Gln. Authentic standards for Met was used to convert peak areas to concentration.

### AT enzyme kinetic assays

Kinetic AT assays are conducted as previously described^117^. Briefly, the transamination of 4-methylthio-2-oxobutyrate, imidazol-5-yl-pyruvate, and phenylpyruvate to Met, His, and Phe, respectively, by *At* KAT and *Ec* ybdL were tested in a reaction mixture with final concentration of 100 mM phosphate buffer pH 8.0, 10 mM Gln, 0.025 mM PLP, 1 mM EDTA, 0 to 8 mM of the keto acceptor given on the figure and 4 ng/µL KAT or 1 ng/µL ybdL, in 100 µL final volume. Two separate master mixes were prepared for each enzyme having all the components except the keto acids and preheated to 37°C for 3 minutes. Reactions were initiated by adding each master mix to reaction tubes with the keto acids to give desired concentrations (0 to 8 mM). The reaction mixtures were incubated at 37°C for 10 min and terminated by adding 400 µL LC-MS grade methanol (final 80% (v/v) methanol). Samples were cooled down on ice, and the soluble phase was collected by centrifugation at 15,000 x *g* for 10 minutes. Formation of Met (m/z: 148.0416 - 148.0460), His (154.0599 - 154.0645) and Phe (164.0692 - 164.0742) was analyzed using Vanquish UHPLC system coupled with the Q Exactive quadrupole-Orbitrap MS (Thermo Fisher Scientific, Waltham, MA) with negative ionization using settings as previously described^117^. Authentic standards for Met, His and Phe were used to convert peak areas to concentration. The reaction with 0 mM keto acid was used as the background control. Kinetic parameters were calculated with GraphPad. All enzyme assays were performed under the condition where the product formation increased proportionally to the enzyme concentration and the reaction time.

### Determination of the variance of active site residues

We initially picked 111 model proteins over 62 AT-like groups (1–4 enzymes from each group) for structural analysis. The crystal structures of 57 proteins were obtained from the RCSB Protein Data Bank (PDB)^127^. For the other proteins, homodimeric structure models were generated by AI-based structure prediction using AlphaFold v2.1.0^128,129^. The predicted structures were manually checked if the dimeric interfaces and the active site formed properly. In the case of a distorted structure or collapsed active site, we constructed another structure model by conventional homology modeling using SWISS-MODEL^130^. When the models were still poor, they were excluded from further analyses. Finally, we obtained the structures for 107 proteins over 56 AT groups (**Dataset S6**). Then, the structures were aligned with each other using the SALIGN module of MODELLER v10.2^131^ and then used for active site analysis. Active site residues were determined manually referring to the protein-ligand complex structures determined previously with Asp and Phe (PDB ID: 1ARG and 1W7M, respectively)^97,103^. The corresponding positions of the active site residues were examined in multiple sequence alignments and minor residues (i.e., <10% conservation among model proteins) were removed. As a result, we obtained 50 and 32 residues for PLP fold type I and IV, respectively. Next, the putative active site residues were extracted from multiple sequence alignments for 2,938 ATs. The lists of active site residues were divided according to AT-like groups, followed by generation of consensus sequences using WebLogo^132^.

### Molecular docking studies

The docking simulations of external aldimine intermediates were performed against 9 ATs (**Dataset S8**) using AutoDock v4.2.6^133^. The identical structures obtained for the active site analysis were used for docking simulation except for the prephenate AT from *Streptomyces bingchenggensis* (Sb Prephenate AT). Sb Prephenate AT was not oriented from the 90 model species, thus its homodimeric structure was additionally constructed by AlphaFold prediction. We preprocessed the crystal structures from the RCSB PDB by deleting all small molecules such as water molecules, PLP cofactors and other ligands. Then, the nine homodimeric structures were aligned altogether using the MatchMaker tool (Needleman-Wunsch algorithm, BLOSUM62 matrix, gap extension penalty=1) in the Chimera v1.16^134,135^ and saved as PDB files. A grid box for docking simulation was defined according to the active site residues with the spacing of 0.375 Å and the x/y/z grid points numbers of 50/50/50. The external aldimine intermediate ligands were prepared as MDL Molfile using the Chemdraw v20.0 (PerkinElmer Informatics, Waltham, MA, USA) and then converted to PDB using the Discovery Studio Visualizer 2021 (Dassault Systemes BIOVIA, San Diego, CA, USA). The C4′-Nα bond of the external aldimine and the catalytic lysine of the protein were manually assigned to a non-rotatable bond and a flexible residue, respectively. Lamarckian genetic algorithm was used for 100 docking runs with a population size of 300. Then, the protein-ligand interactions were analyzed using the Discovery Studio Visualizer.

## Acknowledgements

We thank Dr. Prashant Sharma for helping us with the selection of animal species to be used in the ToL analysis. We thank Dr. Anne Pringle and Dr. Yen Wen Wang for helping us with the selection of fungi species to be used in the ToL analysis. Dr. Garret Suen for helping us with the selection of eubacteria species to be used in the ToL analysis. We thank Dr. Karthik Anantharaman for helping us with the selection of archaea species to be used in the ToL analysis. This work was supported by the US Department of Energy (DOE), Office of Science, Office of Biological and Environmental Research, Genomic Science Program (grant no. DE-SC0020390 and DE-AC02-05CH11231), the Joint Genome Institute award no. CSP-503757, as well as the U.S. National Science Foundation award PGRP-IOS-2312181. The work (10.46936/10.25585/60001160) conducted by the U.S. Department of Energy Joint Genome Institute (https://ror.org/04xm1d337), a DOE Office of Science User Facility, is supported by the Office of Science of the U.S. Department of Energy operated under Contract No. DE-AC02-05CH11231.

## Author Contributions

Conceptualization: K.K., S.H.W., Y.Y., H.A.M.

Investigation: K.K., S.H.W., R.K., H.S.

Analysis: K.K., S.H.W., H.A.M.

Visualization: K.K., S.H.W., H.A.M.

Project administration: Y.Y., H.A.M.

Writing – original draft: K.K., S.H.W., H.A.M.

Writing – review & editing: K.K., S.H.W., Y.Y., H.A.M.

## Classification

BIOLOGICAL SCIENCES, Biochemistry; Evolution.

## SUPPLEMENTAL FIGURES AND TABLES

**Fig. S1.**
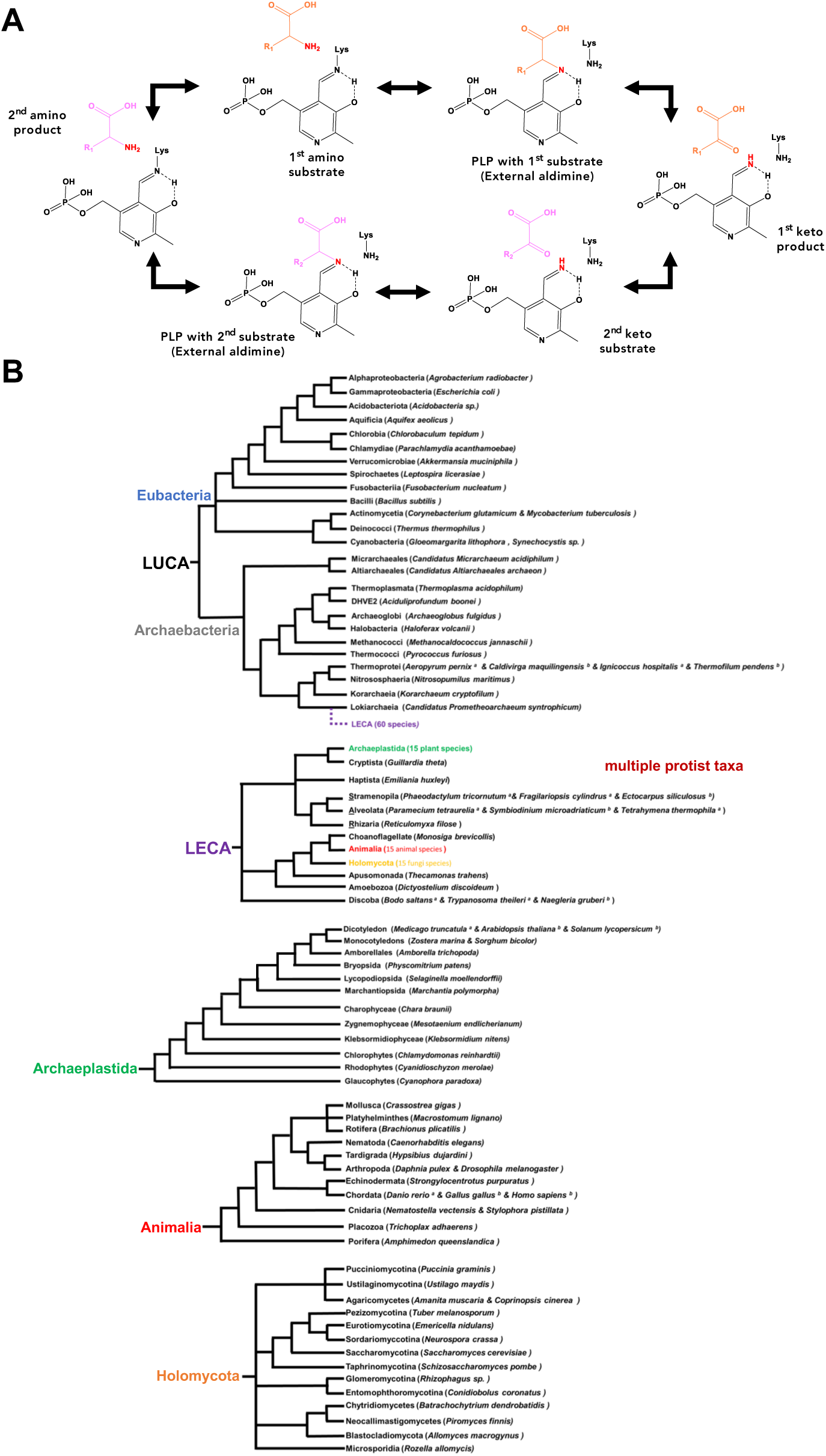
***(A)*** Ping-pong bi-bi reaction mechanisms of ATs. In the 1^st^ half reaction, an amino acid substrate (R_1_) displaces Schiff-base lysine of the PLP-enzyme complex which leads to the transfer of the amino group onto pyridoxamine phosphate (PMP) and the 1^st^ keto product is released. In the 2^nd^ half reaction, a keto acid substrate (R_2_) attacks the nitrogen atom on PMP which forms the 2^nd^ amino acid product and regenerates enzyme-PLP complex. Note that only key intermediates are shown for representative purposes. ***(B)*** Taxonomic relationship of eubacteria ^1^, archaebacteria^2^, protists^3^, plants^4^, animals^5^ and fungi^6^ used in the study. LUCA, last universal common ancestor; LECA, last eukaryotic common ancestor. Taxonomic tree only depicts the topology with arbitrary branch lengths. Blue, eubacteria; gray, archaebacteria; brown, protists; orange, fungi; green, plants; red, animals.

**Fig. S2.**
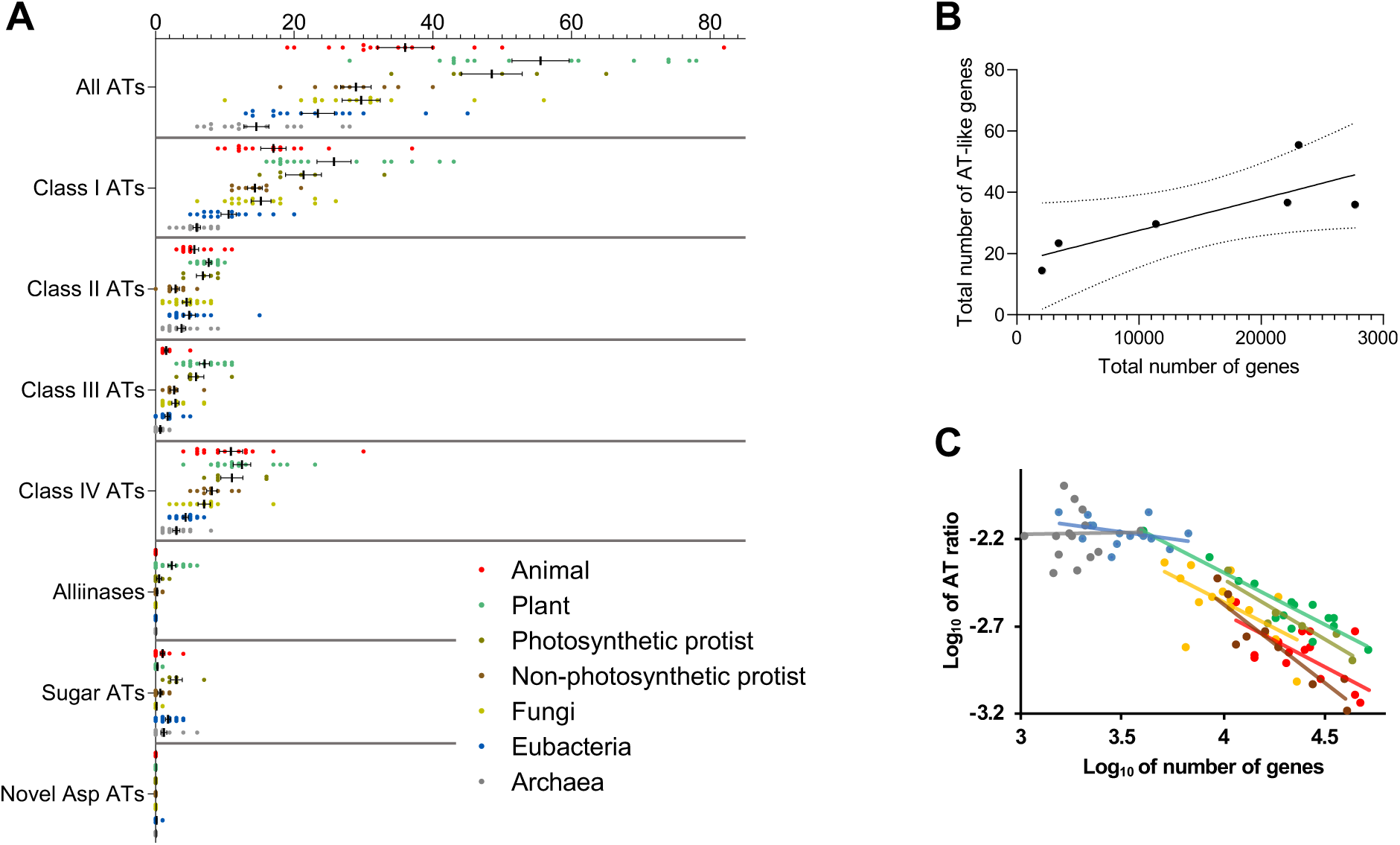
AT and related genes across different taxonomic groups. ***(A***) Distribution and number of AT genes for all ATs or ATs from each class across animals, plant, photosynthetic protists, non-photosynthetic protists, fungi, eubacteria and archaea. For each dataset means and SEM are shown. Statistical significance between the groups is shown in **Dataset S3**. ***(B)*** Comparison of total number of AT genes and the total number of genes across the studied taxonomic groups. A positive correlation was determined between AT gene number and total gene number (Pearson correlation = 0.8), but it was not statistically significant. ***(C)*** The Log ratio of AT and related genes to total number of genes.

**Fig. S3.**
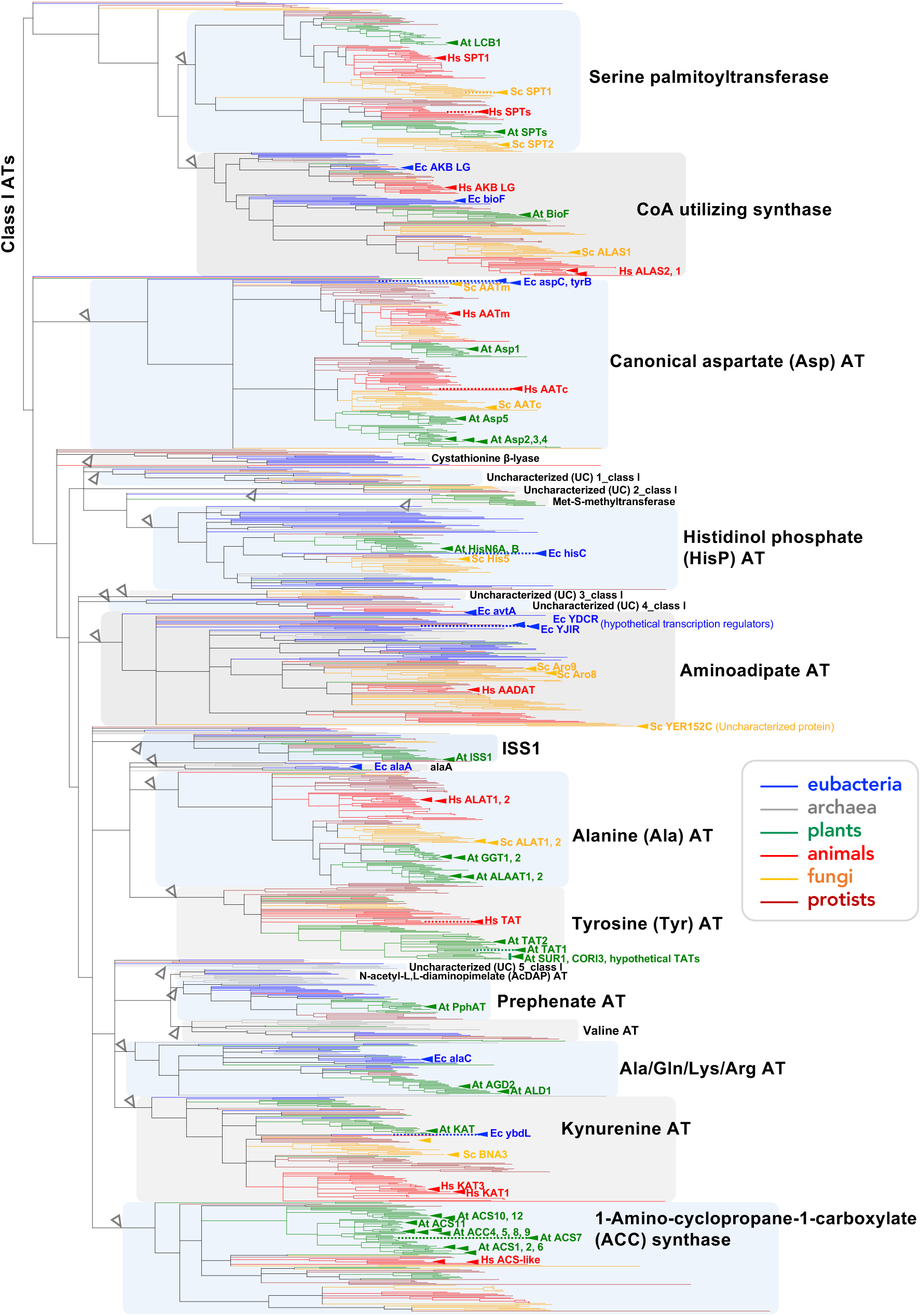
Phylogeny of **Class I** AT and related enzymes. Branch colors of blue, gray, green, red, orange, and maroon represent enzymes from eubacteria, archaea, plants, animals, fungi, and protists, receptively. Monophyletic AT groups are named according to the activities reported for the labeled enzymes from the four model organisms—*Arabidopsis thaliana* (green), *Homo sapiens* (red), *Saccharomyces cerevisiae* (orange), and *Escherichia coli* (blue). Groups without any representative enzymes are labeled as “uncharacterized (UC)”. Open arrow heads depict the root of each AT and related group. The corresponding full phylogenetic tree with all sequence IDs is provided as **Dataset S4**.

**Fig. S4.**
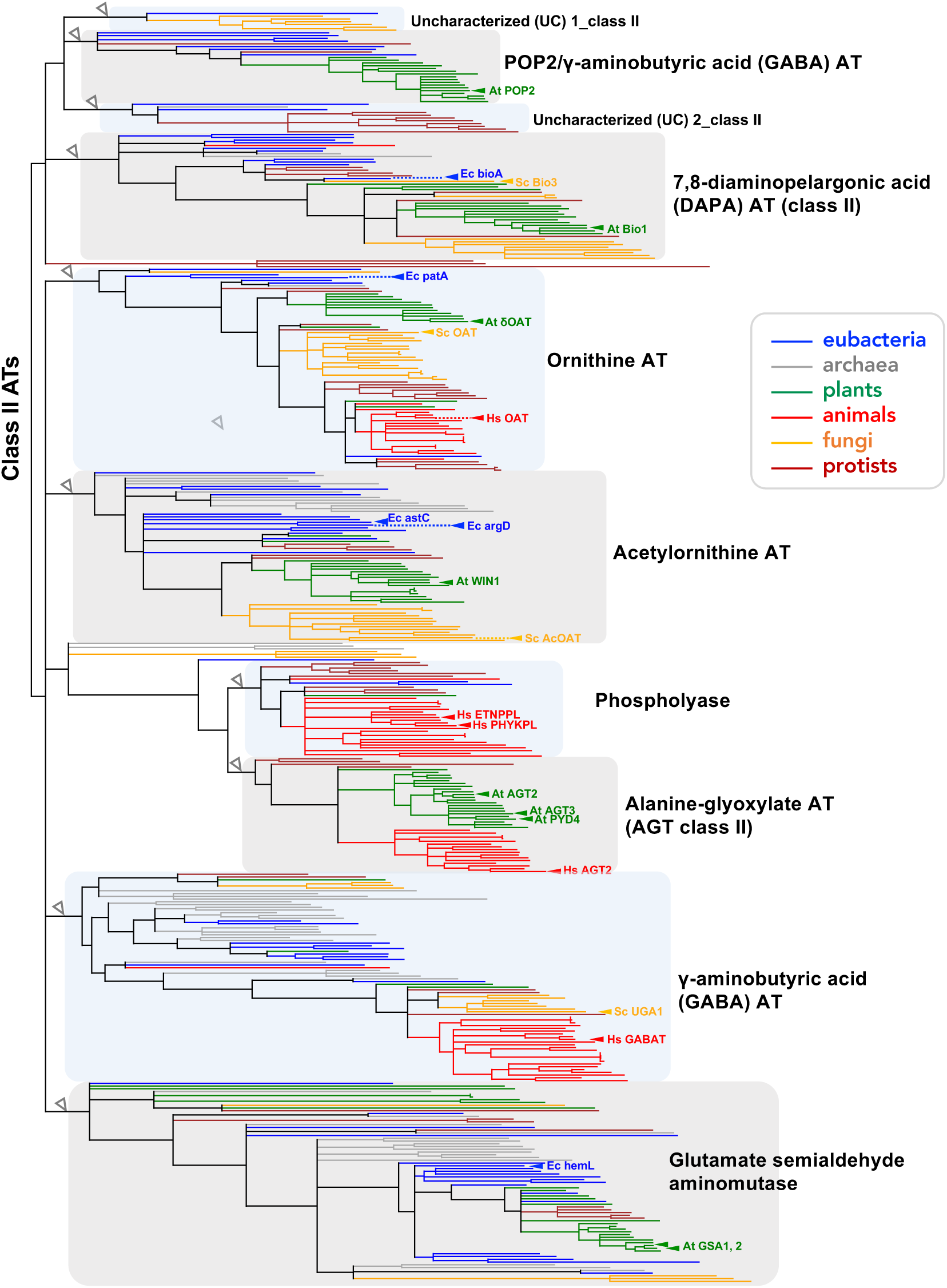
Phylogeny of **Class II** AT and related enzymes. Branch colors of blue, gray, green, red, orange, and maroon represent enzymes from eubacteria, archaea, plants, animals, fungi, and protists, receptively. Monophyletic AT groups are named according to the activities reported for the labeled enzymes from the four model organisms—*Arabidopsis thaliana* (green), *Homo sapiens* (red), *Saccharomyces cerevisiae* (orange), and *Escherichia coli* (blue). Groups without any representative enzymes are labeled as “uncharacterized (UC)”. Open arrow heads depict the root of each AT and related group. The corresponding full phylogenetic tree with all sequence IDs is provided as **Dataset S4**.

**Fig. S5.**
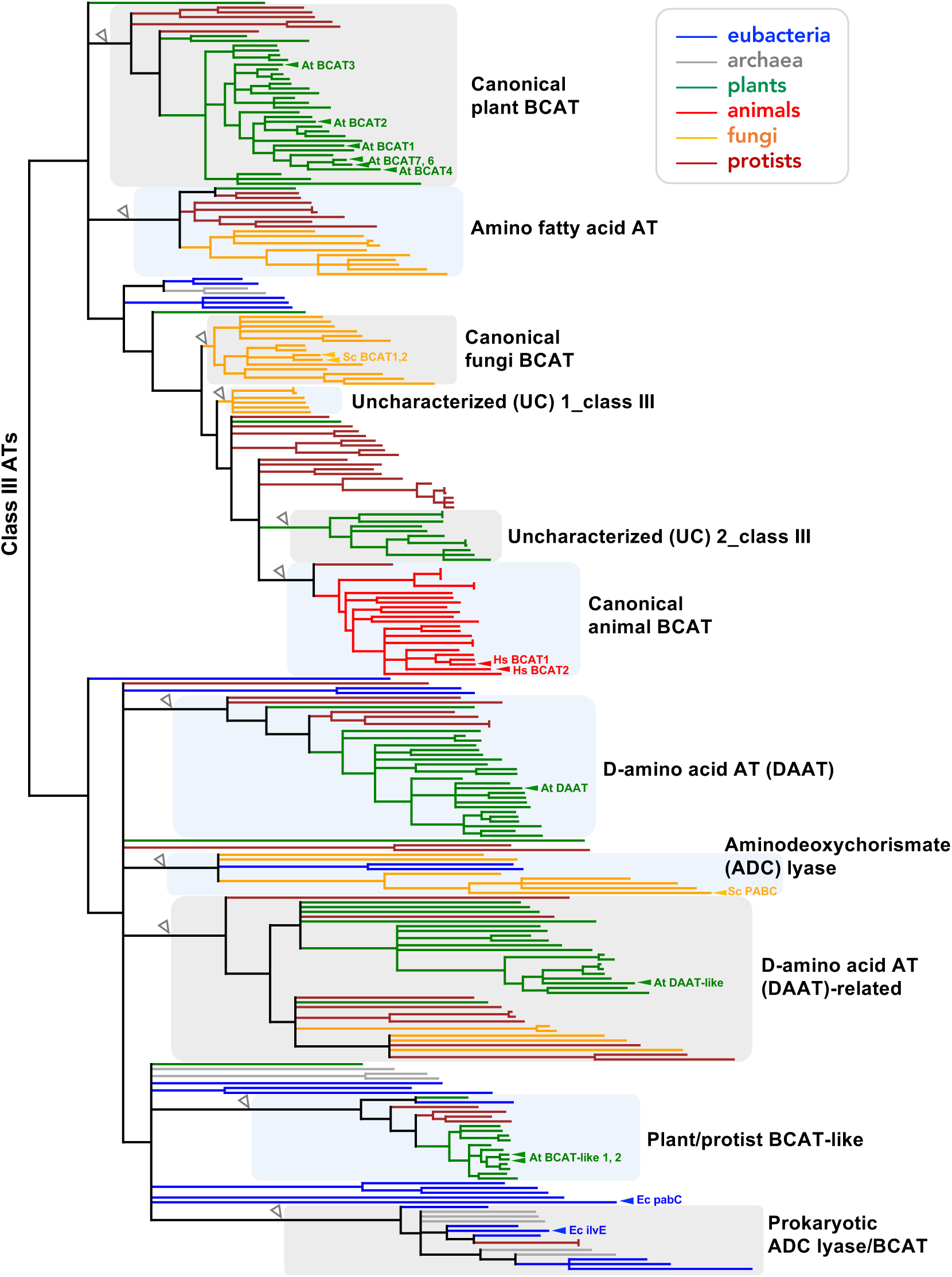
Phylogeny of **Class III** AT and related enzymes. Branch colors of blue, gray, green, red, orange, and maroon represent enzymes from eubacteria, archaea, plants, animals, fungi, and protists, receptively. Monophyletic AT groups are named according to the activities reported for the labeled enzymes from the four model organisms—*Arabidopsis thaliana* (green), *Homo sapiens* (red), *Saccharomyces cerevisiae* (orange), and *Escherichia coli* (blue). Groups without any representative enzymes are labeled as “uncharacterized (UC)”. Open arrow heads depict the root of each AT and related group. The corresponding full phylogenetic tree with all sequence IDs is provided as **Dataset S4**.

**Fig. S6.**
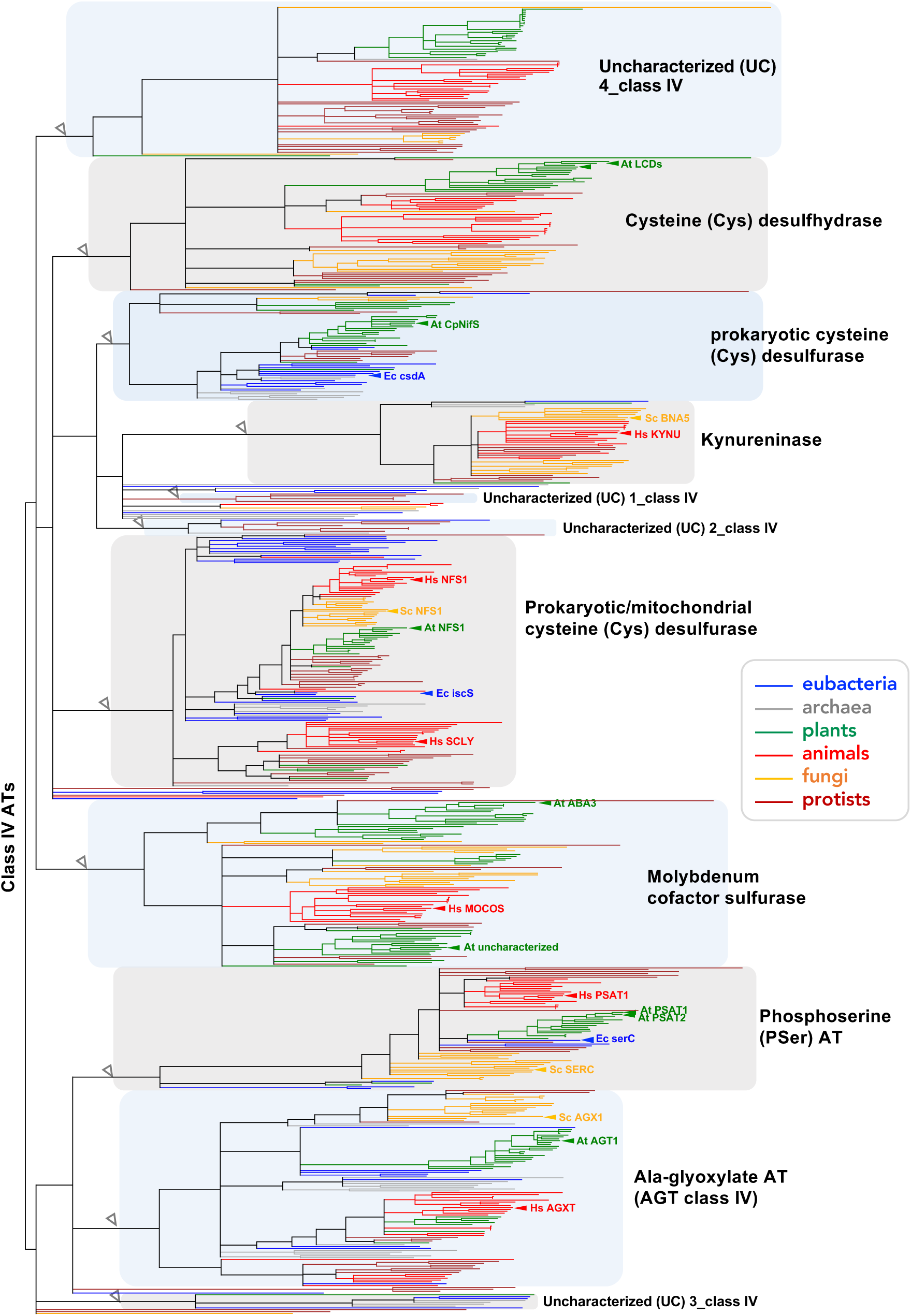
Phylogeny of **Class IV** AT and related enzymes. Branch colors of blue, gray, green, red, orange, and maroon represent enzymes from eubacteria, archaea, plants, animals, fungi, and protists, receptively. Monophyletic AT groups are named according to the activities reported for the labeled enzymes from the four model organisms—*Arabidopsis thaliana* (green), *Homo sapiens* (red), *Saccharomyces cerevisiae* (orange), and *Escherichia coli* (blue). Groups without any representative enzymes are labeled as “uncharacterized”. Open arrow heads depict the root of each AT and related group. The corresponding full phylogenetic tree with all sequence IDs is provided as **Dataset S4**.

**Fig. S7.**
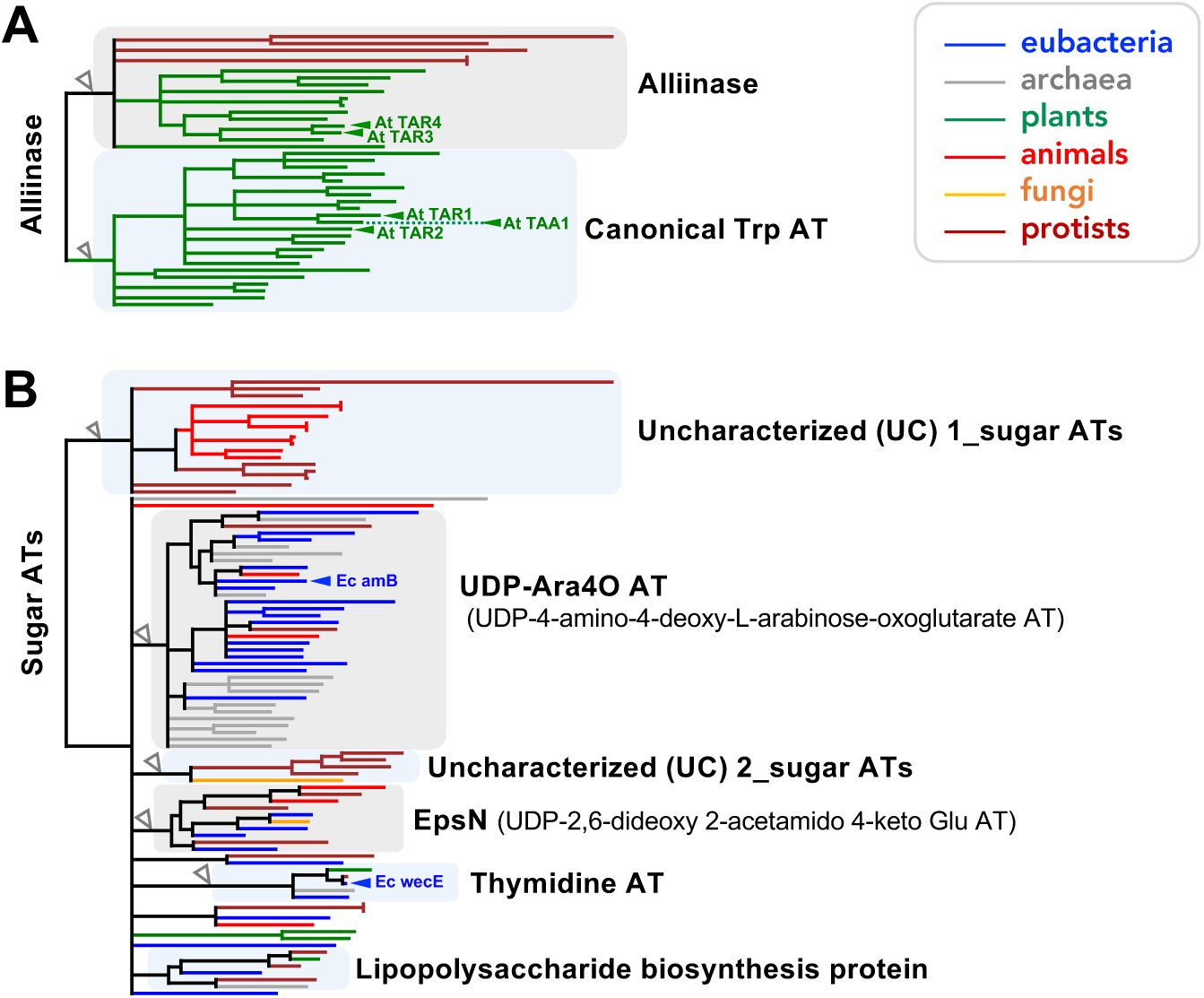
Phylogeny of *(A)* **Alliinases** and *(B)* **Sugar ATs** and related enzymes. Branch colors of blue, gray, green, red, orange, and maroon represent enzymes from eubacteria, archaea, plants, animals, fungi, and protists, receptively. Monophyletic AT groups are named according to the activities reported for the labeled enzymes from the four model organisms—*Arabidopsis thaliana* (green), *Homo sapiens* (red), *Saccharomyces cerevisiae* (orange), and *Escherichia coli* (blue). Groups without any representative enzymes are labeled as “uncharacterized (UC)”. Open arrow heads depict the root of each AT and related group. The corresponding full phylogenetic trees with all sequence IDs are provided as **Dataset S4**.

**Fig. S8.**
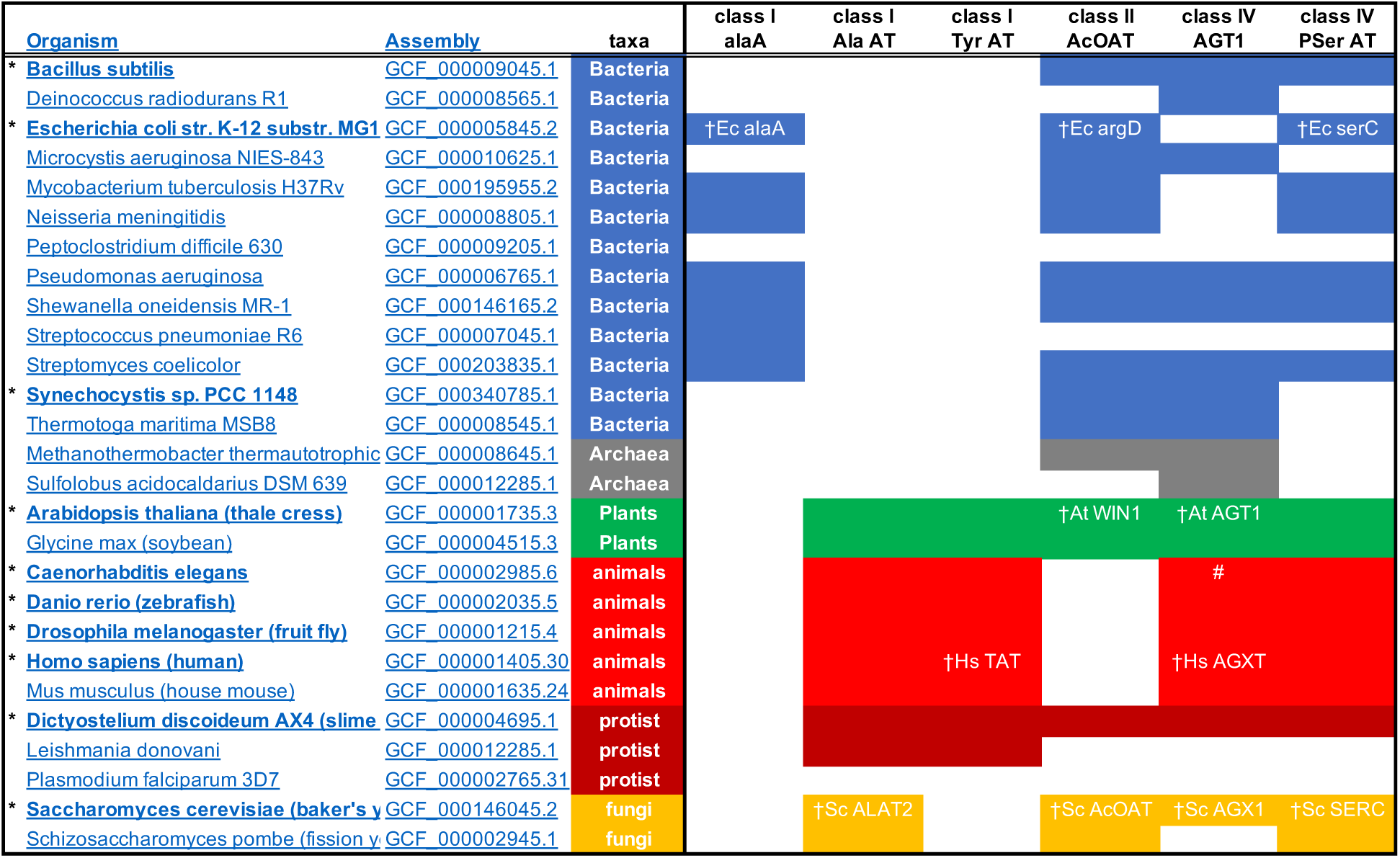
Distribution of selected AT groups across the 27 model species of the NCBI Landmark Database. The presence (filled boxes) and absence (white open boxes) of the indicated six AT groups were examined by BlastP search against the NCBI Landmark Proteome Database, derived from 27 genomes spanning a wide taxonomic range. The database utilizes the most recent and the best available representative assembly from each organism. Thus, the search provides taxonomically diverse non-redundant set of proteins supported by genomic assemblies. * Species also included in the 90 species used in the main analysis. ^†^ Query sequences are indicated by protein names. ^#^ Class IV AGT1 was represented by NP_495885.1 in the genome of *C. elegans* strain Bristol N2.

**Fig. S9.**
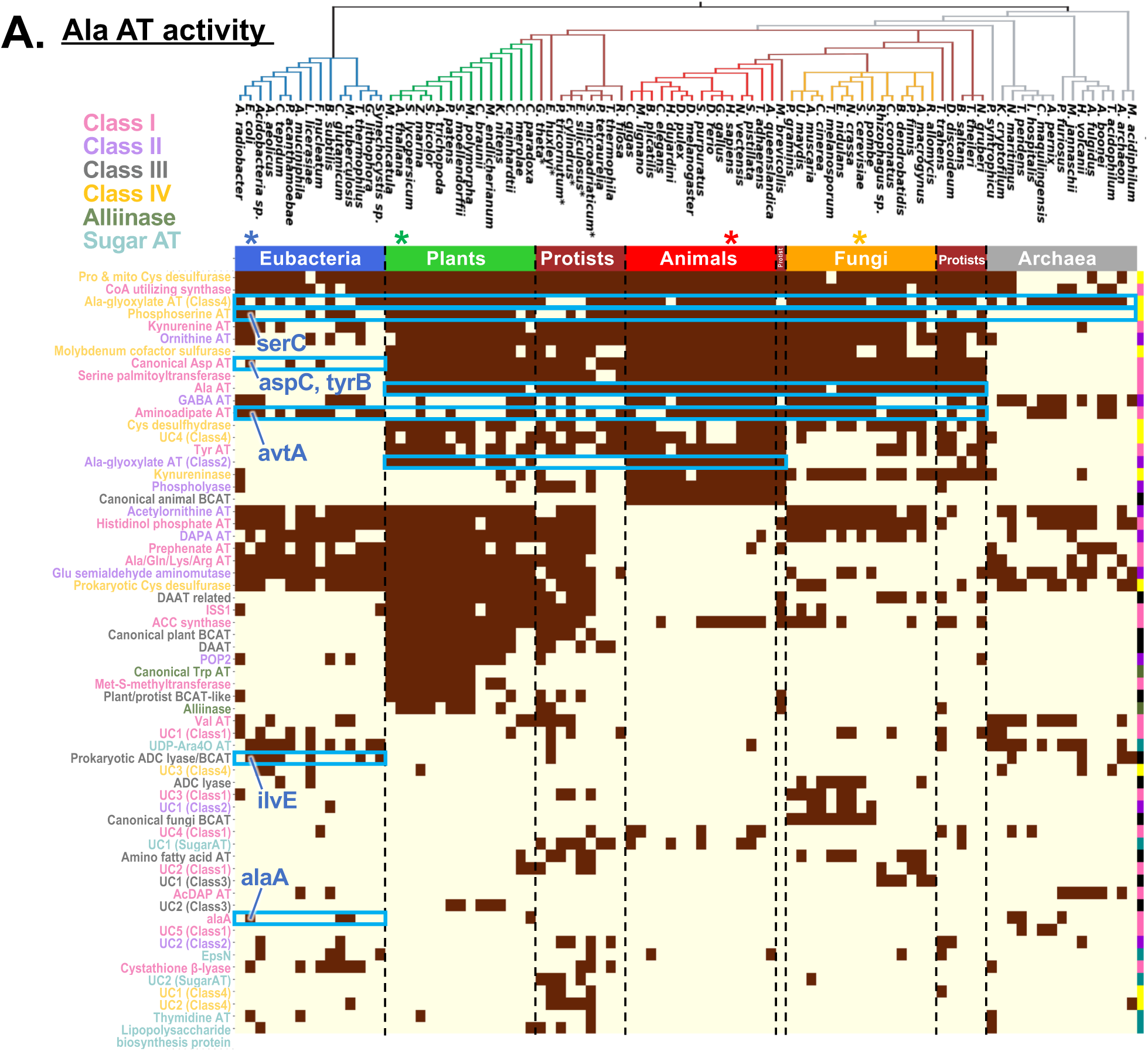
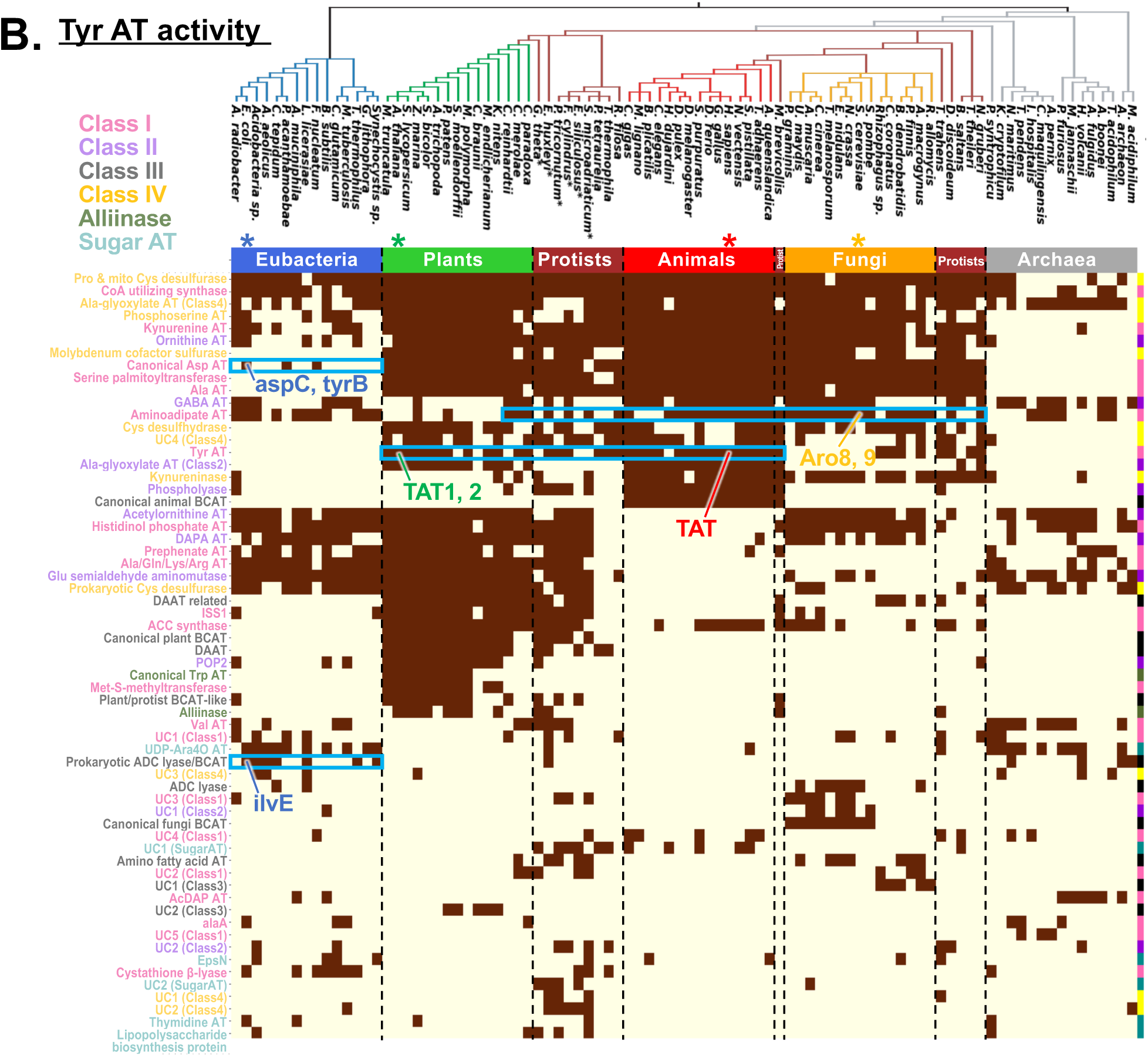
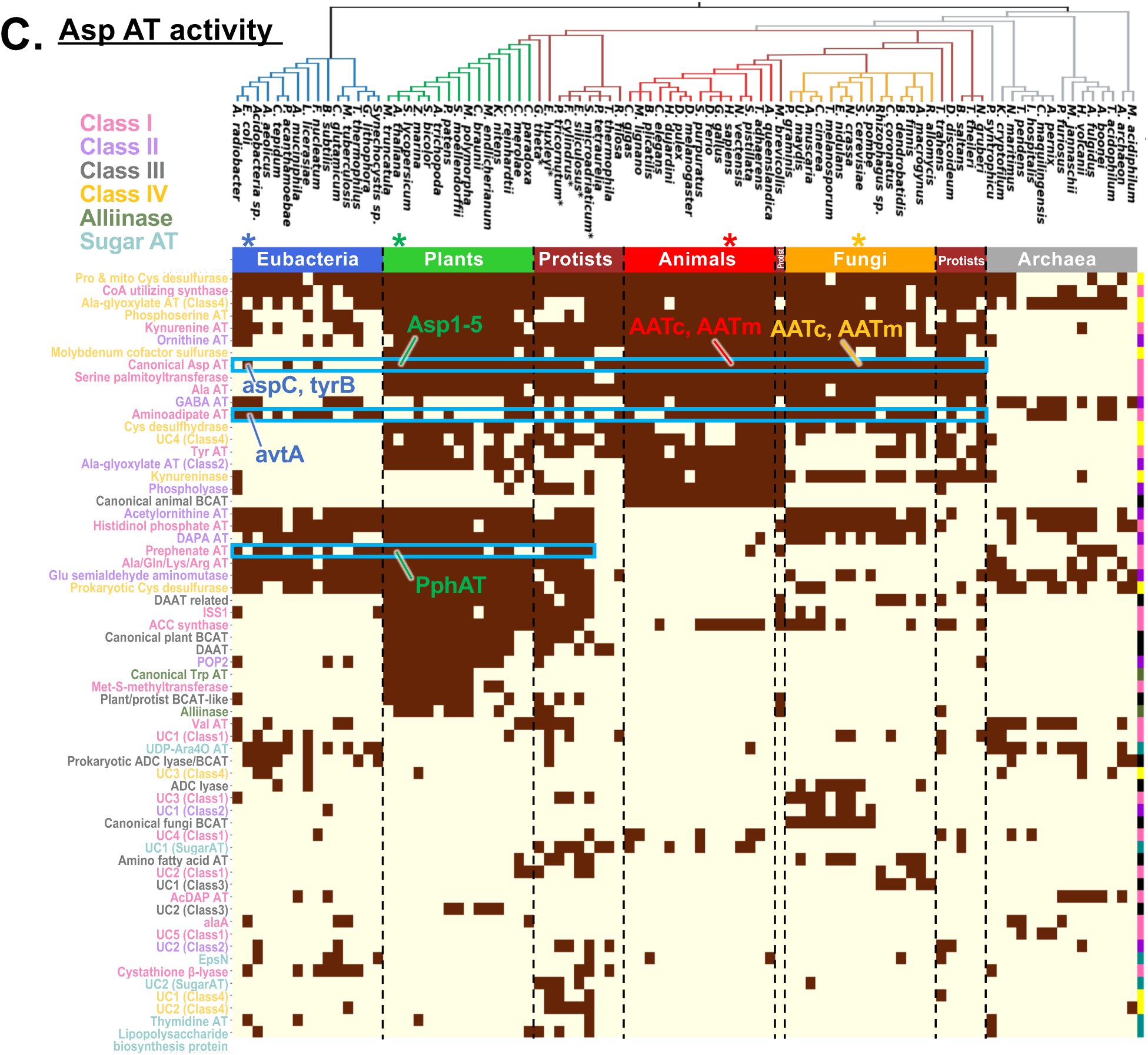
Examples of essential AT activities carried out by taxon-specific AT enzymes from distinct AT groups. Enzymes responsible for essential **(*A*)** Ala AT, **(*B*)** Tyr AT, and **(*C*)** Asp AT activities are marked by sky blue boxes on the same AT mapping result from Figure 2B. Corresponding enzymes are shown from representative model organisms from four kingdoms—*E. coli*, *A. thaliana*, *H. sapiens*, and *S. cerevisiae* (marked by *). Species were arranged based on the taxonomic relationship at the top: Gray, brown, orange, red, green and blue depict archaea, protists, fungi, animals, plants and eubacteria, respectively. Brown boxes indicate certain enzymes (left) are present in the corresponding species (top).

**Fig. S10.**
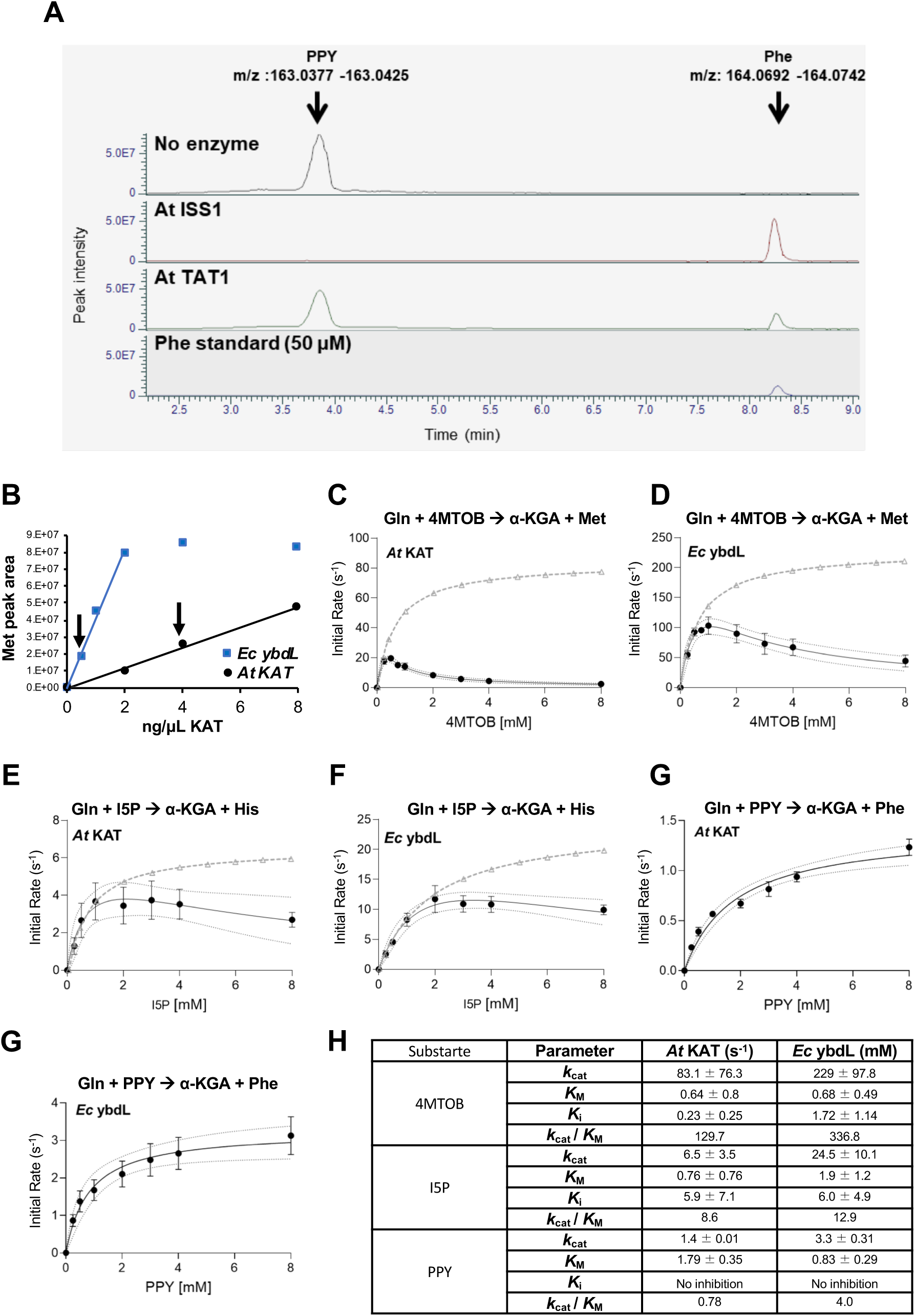
*In vitro* characterization of ATs. ***(A)*** LC/MS based detection of AT catalyzed conversion of keto acids to amino acids. Keto acid PPY is converted to Phe by ATs ISS1 and TAT1. Spectra is filtered based on the predicted mass-to-charge ratio (m/z) PPY and Phe. No enzyme and authentic Phe standard samples are used to validate the retention times of detected peaks. PPY, phenylpyruvate. ***(B)*** Determination of the range where methionine production linearly increases with the amount of enzyme added to the reaction. Arrows show the enzyme concentrations used for the kinetic studies. ***(C to G)*** Kinetic characterization of *A. thaliana* (*At*) KAT and *E. coli* (*Ec*) ybdL. Enzymatic activity of *Ec* ybdL and *At* KAT were tested with 10 mM Gln and 0 to 8 mM of 4MTOB, I5P or PPY. Each data point is an average of three separate assays (n=3). Error bars show standard error of the mean (SEM). For 4MTOB and I5P, a modified Michaelis Menten equation with substrate inhibition was fitted using non-linear regression. Dash lines with triangle datapoints depict hypothetical curves if there was no substrate inhibition. ***(H)*** Summary of kinetic parameters obtained in the study.

**Fig. S11.**
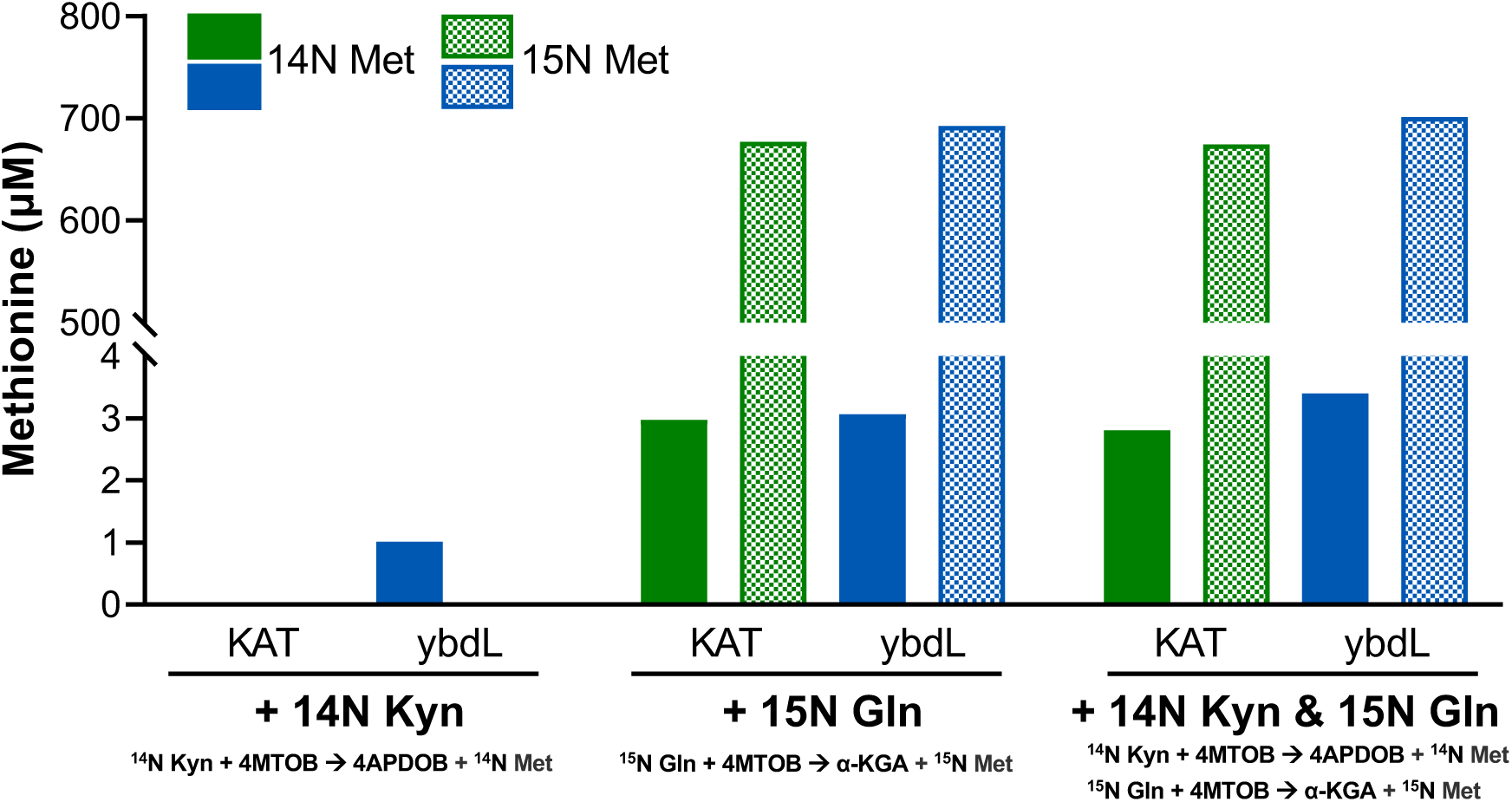
Kynurenine and glutamine AT activities of *At* KAT and *Ec* ybdL. Ability and preference of *At* KAT (green bars) and *Ec* ybdL (blue bars) to utilize kynurenine (Kyn) and/or glutamine (Gln) as amino donor was tested by three separate AT assays that utilized ^14^N-Kyn, amino-^15^N-Gln or ^14^N-Kyn + amino-^15^N-Gln as donors and 4MTOB as acceptor. Formation of ^14^N (solid colors) or ^15^N (crosshatched) methionine (Met) was detected using LC/MS and converted to concentration using a Met standard curve. Both enzymes were able to effectively use Gln as amino donor, forming ^15^N-Met. When Kyn was the sole amino donor, only *Ec* ybdL was able to form ^14^N Met, but the conversion amount was very low. Formation of ^14^N Met in the assays with amino-^15^N-Gln as donor was attributed to the isotopic impurity of Gln (98% atom pure according to the vendor). When both Gln and Kyn was used as amino donors, only *Ec* ybdL had increased production of 14N Met, which further supported the very minor utilization of Kyn by *Ec* ybdL. 4APDOB, 4-(2-aminophenyl)-2,4-dioxobutanoate.

**Fig. S12.**
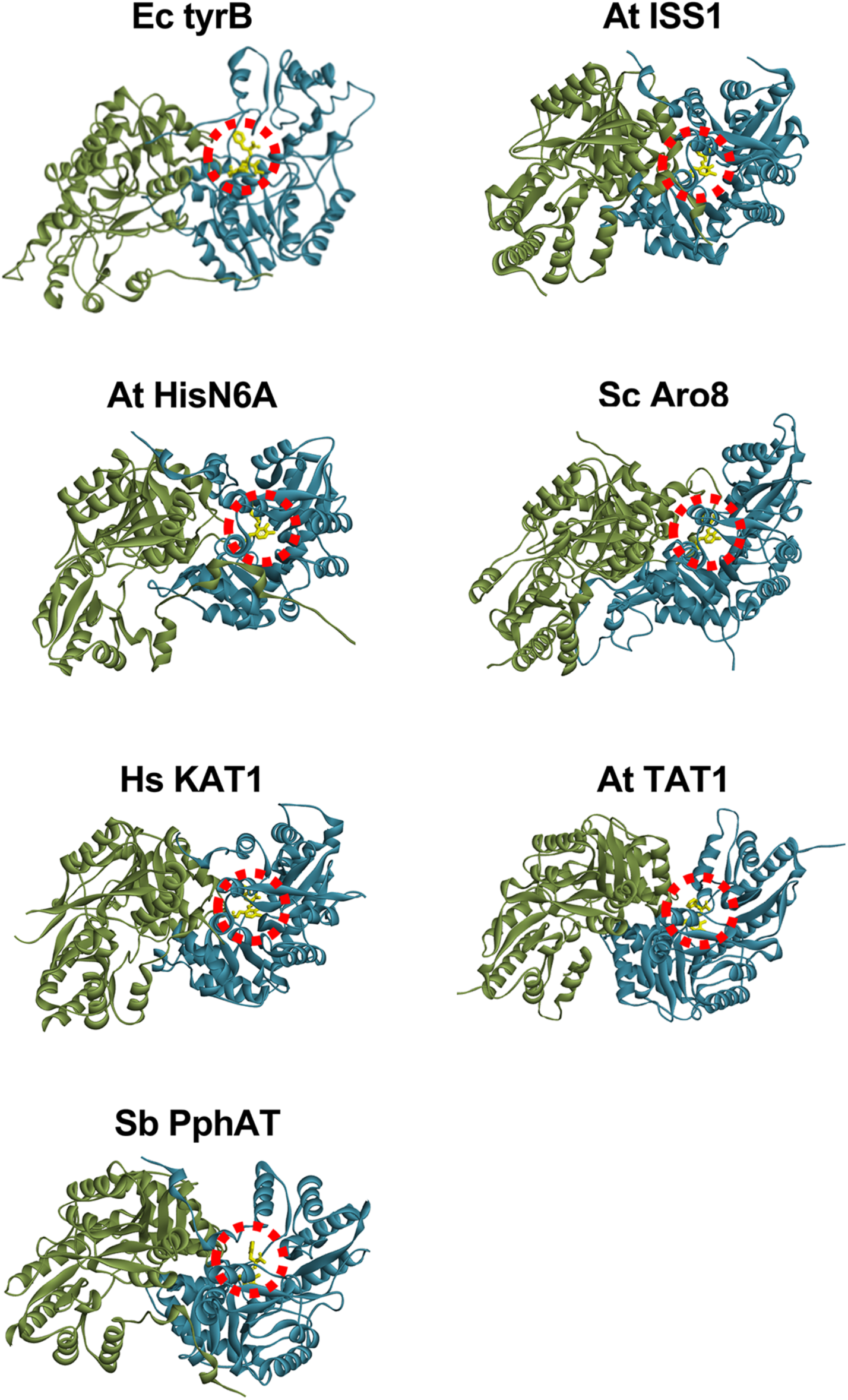
Docking of PLP-Phe aldimine form (yellow molecule shown inside red circle) to the active site of Aro ATs using Autodock v4.2.6 ^7^. Green and blue show each monomer of the AT dimer.

**Fig. S13.**
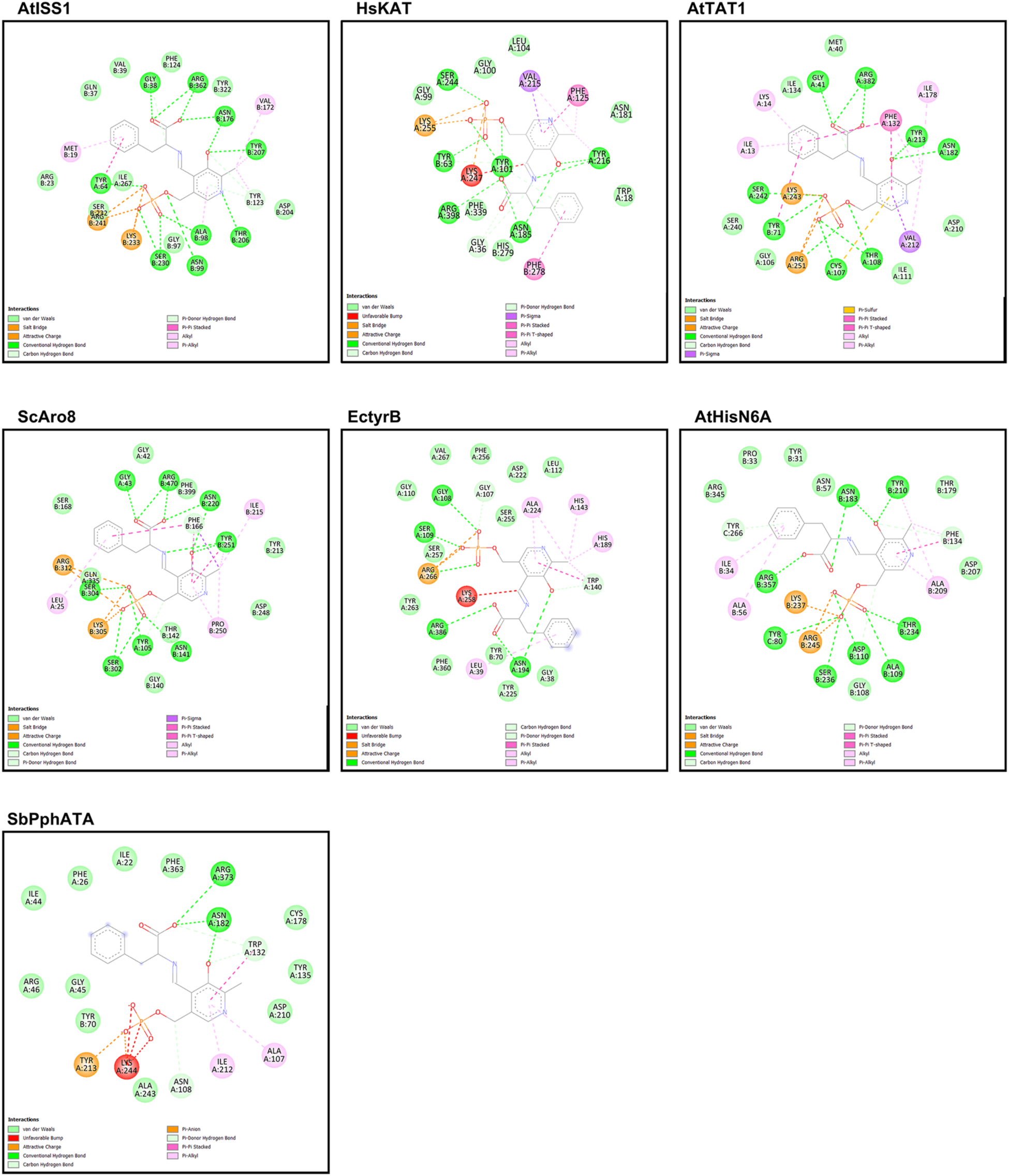
The 2-D diagrams for residues that interact with the PLP-Phe aldimine in Aro AT active sites were determined using Autodock v4.2.6^7^ and Discovery studio 2021 (Dassault Systèmes, Vélizy-Villacoublay, France). Dashed lines represent different types of interactions, which are named under each diagram.

**Fig. S14.**
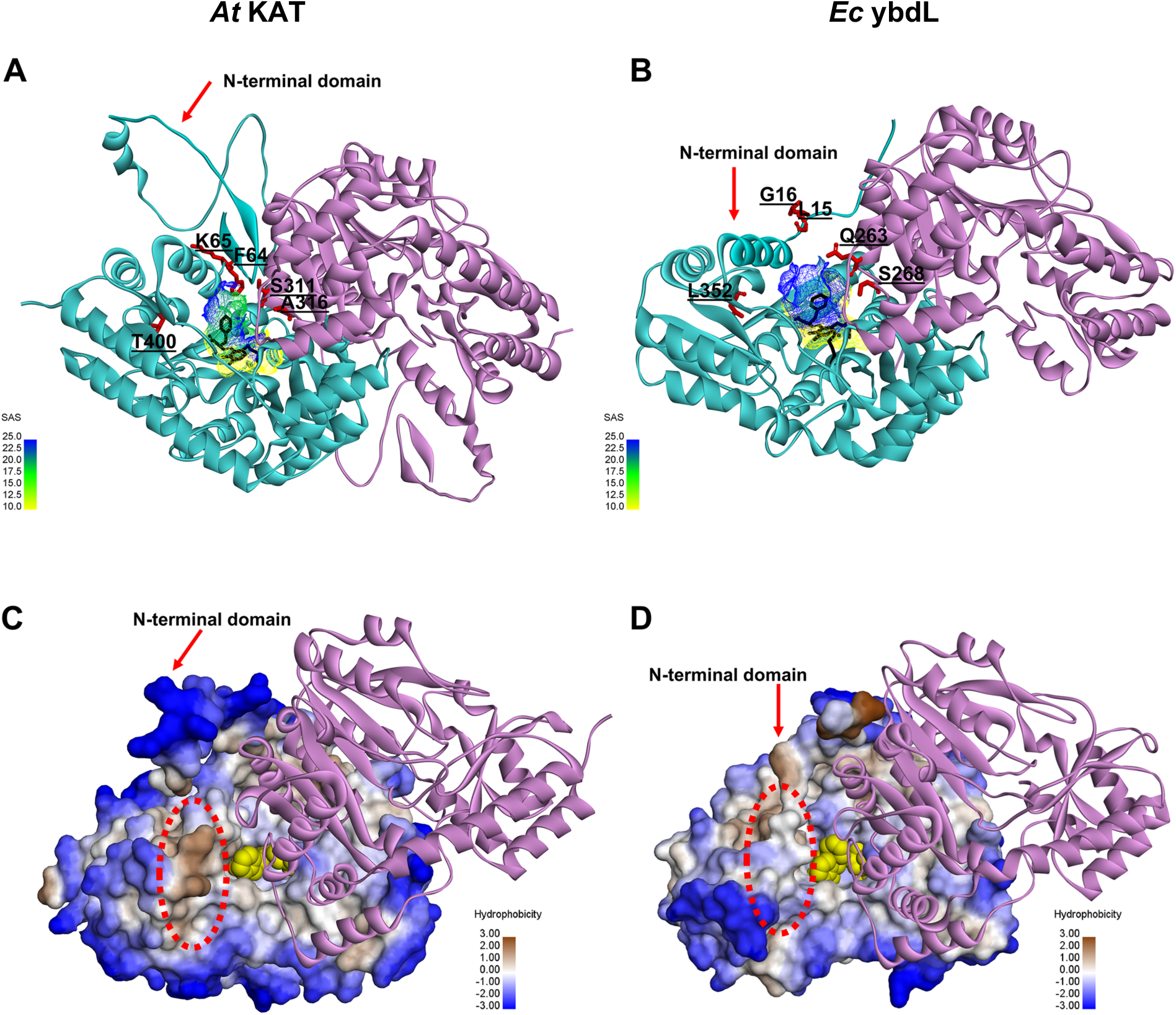
Structural differences between *At* KAT and *Ec* ybdL. Five residues around the active site entrance (red sticks) that are not conserved between ***(A)*** *At* KAT and ***(B)*** *Ec* ybdL may influence the higher solvent accessible surface (SAS) inside the ybdL active site. Additionally, differences in hydrophobicity of the surface (highlighted with the red circle) below the flexible N-terminal domain of ***(C)*** *At* KAT and ***(D)*** *Ec* ybdL that may influence the open/closed conformational change^8–10^.

**Table S1:**
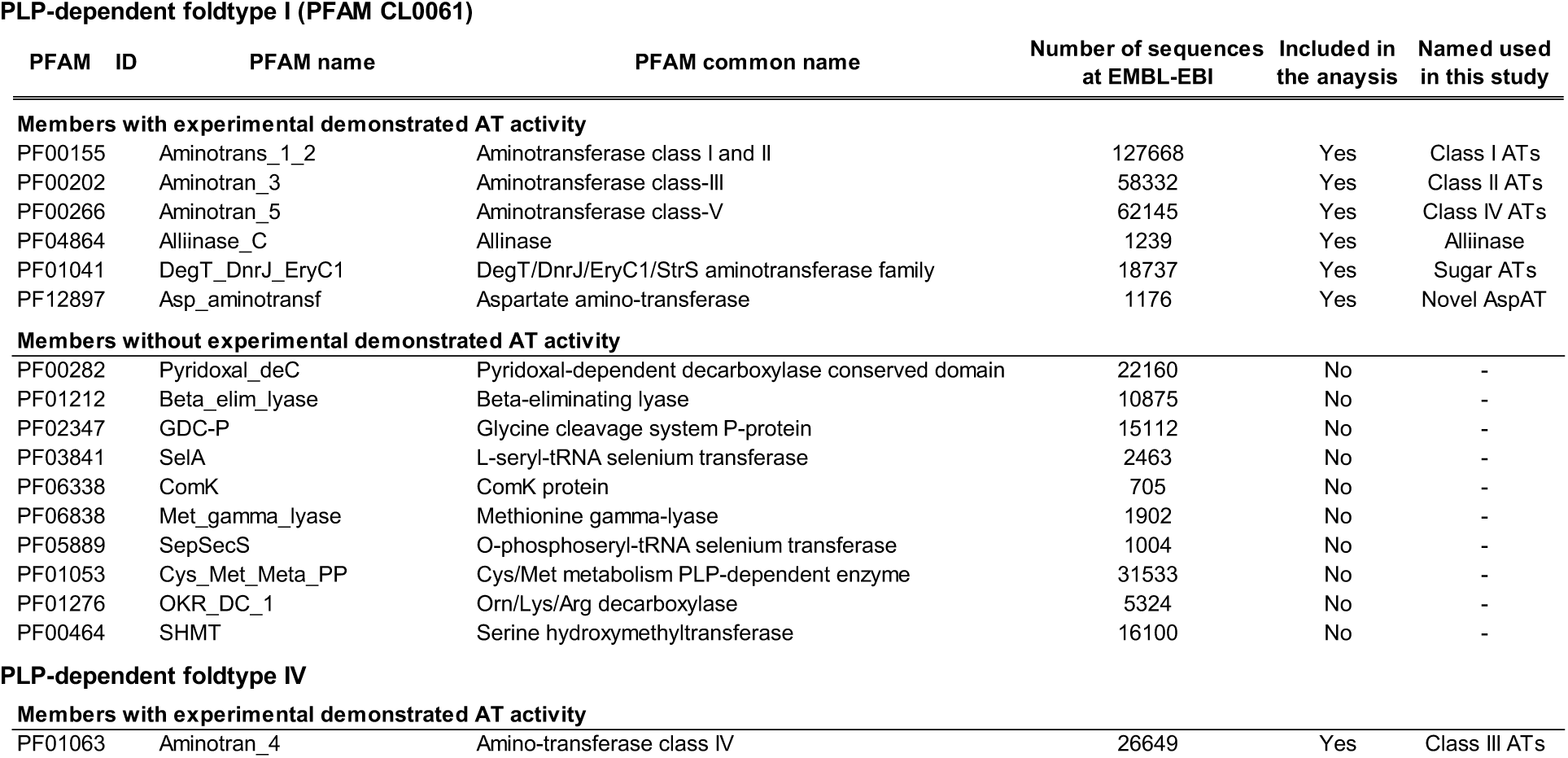
AT-like PFAM families from PLP-fold type I and IV enzymes.

**Table S2:**
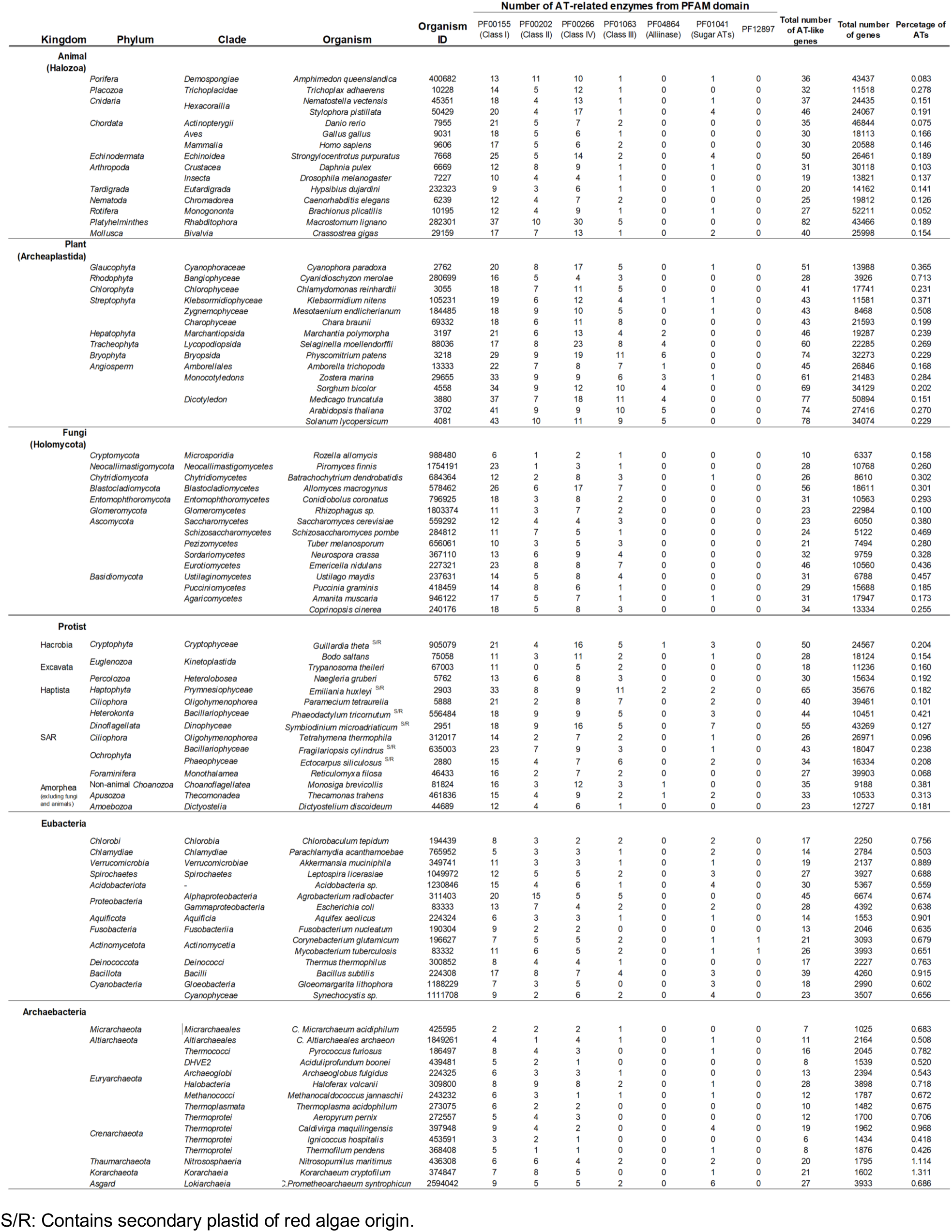
List of PFAM domains and species used for the phylogeny.

**Table S3.** AT enzymes used for substrate specificity screening.

| AT Group | Origin | Name | Gene number | Group Status | Enzyme Status |
| --- | --- | --- | --- | --- | --- |
| ISS1 (or also known as VAS1) | *Arabidopsis thaliana* | ISS1 | AT1G80360 | Canonical Aro AT group | Aro AT |
| Histidinol phosphate AT | *Arabidopsis thaliana* | HisN6A<br>HisN6B | AT5G10330<br>AT1G71920 | Non-canonical Aro AT group | No evidence for Aro<br>AT activity |
|  | *Escherichia coli* | hisC | P06986 |  | No evidence for Aro<br>AT activity |
| Tyrosine AT | *Arabidopsis thaliana* | TAT1 | AT5G53970 | Canonical Aro AT group | Aro AT |
|  |  | SUR1 | AT2G20610 |  | Non-AT (lyase) |
| Alanine AT | *Arabidopsis thaliana* | AlaAT1 | AT1G17290 | Non-Aro AT group<br>(Sister to Tyr ATs) | No evidence for Aro<br>AT activity |
| alaA | *Escherichia coli* | alaA | P0A959 | Non-Aro AT group<br>(Sister to Tyr ATs) | No evidence for Aro<br>AT activity |
| Kynurenine AT | *Arabidopsis thaliana* | KAT | AT1G77670 | Canonical Aro AT group | No evidence for Aro<br>AT activity |
|  | *Escherichia coli* | ybdL | P77806 |  | Aro AT |
| Prephenate AT | *Arabidopsis thaliana* | PphAT | AT2G22250 | Non-canonical Aro AT group | No evidence for Aro<br>AT activity |

**Table S4:** Percent identities of ATs.

|  | At ISS1 | At HisN6A | At HisN6B | Ec hisC | At SUR1 | At TAT1 | At AlaAT1 | Ec alaA | At KAT | Ec ybdL | At PphAT |
| --- | --- | --- | --- | --- | --- | --- | --- | --- | --- | --- | --- |
| **At ISS1** |  | 21.8 | 23.0 | 21.5 | 24.9 | 23.7 | 25.0 | 21.5 | 22.8 | 26.2 | 27.3 |
| **At HisN6A** | 21.8 |  | 99.0 | 31.6 | 20.6 | 24.9 | 23.1 | 22.4 | 25.9 | 25.0 | 26.5 |
| **At HisN6B** | 23.0 | 99.0 |  | 31.8 | 20.6 | 24.9 | 23.1 | 22.4 | 25.9 | 25.0 | 25.9 |
| **Ec hisC** | 21.5 | 31.6 | 31.8 |  | 25.2 | 24.2 | 29.7 | 25.3 | 23.8 | 24.0 | 27.7 |
| **At SUR1** | 24.9 | 20.6 | 20.6 | 25.2 |  | 52.0 | 22.4 | 27.0 | 26.2 | 24.4 | 22.6 |
| **At TAT1** | 23.7 | 24.9 | 24.9 | 24.2 | 52.0 |  | 26.7 | 24.9 | 25.2 | 22.7 | 21.4 |
| **At AlaAT1** | 25.0 | 23.1 | 23.1 | 29.7 | 22.4 | 26.7 |  | 27.6 | 21.9 | 23.5 | 22.7 |
| **Ec alaA** | 21.5 | 22.4 | 22.4 | 25.3 | 27.0 | 24.9 | 27.6 |  | 27.9 | 25.1 | 31.9 |
| **At KAT** | 22.8 | 25.9 | 25.9 | 23.8 | 26.2 | 25.2 | 21.9 | 27.9 |  | 40.0 | 30.2 |
| **Ec ybdL** | 26.2 | 25.0 | 25.0 | 24.0 | 24.4 | 22.7 | 23.5 | 25.1 | 40.0 |  | 29.6 |
| **At PphAT** | 27.3 | 26.5 | 25.9 | 27.7 | 22.6 | 21.4 | 22.7 | 31.9 | 30.2 | 29.6 |  |

**Table S5.** Substrate specificity of Aromatic AT. Percent conversion of keto acids to amino acids by different Aro AT and control enzymes, as shown in **Fig. 3*A***.

**Average**
|  | ISS1 | HisN6A | HisN6B | hisC | KAT | ybdL | TAT1 | alaA | AlaAT1 | PphAT | SUR1 |
| --- | --- | --- | --- | --- | --- | --- | --- | --- | --- | --- | --- |
| **Phe** | 109.36 | 5.07 | 5.22 | 3.64 | 12.76 | 76.93 | 51.88 | 0.83 | 0.13 | 0.55 | 0.06 |
| **Aro** | 0.01 | 0.01 | 0.01 | 0.01 | 0.01 | 0.00 | 0.00 | 0.01 | 0.44 | 21.39 | 0.01 |
| **Tyr** | 95.08 | 8.09 | 6.48 | 10.42 | -4.24 | 12.91 | 65.45 | 0.40 | -0.39 | 0.12 | 1.63 |
| **Trp** | 92.32 | 15.26 | 15.66 | 63.06 | 0.37 | 0.72 | 35.32 | 3.55 | 0.12 | 0.03 | 0.00 |
| **Met** | 80.04 | 0.43 | 0.43 | 6.06 | 80.78 | 77.44 | 15.88 | 1.18 | 0.36 | 0.53 | 0.00 |
| **Leu** | 40.67 | 0.12 | 0.13 | 1.20 | 4.82 | 11.68 | 7.85 | 0.30 | 0.16 | 0.12 | 0.04 |
| **Ile** | 0.22 | 0.03 | 0.02 | 0.01 | 0.00 | 0.00 | 0.01 | 0.00 | 0.05 | 0.01 | 0.02 |
| **Val** | 0.37 | 0.02 | 0.03 | 0.04 | 0.03 | 0.03 | 0.03 | 0.04 | 0.06 | 0.03 | 0.03 |
| **His** | 73.49 | 42.90 | 44.08 | 57.83 | 67.86 | 61.87 | 10.25 | 5.37 | 0.25 | 0.26 | 0.02 |
| **Ala** | 4.01 | 0.13 | 0.13 | 2.11 | 5.00 | 14.52 | 0.35 | 48.71 | 52.20 | 0.27 | 0.11 |
| **Ser\*** | 2.55 | 0.05 | 0.09 | 0.64 | 3.92 | 10.10 | 0.10 | 0.19 | 1.79 | 0.15 | 0.27 |
| **hAla** | 18.05 | 0.18 | 0.18 | 3.18 | 32.44 | 33.35 | 1.83 | 22.44 | 50.75 | 0.03 | 0.00 |
| **Asp** | 0.05 | 0.11 | 0.11 | 0.39 | 0.01 | 0.00 | 0.04 | 0.11 | 0.11 | 0.64 | 0.02 |
| **Glu** | 1.22 | N/A | N/A | N/A | 0.56 | 3.00 | N/A | N/A | N/A | N/A | N/A |
| **AAD** | 0.12 | 2.39 | 2.77 | 20.06 | 0.00 | 0.03 | 1.19 | 2.71 | 1.58 | 7.15 | 0.00 |

**Standard Deviation**
|  | ISS1 | HisN6A | HisN6B | hisC | KAT | ybdL | TAT1 | alaA | AlaAT1 | PphAT | SUR1 |
| --- | --- | --- | --- | --- | --- | --- | --- | --- | --- | --- | --- |
| **Phe** | 5.61 | 0.24 | 0.64 | 0.34 | 1.56 | 4.55 | 9.66 | 0.29 | 0.05 | 0.12 | 0.01 |
| **Aro** | 0.01 | 0.01 | 0.01 | 0.01 | 0.01 | 0.00 | 0.00 | 0.01 | 0.04 | 1.11 | 0.01 |
| **Tyr** | 1.24 | 4.35 | 2.20 | 4.99 | 7.14 | 1.51 | 6.65 | 1.67 | 2.70 | 5.07 | 5.17 |
| **Trp** | 9.38 | 1.21 | 2.39 | 10.48 | 0.05 | 0.06 | 8.73 | 1.51 | 0.01 | 0.01 | 0.00 |
| **Met** | 2.53 | 0.02 | 0.04 | 0.51 | 4.45 | 5.70 | 3.50 | 0.42 | 0.04 | 0.07 | 0.00 |
| **Leu** | 7.02 | 0.01 | 0.02 | 0.11 | 0.72 | 1.19 | 1.89 | 0.10 | 0.06 | 0.01 | 0.02 |
| **Ile** | 0.30 | 0.02 | 0.02 | 0.01 | 0.00 | 0.01 | 0.01 | 0.00 | 0.05 | 0.02 | 0.02 |
| **Val** | 0.06 | 0.01 | 0.00 | 0.02 | 0.00 | 0.01 | 0.01 | 0.01 | 0.03 | 0.00 | 0.00 |
| **His** | 3.96 | 2.22 | 4.61 | 8.24 | 2.98 | 7.33 | 2.72 | 2.21 | 0.01 | 0.06 | 0.01 |
| **Ala** | 0.89 | 0.03 | 0.01 | 0.30 | 0.65 | 1.52 | 0.06 | 9.28 | 5.53 | 0.05 | 0.01 |
| **Ser** | N/A | N/A | N/A | N/A | N/A | N/A | N/A | N/A | N/A | N/A | N/A |
| **hAla** | 3.53 | 0.02 | 0.01 | 0.40 | 3.68 | 3.07 | 0.37 | 7.90 | 0.54 | 0.01 | 0.01 |
| **Asp** | 0.05 | 0.03 | 0.01 | 0.07 | 0.01 | 0.00 | 0.01 | 0.04 | 0.08 | 0.15 | 0.01 |
| **Glu** | 1.23 | N/A | N/A | N/A | 0.48 | 0.73 | N/A | N/A | N/A | N/A | N/A |
| **AAD** | 0.02 | 0.17 | 0.38 | 2.25 | 0.00 | 0.01 | 0.28 | 0.94 | 0.03 | 1.22 | 0.00 |

**Table S6.**
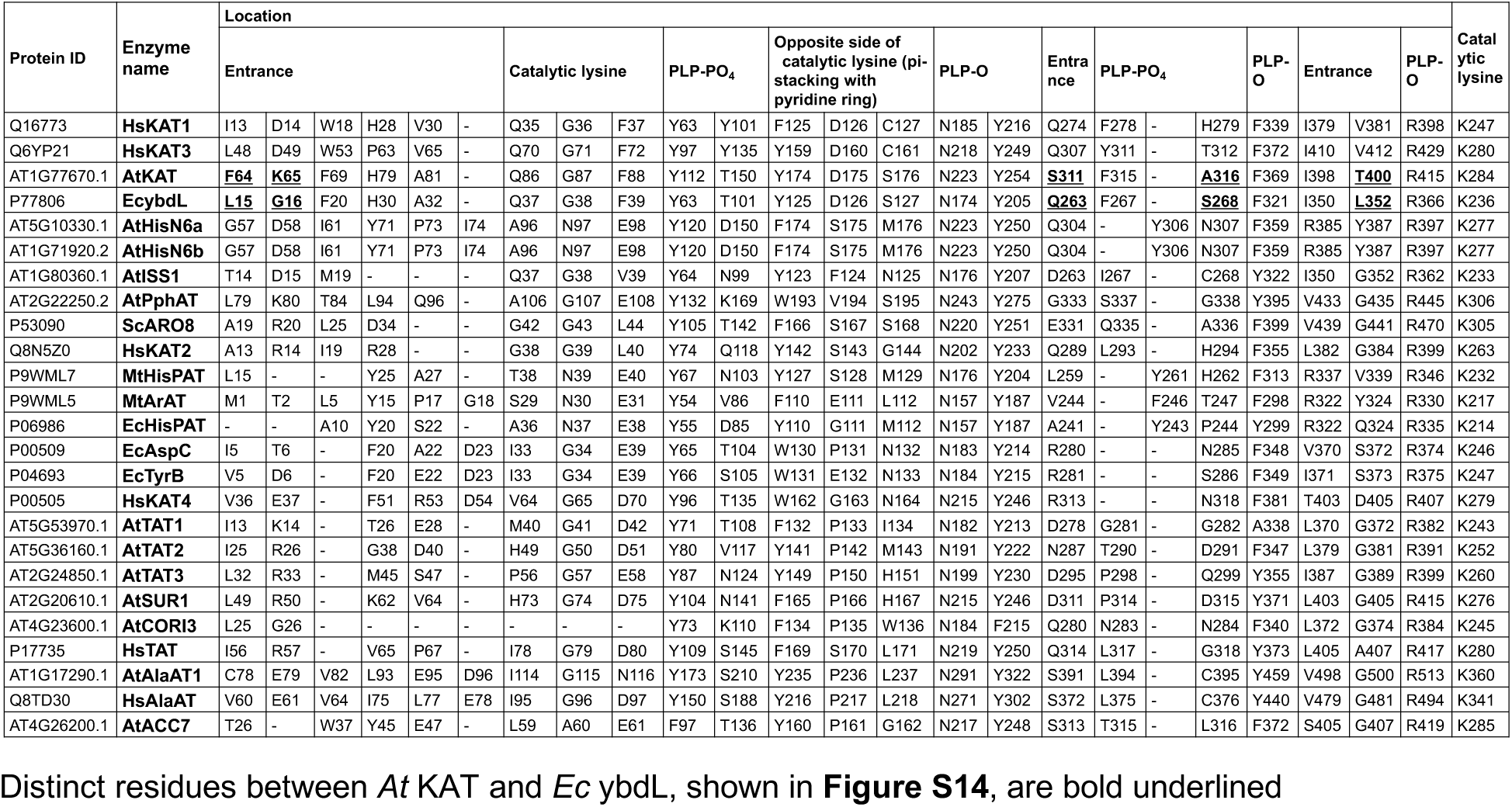
Residues involved in substrate recognition of canonical Aro ATs and related enzymes.

## Supplemental Datasets

The following materials are available in the online version of this article.

**Dataset S1: Corrected PFAMs.**

**Dataset S2: AT and related sequences analyzed in this study.**

**Dataset S3: Statistical significance of number of ATs between different taxonomic groups.**

**Dataset S4: ToL-scale phylogenetic trees of AT and related enzymes with sequence IDs.**

**Dataset S5: AT copy numbers in different species.**

**Dataset S6: Representative AT structures used to determine active sites.**

**Dataset S7: Sequence database IDs of 90 species used in this study.**

**Dataset S8: Aro AT structures used for the docking analysis.**

## References

1. Yčas, M. On earlier states of the biochemical system. Journal of Theoretical Biology 44, 145–160 (1974).

2. Jensen, R. A. Enzyme Recruitment in Evolution of New Function. Annual Review of Microbiology 30, 409–425 (1976).

3. Ferla, M. P., Brewster, J. L., Hall, K. R., Evans, G. B. & Patrick, W. M. Primordial-like enzymes from bacteria with reduced genomes. Molecular Microbiology 105, 508–524 (2017).

4. Wagner, A. Energy costs constrain the evolution of gene expression. Journal of Experimental Zoology Part B: Molecular and Developmental Evolution 308B, 322–324 (2007).

5. Nam, H. et al. Network Context and Selection in the Evolution to Enzyme Specificity. Science 337, 1101–1104 (2012).

6. Noffke, N., Christian, D., Wacey, D. & Hazen, R. M. Microbially Induced Sedimentary Structures Recording an Ancient Ecosystem in the ca. 3.48 Billion-Year-Old Dresser Formation, Pilbara, Western Australia. Astrobiology 13, 1103–1124 (2013).

7. Bell, E. A., Boehnke, P., Harrison, T. M. & Mao, W. L. Potentially biogenic carbon preserved in a 4.1 billion-year-old zircon. Proc Natl Acad Sci U S A 112, 14518–14521 (2015).

8. Betts, H. C. et al. Integrated genomic and fossil evidence illuminates life’s early evolution and eukaryote origin. Nat Ecol Evol 2, 1556–1562 (2018).

9. Ditzler, M. A., Popović, M. & Zajkowski, T. Chapter 5 - From building blocks to cells. in New Frontiers in Astrobiology (eds. Thombre, R. & Vaishampayan, P.) 111–133 (Elsevier, 2022). doi:10.1016/B978-0-12-824162-2.00010-5.

10. Pandurangan, A. P., Stahlhacke, J., Oates, M. E., Smithers, B. & Gough, J. The SUPERFAMILY 2.0 database: a significant proteome update and a new webserver. Nucleic Acids Research 47, D490–D494 (2019).

11. Petrov, A. S. et al. Evolution of the ribosome at atomic resolution. Proceedings of the National Academy of Sciences 111, 10251–10256 (2014).

12. Nagel, G. M. & Doolittle, R. F. Phylogenetic analysis of the aminoacyl-tRNA synthetases. J Mol Evol 40, 487–498 (1995).

13. Fournier, G. P. & Alm, E. J. Ancestral Reconstruction of a Pre-LUCA Aminoacyl-tRNA Synthetase Ancestor Supports the Late Addition of Trp to the Genetic Code. J Mol Evol 80, 171–185 (2015).

14. Rubio Gomez, M. A. & Ibba, M. Aminoacyl-tRNA synthetases. RNA 26, 910–936 (2020).

15. Smith, K. S., Jakubzick, C., Whittam, T. S. & Ferry, J. G. Carbonic anhydrase is an ancient enzyme widespread in prokaryotes. Proc Natl Acad Sci U S A 96, 15184–15189 (1999).

16. Rawlings, N. D. & Barrett, A. J. Evolutionary families of peptidases. Biochem J 290 (Pt 1), 205– 218 (1993).

17. Rawlings, N. D., Barrett, A. J. & Bateman, A. MEROPS: the peptidase database. Nucleic Acids Res 38, D227–233 (2010).

18. Weiss, M. C., Preiner, M., Xavier, J. C., Zimorski, V. & Martin, W. F. The last universal common ancestor between ancient Earth chemistry and the onset of genetics. PLOS Genetics 14, e1007518 (2018).

19. Copley, S. D. Evolution of new enzymes by gene duplication and divergence. FEBS J 287, 1262– 1283 (2020).

20. Ochman, H., Lawrence, J. G. & Groisman, E. A. Lateral gene transfer and the nature of bacterial innovation. Nature 405, 299–304 (2000).

21. Tria, F. D. K. & Martin, W. F. Gene Duplications Are At Least 50 Times Less Frequent than Gene Transfers in Prokaryotic Genomes. Genome Biology and Evolution 13, evab224 (2021).

22. Tawfik, O. K. & S, D. Enzyme Promiscuity: A Mechanistic and Evolutionary Perspective. Annual Review of Biochemistry 79, 471–505 (2010).

23. Weng, J.-K. & Noel, J. P. The Remarkable Pliability and Promiscuity of Specialized Metabolism. Cold Spring Harb Symp Quant Biol 77, 309–320 (2012).

24. Weng, J.-K. The evolutionary paths towards complexity: a metabolic perspective. New Phytologist 201, 1141–1149 (2014).

25. Lou, Y.-R., Pichersky, E. & Last, R. L. Deep roots and many branches: Origins of plant-specialized metabolic enzymes in general metabolism. Current Opinion in Plant Biology 66, 102192 (2022).

26. Maeda, H., Yoo, H. & Dudareva, N. Prephenate aminotransferase directs plant phenylalanine biosynthesis via arogenate. Nat Chem Biol 7, 19–21 (2011).

27. Koper, K., Han, S.-W., Pastor, D. C., Yoshikuni, Y. & Maeda, H. A. Evolutionary Origin and Functional Diversification of Aminotransferases. Journal of Biological Chemistry 102122 (2022) doi:10.1016/j.jbc.2022.102122.

28. Mehta, P. K., Hale, T. I. & Christen, P. Aminotransferases: demonstration of homology and division into evolutionary subgroups. European Journal of Biochemistry 214, 549–561 (1993).

29. Qian, K. et al. Hepatic ALT isoenzymes are elevated in gluconeogenic conditions including diabetes and suppressed by insulin at the protein level. Diabetes Metab Res Rev 31, 562–571 (2015).

30. Jansen, R. S. et al. Aspartate aminotransferase Rv3722c governs aspartate-dependent nitrogen metabolism in Mycobacterium tuberculosis. Nat Commun 11, 1960 (2020).

31. Ling, Z.-N. et al. Amino acid metabolism in health and disease. Signal Transduct Target Ther 8, 345 (2023).

32. Schulz-Mirbach, H. et al. On the flexibility of the cellular amination network in E coli. Elife 11, e77492 (2022).

33. Shrawat, A. K., Carroll, R. T., DePauw, M., Taylor, G. J. & Good, A. G. Genetic engineering of improved nitrogen use efficiency in rice by the tissue-specific expression of alanine aminotransferase. Plant Biotechnol J 6, 722–732 (2008).

34. Toney, M. D. Aspartate aminotransferase: An old dog teaches new tricks. Archives of Biochemistry and Biophysics 544, 119–127 (2014).

35. Agapova, A. et al. Flexible nitrogen utilisation by the metabolic generalist pathogen Mycobacterium tuberculosis. Elife 8, e41129 (2019).

36. Koonin, E. V., Mushegian, A. R. & Bork, P. Non-orthologous gene displacement. Trends Genet 12, 334–336 (1996).

37. Moghe, G. D. & Last, R. L. Something Old, Something New: Conserved Enzymes and the Evolution of Novelty in Plant Specialized Metabolism1. Plant Physiol 169, 1512–1523 (2015).

38. Kusano, H. et al. Evolutionary Developments in Plant Specialized Metabolism, Exemplified by Two Transferase Families. Front Plant Sci 10, 794 (2019).

39. Weng, J.-K., Philippe, R. N. & Noel, J. P. The rise of chemodiversity in plants. Science 336, 1667– 1670 (2012).

40. Mizutani, M. & Ohta, D. Diversification of P450 genes during land plant evolution. Annu Rev Plant Biol 61, 291–315 (2010).

41. Jia, Q. et al. Origin and early evolution of the plant terpene synthase family. Proc Natl Acad Sci U S A 119, e2100361119 (2022).

42. Torres, J. P. & Schmidt, E. W. The biosynthetic diversity of the animal world. J Biol Chem 294, 17684–17692 (2019).

43. Zhu, Q. et al. Phylogenomics of 10,575 genomes reveals evolutionary proximity between domains Bacteria and Archaea. Nat Commun 10, 5477 (2019).

44. Cavalier-Smith, T. & Chao, E. E.-Y. Multidomain ribosomal protein trees and the planctobacterial origin of neomura (eukaryotes, archaebacteria). Protoplasma 257, 621–753 (2020).

45. Pires, N. & Dolan, L. Early evolution of bHLH proteins in plants. Plant Signaling & Behavior 5, 911–912 (2010).

46. Goodstein, D. M. et al. Phytozome: a comparative platform for green plant genomics. Nucleic Acids Research 40, D1178–D1186 (2012).

47. Dunn, C. W., Giribet, G., Edgecombe, G. D. & Hejnol, A. Animal Phylogeny and Its Evolutionary Implications. Annual Review of Ecology, Evolution, and Systematics 45, 371–395 (2014).

48. Grigoriev, I. V. et al. MycoCosm portal: gearing up for 1000 fungal genomes. Nucleic Acids Res 42, D699–D704 (2014).

49. Burki, F., Roger, A. J., Brown, M. W. & Simpson, A. G. B. The New Tree of Eukaryotes. Trends in Ecology & Evolution 35, 43–55 (2020).

50. Mistry, J. et al. Pfam: The protein families database in 2021. Nucleic Acids Research 49, D412– D419 (2021).

51. Illergård, K., Ardell, D. H. & Elofsson, A. Structure is three to ten times more conserved than sequence--a study of structural response in protein cores. Proteins 77, 499–508 (2009).

52. Holm, L. & Sander, C. Mapping the protein universe. Science 273, 595–603 (1996).

53. Rozewicki, J., Li, S., Amada, K. M., Standley, D. M. & Katoh, K. MAFFT-DASH: integrated protein sequence and structural alignment. Nucleic Acids Res 47, W5–W10 (2019).

54. Finn, R. D. et al. Pfam: clans, web tools and services. Nucleic Acids Research 34, D247–D251 (2006).

55. Price, M. N., Dehal, P. S. & Arkin, A. P. FastTree 2 – Approximately Maximum-Likelihood Trees for Large Alignments. PLOS ONE 5, e9490 (2010).

56. Breazeale, S. D., Ribeiro, A. A. & Raetz, C. R. H. Origin of Lipid A Species Modified with 4-Amino-4-deoxy-l-arabinose in Polymyxin-resistant Mutants of Escherichia coli: AN AMINOTRANSFERASE (ArnB) THAT GENERATES UDP-4-AMINO-4-DEOXY-l-ARABINOSE*. Journal of Biological Chemistry 278, 24731–24739 (2003).

57. Noland, B. W. et al. Structural studies of Salmonella typhimurium ArnB (PmrH) aminotransferase: a 4-amino-4-deoxy-L-arabinose lipopolysaccharide-modifying enzyme. Structure 10, 1569–1580 (2002).

58. Amann, R. et al. Obligate intracellular bacterial parasites of acanthamoebae related to Chlamydia spp. Appl Environ Microbiol 63, 115–121 (1997).

59. Baker, B. J. et al. Enigmatic, ultrasmall, uncultivated Archaea. Proc Natl Acad Sci U S A 107, 8806–8811 (2010).

60. Heimerl, T. et al. A Complex Endomembrane System in the Archaeon Ignicoccus hospitalis Tapped by Nanoarchaeum equitans. Front Microbiol 8, 1072 (2017).

61. Huang, X. et al. Ancestral Genomes: a resource for reconstructed ancestral genes and genomes across the tree of life. Nucleic Acids Res 47, D271–D279 (2019).

62. Von Wettstein, D., Gough, S. & Kannangara, C. Chlorophyll Biosynthesis. Plant Cell 7, 1039– 1057 (1995).

63. Lu, M. et al. Role of the malate–aspartate shuttle on the metabolic response to myocardial ischemia. Journal of Theoretical Biology 254, 466–475 (2008).

64. Han, Q., Robinson, H., Cai, T., Tagle, D. A. & Li, J. Biochemical and structural characterization of mouse mitochondrial aspartate aminotransferase, a newly identified kynurenine aminotransferase-IV. Biosci Rep 31, 323–332 (2011).

65. Yang, R.-Z., Blaileanu, G., Hansen, B. C., Shuldiner, A. R. & Gong, D.-W. cDNA cloning, genomic structure, chromosomal mapping, and functional expression of a novel human alanine aminotransferase. Genomics 79, 445–450 (2002).

66. Stepanova, A. N. et al. TAA1-Mediated Auxin Biosynthesis Is Essential for Hormone Crosstalk and Plant Development. Cell 133, 177–191 (2008).

67. de Raad, M. et al. Mass spectrometry imaging-based assays for aminotransferase activity reveal a broad substrate spectrum for a previously uncharacterized enzyme. Journal of Biological Chemistry 102939 (2023) doi:10.1016/j.jbc.2023.102939.

68. Rebeaud, M. E., Mallik, S., Goloubinoff, P. & Tawfik, D. S. On the evolution of chaperones and cochaperones and the expansion of proteomes across the Tree of Life. Proc Natl Acad Sci U S A 118, e2020885118 (2021).

69. Zhou, X., Shen, X.-X., Hittinger, C. T. & Rokas, A. Evaluating Fast Maximum Likelihood-Based Phylogenetic Programs Using Empirical Phylogenomic Data Sets. Molecular Biology and Evolution 35, 486–503 (2018).

70. Son, H. F. & Kim, K.-J. Structural Insights into a Novel Class of Aspartate Aminotransferase from Corynebacterium glutamicum. PLoS One 11, e0158402 (2016).

71. Carrillo-Carrasco, V. P., Hernandez-Garcia, J., Mutte, S. K. & Weijers, D. The birth of a giant: evolutionary insights into the origin of auxin responses in plants. EMBO J 42, e113018 (2023).

72. Wu, G. Amino acids: metabolism, functions, and nutrition. Amino Acids 37, 1–17 (2009).

73. Davis, G. R. F. Essential Dietary Amino Acids for Growth of Larvae of the Yellow Mealworm, Tenebrio molitor L. The Journal of Nutrition 105, 1071–1075 (1975).

74. Fitzgerald, L. M. & Szmant, A. M. Biosynthesis of ‘essential’ amino acids by scleractinian corals. Biochem J 322, 213–221 (1997).

75. King, N. et al. The genome of the choanoflagellate Monosiga brevicollis and the origin of metazoans. Nature 451, 783–788 (2008).

76. Ilag, L. L., Kumar, A. M. & Söll, D. Light regulation of chlorophyll biosynthesis at the level of 5-aminolevulinate formation in Arabidopsis. Plant Cell 6, 265–275 (1994).

77. Zheng, Z. et al. Coordination of auxin and ethylene biosynthesis by the aminotransferase VAS1. Nat Chem Biol 9, 244–246 (2013).

78. Shih, P. M. et al. Biochemical characterization of predicted Precambrian RuBisCO. Nat Commun 7, 10382 (2016).

79. Hsiao, C., Mohan, S., Kalahar, B. K. & Williams, L. D. Peeling the Onion: Ribosomes Are Ancient Molecular Fossils. Molecular Biology and Evolution 26, 2415–2425 (2009).

80. Alva, V., Ammelburg, M., Söding, J. & Lupas, A. N. On the origin of the histone fold. BMC Struct Biol 7, 17 (2007).

81. Cerutti, H. & Casas-Mollano, J. A. On the origin and functions of RNA-mediated silencing: from protists to man. Curr Genet 50, 81–99 (2006).

82. Iwasaki, T. et al. Escherichia coli amino acid auxotrophic expression host strains for investigating protein structure–function relationships. The Journal of Biochemistry 169, 387–394 (2021).

83. Urrestarazu, A., Vissers, S., Iraqui, I. & Grenson, M. Phenylalanine- and tyrosine-auxotrophic mutants of Saccharomyces cerevisiae impaired in transamination. Mol Gen Genet 257, 230–237 (1998).

84. Martínez-Carranza, E. et al. Variability of Bacterial Essential Genes Among Closely Related Bacteria: The Case of Escherichia coli. Front Microbiol 9, 1059 (2018).

85. Charlebois, R. L. & Doolittle, W. F. Computing prokaryotic gene ubiquity: rescuing the core from extinction. Genome Res 14, 2469–2477 (2004).

86. Pieck, M. et al. Auxin and Tryptophan Homeostasis Are Facilitated by the ISS1/VAS1 Aromatic Aminotransferase in Arabidopsis. Genetics 201, 185–199 (2015).

87. Mavrides, C. & Orr, W. Multispecific aspartate and aromatic amino acid aminotransferases in Escherichia coli. Journal of Biological Chemistry 250, 4128–4133 (1975).

88. Nasir, N., Anant, A., Vyas, R. & Biswal, B. K. Crystal structures of Mycobacterium tuberculosis HspAT and ArAT reveal structural basis of their distinct substrate specificities. Sci Rep 6, 18880 (2016).

89. Wang, M., Toda, K. & Maeda, H. A. Biochemical properties and subcellular localization of tyrosine aminotransferases in Arabidopsis thaliana. Phytochemistry 132, 16–25 (2016).

90. Han, Q., Cai, T., Tagle, D. A. & Li, J. Structure, expression, and function of kynurenine aminotransferases in human and rodent brains. Cell Mol Life Sci 67, 353–368 (2010).

91. Dornfeld, C. et al. Phylobiochemical Characterization of Class-Ib Aspartate/Prephenate Aminotransferases Reveals Evolution of the Plant Arogenate Phenylalanine Pathway. The Plant Cell 26, 3101–3114 (2014).

92. Kim, S. H., Schneider, B. L. & Reitzer, L. Genetics and regulation of the major enzymes of alanine synthesis in Escherichia coli. J. Bacteriol. 192, 5304–5311 (2010).

93. Mikkelsen, M. D., Naur, P. & Halkier, B. A. Arabidopsis mutants in the C-S lyase of glucosinolate biosynthesis establish a critical role for indole-3-acetaldoxime in auxin homeostasis. Plant J. 37, 770–777 (2004).

94. Wang, M. & Maeda, H. A. Aromatic amino acid aminotransferases in plants. Phytochem Rev 17, 131–159 (2018).

95. Han, Q., Li, J. & Li, J. pH dependence, substrate specificity and inhibition of human kynurenine aminotransferase I. European Journal of Biochemistry 271, 4804–4814 (2004).

96. Rossi, F., Garavaglia, S., Montalbano, V., Walsh, M. A. & Rizzi, M. Crystal Structure of Human Kynurenine Aminotransferase II, a Drug Target for the Treatment of Schizophrenia*. Journal of Biological Chemistry 283, 3559–3566 (2008).

97. Graber, R. et al. Changing the Reaction Specificity of a Pyridoxal-5′-phosphate-dependent Enzyme. European Journal of Biochemistry 232, 686–690 (1995).

98. Palanivelu, R., Brass, L., Edlund, A. F. & Preuss, D. Pollen tube growth and guidance is regulated by POP2, an Arabidopsis gene that controls GABA levels. Cell 114, 47–59 (2003).

99. Stoner, G. L. & Eisenberg, M. A. Purification and properties of 7, 8-diaminopelargonic acid aminotransferase. J Biol Chem 250, 4029–4036 (1975).

100. Christen, P. & Mehta, P. K. From cofactor to enzymes. The molecular evolution of pyridoxal-5′-phosphate-dependent enzymes. The Chemical Record 1, 436–447 (2001).

101. Dolzan, M. et al. Crystal structure and reactivity of YbdL from Escherichia coli identify a methionine aminotransferase function. FEBS Letters 571, 141–146 (2004).

102. Fukumoto, Y. et al. Structural and functional role of the amino-terminal region of porcine cytosolic aspartate aminotransferase: Catalytic and structural properties of enzyme derivatives truncated on the amino-terminal side *. Journal of Biological Chemistry 266, 4187–4193 (1991).

103. Rossi, F., Han, Q., Li, J., Li, J. & Rizzi, M. Crystal structure of human kynurenine aminotransferase I. J Biol Chem 279, 50214–50220 (2004).

104. Han, Q., Robinson, H., Cai, T., Tagle, D. A. & Li, J. Structural Insight into the Inhibition of Human Kynurenine Aminotransferase I/Glutamine transaminase K. J Med Chem 52, 2786–2793 (2009).

105. Clark, J. W. & Donoghue, P. C. J. Whole-Genome Duplication and Plant Macroevolution. Trends in Plant Science 23, 933–945 (2018).

106. Albertin, W. & Marullo, P. Polyploidy in fungi: evolution after whole-genome duplication. Proceedings of the Royal Society B: Biological Sciences 279, 2497–2509 (2012).

107. Ahrens, J. B., Nunez-Castilla, J. & Siltberg-Liberles, J. Evolution of intrinsic disorder in eukaryotic proteins. Cell. Mol. Life Sci. 74, 3163–3174 (2017).

108. Holland, L. Z. & Ocampo Daza, D. A new look at an old question: when did the second whole genome duplication occur in vertebrate evolution? Genome Biology 19, 209 (2018).

109. Li, Z. et al. Multiple large-scale gene and genome duplications during the evolution of hexapods. Proceedings of the National Academy of Sciences 115, 4713–4718 (2018).

110. Dadras, A. et al. Accessible versatility underpins the deep evolution of plant specialized metabolism. Phytochem Rev (2023) doi:10.1007/s11101-023-09863-2.

111. Maeda, H. A. & Fernie, A. R. Evolutionary History of Plant Metabolism. Annu Rev Plant Biol 72, 185–216 (2021).

112. Nowack, E. C. M. et al. Gene transfers from diverse bacteria compensate for reductive genome evolution in the chromatophore of Paulinella chromatophora. Proc Natl Acad Sci U S A 113, 12214–12219 (2016).

113. de Crécy-Lagard, V. Variations in metabolic pathways create challenges for automated metabolic reconstructions: Examples from the tetrahydrofolate synthesis pathway. Comput Struct Biotechnol J 10, 41–50 (2014).

114. Denise, R., Babor, J., Gerlt, J. A. & de Crécy-Lagard, V. Pyridoxal 5’-phosphate synthesis and salvage in Bacteria and Archaea: predicting pathway variant distributions and holes. Microb Genom 9, mgen000926 (2023).

115. Dembech, E. et al. Identification of hidden associations among eukaryotic genes through statistical analysis of coevolutionary transitions. Proc Natl Acad Sci U S A 120, e2218329120 (2023).

116. Hirotsu, K., Goto, M., Okamoto, A. & Miyahara, I. Dual substrate recognition of aminotransferases. The Chemical Record 5, 160–172 (2005).

117. Koper, K., Hataya, S., Hall, A. G., Takasuka, T. E. & Maeda, H. A. Biochemical characterization of plant aromatic aminotransferases. in Methods in Enzymology (Academic Press, 2022). doi:10.1016/bs.mie.2022.07.034.

118. Dietrich, M. R., Ankeny, R. A. & Chen, P. M. Publication Trends in Model Organism Research. Genetics 198, 787–794 (2014).

119. The UniProt Consortium. UniProt: the universal protein knowledgebase in 2021. Nucleic Acids Research 49, D480–D489 (2021).

120. Finn, R. D., Clements, J. & Eddy, S. R. HMMER web server: interactive sequence similarity searching. Nucleic Acids Research 39, W29–W37 (2011).

121. Shimodaira, H. & Hasegawa, M. Multiple Comparisons of Log-Likelihoods with Applications to Phylogenetic Inference. Molecular Biology and Evolution 16, 1114 (1999).

122. Huerta-Cepas, J., Serra, F. & Bork, P. ETE 3: Reconstruction, Analysis, and Visualization of Phylogenomic Data. Molecular Biology and Evolution 33, 1635–1638 (2016).

123. Chang, A. et al. BRENDA, the ELIXIR core data resource in 2021: new developments and updates. Nucleic Acids Research 49, D498–D508 (2021).

124. Wittig, U., Rey, M., Weidemann, A., Kania, R. & Müller, W. SABIO-RK: an updated resource for manually curated biochemical reaction kinetics. Nucleic Acids Research 46, D656–D660 (2018).

125. Waskom, M. L. seaborn: statistical data visualization. Journal of Open Source Software 6, 3021 (2021).

126. Armenteros, J. J. A. et al. Detecting sequence signals in targeting peptides using deep learning. Life Science Alliance 2, (2019).

127. Burley, S. K. et al. RCSB Protein Data Bank: powerful new tools for exploring 3D structures of biological macromolecules for basic and applied research and education in fundamental biology, biomedicine, biotechnology, bioengineering and energy sciences. Nucleic Acids Research 49, D437–D451 (2021).

128. Jumper, J. et al. Highly accurate protein structure prediction with AlphaFold. Nature 596, 583– 589 (2021).

129. Evans, R. et al. Protein complex prediction with AlphaFold-Multimer. 2021.10.04.463034 Preprint at 10.1101/2021.10.04.463034 (2022).

130. Waterhouse, A. et al. SWISS-MODEL: homology modelling of protein structures and complexes. Nucleic Acids Research 46, W296–W303 (2018).

131. Webb, B. & Sali, A. Comparative Protein Structure Modeling Using MODELLER. Current Protocols in Bioinformatics 54, 5.6.1–5.6.37 (2016).

132. Crooks, G. E., Hon, G., Chandonia, J.-M. & Brenner, S. E. WebLogo: A Sequence Logo Generator. Genome Res. 14, 1188–1190 (2004).

133. Morris, G. M. et al. AutoDock4 and AutoDockTools4: Automated Docking with Selective Receptor Flexibility. J Comput Chem 30, 2785–2791 (2009).

134. Pettersen, E. F. et al. UCSF Chimera—A visualization system for exploratory research and analysis. Journal of Computational Chemistry 25, 1605–1612 (2004).

135. Meng, E. C., Pettersen, E. F., Couch, G. S., Huang, C. C. & Ferrin, T. E. Tools for integrated sequence-structure analysis with UCSF Chimera. BMC Bioinformatics 7, 339 (2006).

## References for Supplementary Figures

1. Zhu, Q. et al. Phylogenomics of 10,575 genomes reveals evolutionary proximity between domains Bacteria and Archaea. Nat Commun 10, 5477 (2019).

2. Cavalier-Smith, T. & Chao, E. E.-Y. Multidomain ribosomal protein trees and the planctobacterial origin of neomura (eukaryotes, archaebacteria). Protoplasma 257, 621–753 (2020).

3. Burki, F., Roger, A. J., Brown, M. W. & Simpson, A. G. B. The New Tree of Eukaryotes. Trends in Ecology & Evolution 35, 43–55 (2020).

4. Pires, N. & Dolan, L. Early evolution of bHLH proteins in plants. Plant Signaling & Behavior 5, 911– 912 (2010).

5. Dunn, C. W., Giribet, G., Edgecombe, G. D. & Hejnol, A. Animal Phylogeny and Its Evolutionary Implications. Annual Review of Ecology, Evolution, and Systematics 45, 371–395 (2014).

6. Grigoriev, I. V. et al. MycoCosm portal: gearing up for 1000 fungal genomes. Nucleic Acids Res 42, D699–D704 (2014).

7. Morris, G. M. et al. AutoDock4 and AutoDockTools4: Automated Docking with Selective Receptor Flexibility. J Comput Chem 30, 2785–2791 (2009).

8. Fukumoto, Y. et al. Structural and functional role of the amino-terminal region of porcine cytosolic aspartate aminotransferase: Catalytic and structural properties of enzyme derivatives truncated on the amino-terminal side *. Journal of Biological Chemistry 266, 4187–4193 (1991).

9. Rossi, F., Han, Q., Li, J., Li, J. & Rizzi, M. Crystal structure of human kynurenine aminotransferase I. J Biol Chem 279, 50214–50220 (2004).

10. Han, Q., Robinson, H., Cai, T., Tagle, D. A. & Li, J. Structural Insight into the Inhibition of Human Kynurenine Aminotransferase I/Glutamine transaminase K. J Med Chem 52, 2786–2793 (2009).

