## Supplementary material for "Multi-substrate specificity shaped the complex evolution of the aminotransferase family across the tree of life": Dataset S4

### Dataset S4: ToL-scale phylogenetic trees of AT and related enzymes with sequence IDs.

#### Class I ATs

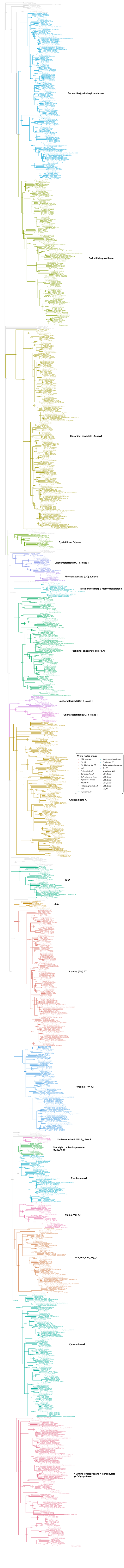

### Class II ATs

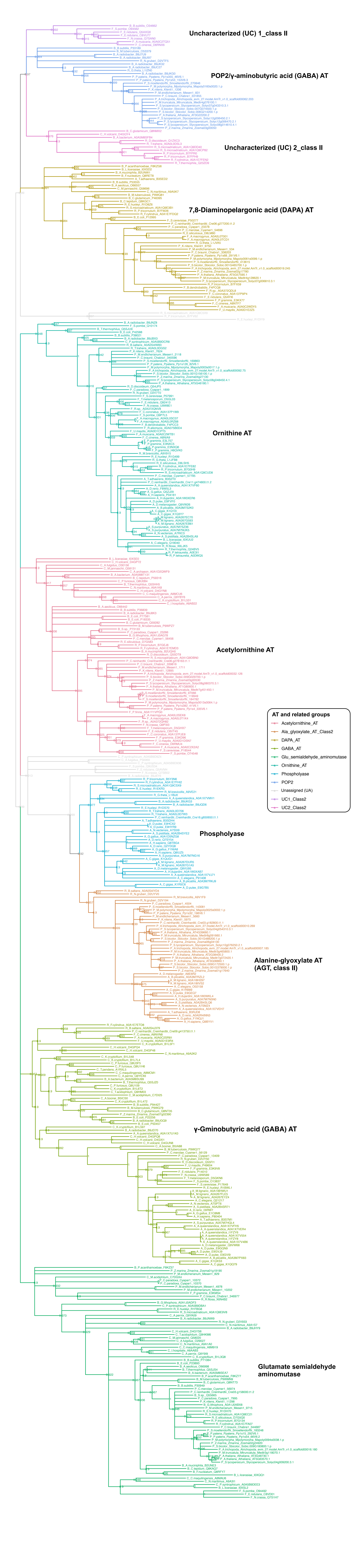
Uncharacterized (UC) 1_class II
POP2/γ-aminobutyric acid (GABA) AT
Uncharacterized (UC) 2_class II
7,8-Diaminopelargonic acid (DAPA) AT
Ornithine AT
Acetylornithine AT
Phospholyase
Alanine-glyoxylate AT (AGT, class II)
γ-Gminobutyric acid (GABA) AT
Glutamate semialdehyde aminomutase
AT and related groups
Acetylornithine_AT
Ala_glyoxylate_AT_Class2
DAPA_AT
GABA_AT
Glu_semialdehyde_aminomutase
Ornithine_AT
Phospholyase
POP2
Unassigned (UA)
UC1_Class2
UC2_Class2

### Class III ATs

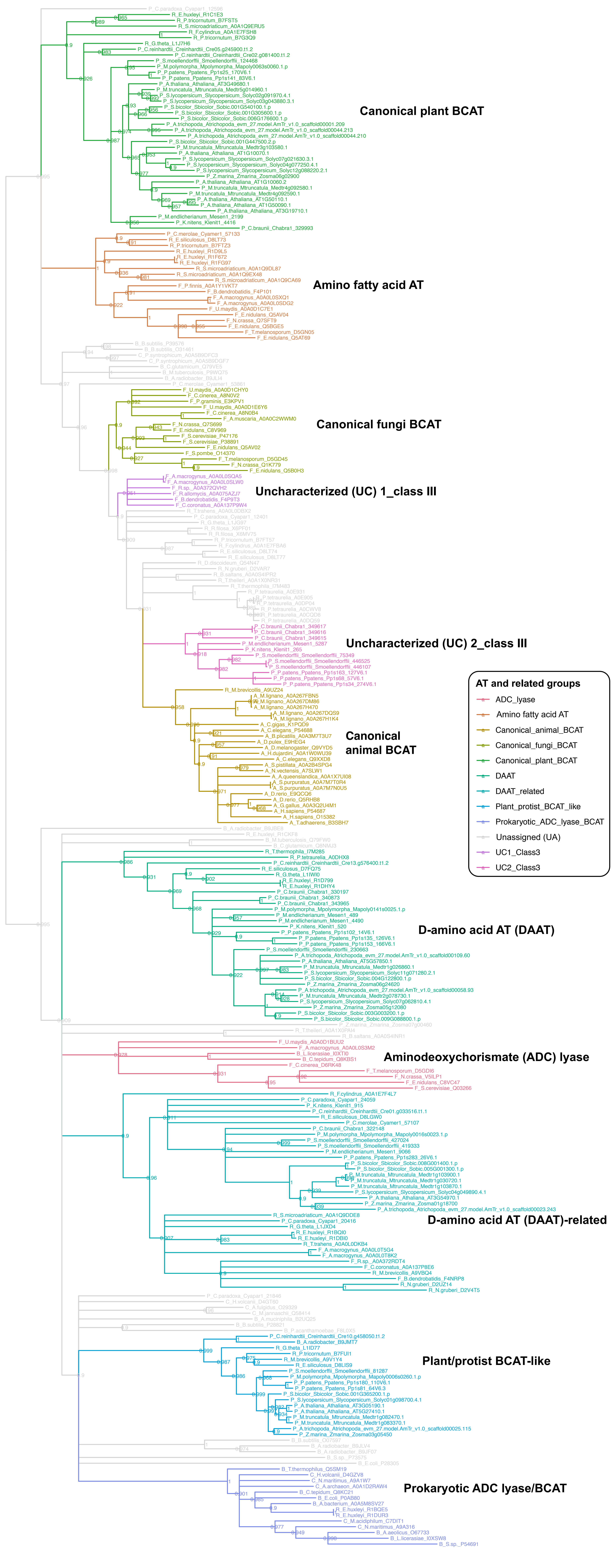

### Class IV

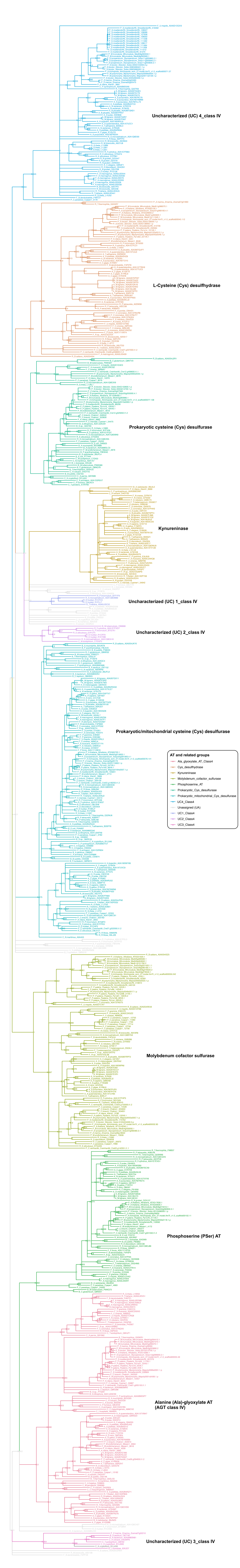

Uncharacterized (UC) 4_class IV
L-Cysteine (Cys) desulfhydrase
Prokaryotic cysteine (Cys) desulfurase
Kynureninase
Uncharacterized (UC) 1_class IV
Uncharacterized (UC) 2_class IV
Prokaryotic/mitochondrial cysteine (Cys) desulfurase
AT and related groups
Ala_glyoxylate_AT_Class4
Cys_desulfhydrase
Kynureninase
Molybdenum_cofactor_sulfurase
Phosphoserine_AT
Prokaryotic_Cys_desulfurase
Prokaryotic_mitochondrial_Cys_desulfurase
UC4_Class4
Unassigned (UA)
UC1_Class4
UC2_Class4
UC3_Class4
Molybdenum cofactor sulfurase
Phosphoserine (PSer) AT
Alanine (Ala)-glyoxylate AT
(AGT class IV)
Uncharacterized (UC) 3_class IV

### Alliinase

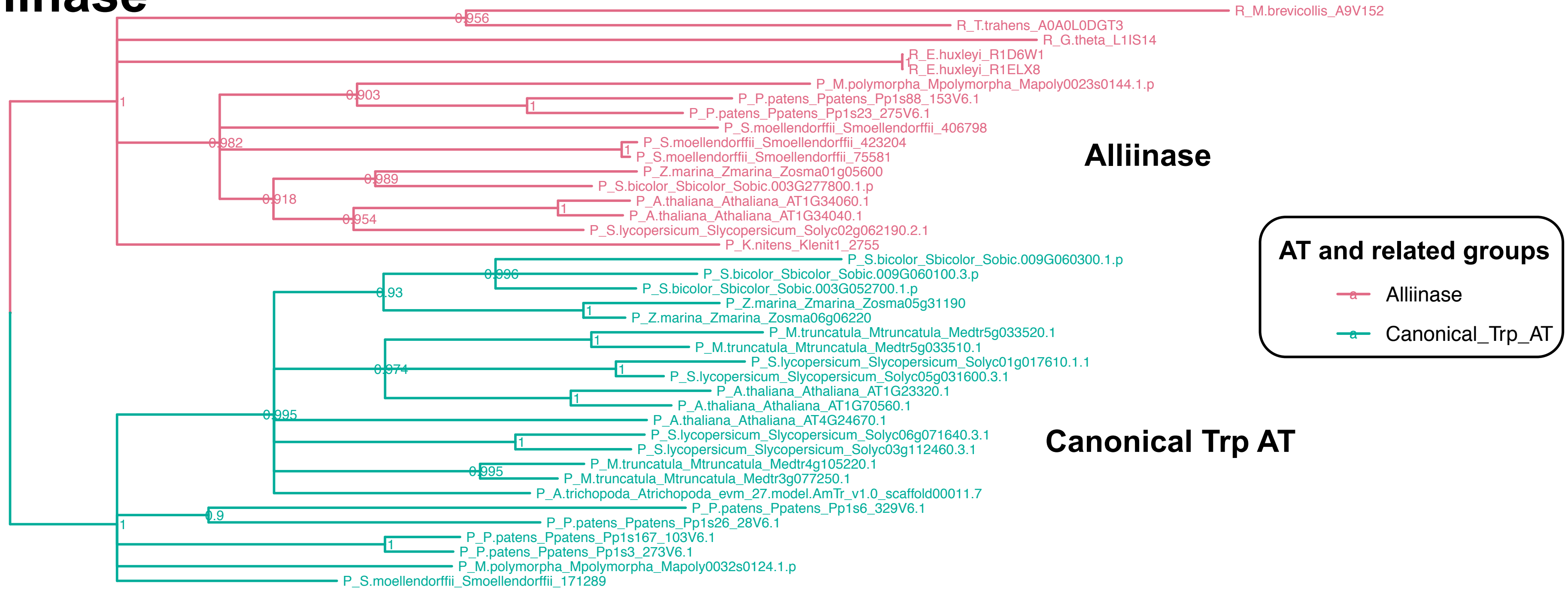

R_M.brevicollis_A9V152
R_T.trahens_A0A0L0DGT3
R_G.theta_L1IS14
R_E.huxleyi_R1D6W1
R_E.huxleyi_R1ELX8
P_M.polymorpha_Mpolymorpha_Mapoly0023s0144.1.p
P_P.patens_Ppatens_Pp1s88_153V6.1
P_P.patens_Ppatens_Pp1s23_275V6.1
P_S.moellendorffii_Smoellendorffii_406798
P_S.moellendorffii_Smoellendorffii_423204
P_S.moellendorffii_Smoellendorffii_75581
P_Z.marina_Zmarina_Zosma01g05600
P_S.bicolor_Sbicolor_Sobic.003G277800.1.p
P_A.thaliana_Athaliana_AT1G34060.1
P_A.thaliana_Athaliana_AT1G34040.1
P_S.lycopersicum_Slycopersicum_Solyc02g062190.2.1
P_K.nitens_Klenit1_2755
P_S.bicolor_Sbicolor_Sobic.009G060300.1.p
P_S.bicolor_Sbicolor_Sobic.009G060100.3.p
P_S.bicolor_Sbicolor_Sobic.003G052700.1.p
P_Z.marina_Zmarina_Zosma05g31190
P_Z.marina_Zmarina_Zosma06g06220
P_M.truncatula_Mtruncatula_Medtr5g033520.1
P_M.truncatula_Mtruncatula_Medtr5g033510.1
P_S.lycopersicum_Slycopersicum_Solyc01g017610.1.1
P_S.lycopersicum_Slycopersicum_Solyc05g031600.3.1
P_A.thaliana_Athaliana_AT1G23320.1
P_A.thaliana_Athaliana_AT1G70560.1
P_A.thaliana_Athaliana_AT4G24670.1
P_S.lycopersicum_Slycopersicum_Solyc06g071640.3.1
P_S.lycopersicum_Slycopersicum_Solyc03g112460.3.1
P_M.truncatula_Mtruncatula_Medtr4g105220.1
P_M.truncatula_Mtruncatula_Medtr3g077250.1
P_A.trichopoda_Atrichopoda_evm_27.model.AmTr_v1.0_scaffold00011.7
P_P.patens_Ppatens_Pp1s6_329V6.1
P_P.patens_Ppatens_Pp1s26_28V6.1
P_P.patens_Ppatens_Pp1s167_103V6.1
P_P.patens_Ppatens_Pp1s3_273V6.1
P_M.polymorpha_Mpolymorpha_Mapoly0032s0124.1.p
P_S.moellendorffii_Smoellendorffii_171289
0.956
0.903
0.982
0.989
0.918
0.954
0.996
0.93
0.974
0.995
0.9
Alliinase
Canonical Trp AT
AT and related groups
Alliinase
Canonical_Trp_AT

### Sugar ATs

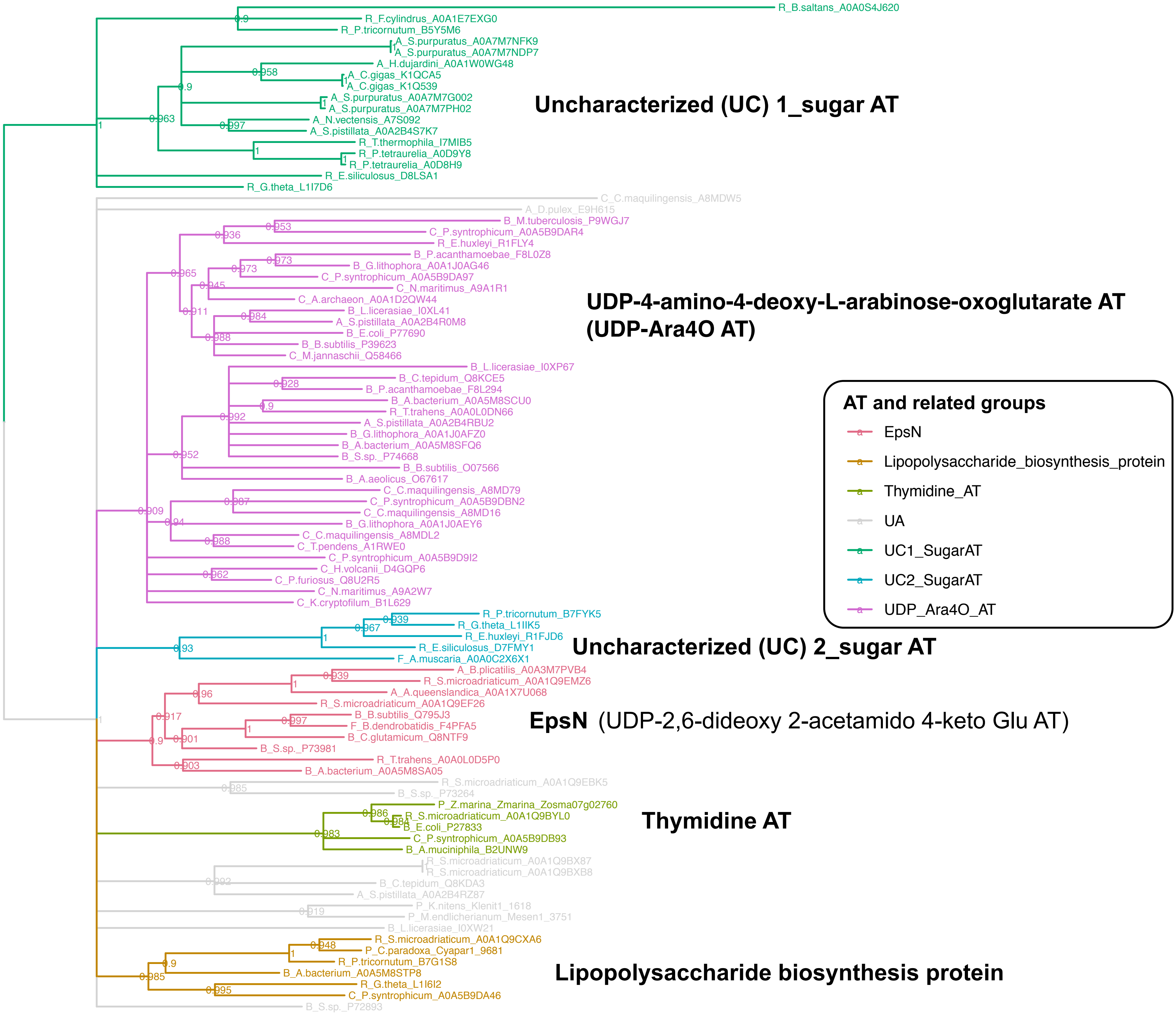

R_B.saltans_A0A0S4J620
R_F.cylindrus_A0A1E7EXG0
R_P.tricornutum_B5Y5M6
A_S.purpuratus_A0A7M7NFK9
A_S.purpuratus_A0A7M7NDP7
A_H.dujardini_A0A1W0WG48
A_C.gigas_K1QCA5
A_C.gigas_K1Q539
A_S.purpuratus_A0A7M7G002
A_S.purpuratus_A0A7M7PH02
A_N.vectensis_A7S092
A_S.pistillata_A0A2B4S7K7
R_T.thermophila_I7MIB5
R_P.tetraurelia_A0D9Y8
R_P.tetraurelia_A0D8H9
R_E.siliculosus_D8LSA1
R_G.theta_L1I7D6
Uncharacterized (UC) 1_sugar AT
C_C.maquilingensis_A8MDW5
A_D.pulex_E9H615
B_M.tuberculosis_P9WGJ7
C_P.syntrophicum_A0A5B9DAR4
R_E.huxleyi_R1FLY4
B_P.acanthamoebae_F8L0Z8
B_G.lithophora_A0A1J0AG46
C_P.syntrophicum_A0A5B9DA97
C_N.maritimus_A9A1R1
C_A.archaeon_A0A1D2QW44
B_L.licerasiae_I0XL41
A_S.pistillata_A0A2B4R0M8
B_E.coli_P77690
B_B.subtilis_P39623
C_M.jannaschii_Q58466
UDP-4-amino-4-deoxy-L-arabinose-oxoglutarate AT
(UDP-Ara4O AT)
B_L.licerasiae_I0XP67
B_C.tepidum_Q8KCE5
B_P.acanthamoebae_F8L294
B_A.bacterium_A0A5M8SCU0
R_T.trahens_A0A0L0DN66
A_S.pistillata_A0A2B4RBU2
B_G.lithophora_A0A1J0AFZ0
B_A.bacterium_A0A5M8SFQ6
B_S.sp._P74668
B_B.subtilis_O07566
B_A.aeolicus_O67617
C_C.maquilingensis_A8MD79
C_P.syntrophicum_A0A5B9DBN2
C_C.maquilingensis_A8MD16
B_G.lithophora_A0A1J0AEY6
C_C.maquilingensis_A8MDL2
C_T.pendens_A1RWE0
C_P.syntrophicum_A0A5B9D9I2
C_H.volcanii_D4GQP6
C_P.furiosus_Q8U2R5
C_N.maritimus_A9A2W7
C_K.cryptofilum_B1L629
AT and related groups
EpsN
Lipopolysaccharide_biosynthesis_protein
Thymidine_AT
UA
UC1_SugarAT
UC2_SugarAT
UDP_Ara4O_AT
R_P.tricornutum_B7FYK5
R_G.theta_L1IIK5
R_E.huxleyi_R1FJD6
R_E.siliculosus_D7FMY1
F_A.muscaria_A0A0C2X6X1
Uncharacterized (UC) 2_sugar AT
A_B.plicatilis_A0A3M7PVB4
R_S.microadriaticum_A0A1Q9EMZ6
A_A.queenslandica_A0A1X7U068
R_S.microadriaticum_A0A1Q9EF26
B_B.subtilis_Q795J3
F_B.dendrobatidis_F4PFA5
B_C.glutamicum_Q8NTF9
B_S.sp._P73981
R_T.trahens_A0A0L0D5P0
B_A.bacterium_A0A5M8SA05
EpsN (UDP-2,6-dideoxy 2-acetamido 4-keto Glu AT)
R_S.microadriaticum_A0A1Q9EBK5
B_S.sp._P73264
P_Z.marina_Zmarina_Zosma07g02760
R_S.microadriaticum_A0A1Q9BYL0
B_E.coli_P27833
C_P.syntrophicum_A0A5B9DB93
B_A.muciniphila_B2UNW9
Thymidine AT
R_S.microadriaticum_A0A1Q9BX87
R_S.microadriaticum_A0A1Q9BXB8
B_C.tepidum_Q8KDA3
A_S.pistillata_A0A2B4RZ87
P_K.nitens_Klenit1_1618
P_M.endlicherianum_Mesen1_3751
B_L.licerasiae_I0XW21
R_S.microadriaticum_A0A1Q9CXA6
P_C.paradoxa_Cyapar1_9681
R_P.tricornutum_B7G1S8
B_A.bacterium_A0A5M8STP8
R_G.theta_L1I6I2
C_P.syntrophicum_A0A5B9DA46
B_S.sp._P72893
Lipopolysaccharide biosynthesis protein
